# A G protein-coupled receptor-like module regulates Cellulose Synthase secretion from the endomembrane system in Arabidopsis

**DOI:** 10.1101/2020.10.20.345868

**Authors:** Heather E. McFarlane, Daniela Mutwil-Anderwald, Jana Verbančič, Kelsey L. Picard, Timothy E. Gookin, Anja Froehlich, Luisa M. Trindade, Jose M. Alonso, Sarah M. Assmann, Staffan Persson

**Author notes:** **Corresponding Authors:** Staffan Persson (Lead Contact), School of Biosciences, University of Melbourne, Parkville, 3010 Victoria, Melbourne, Australia, Heather E. McFarlane, Department of Cell & Systems Biology, University of Toronto, 25 Harbord St, Toronto, ON, M5S 3G5, Canada.

## Abstract

Cellulose synthesis is essential for plant morphology, water transport and defense, and provides raw material for biomaterials and fuels. Cellulose is produced at the plasma membrane by Cellulose Synthase (CESA) protein complexes (CSCs). CSCs are assembled in the endomembrane system and then trafficked from the Golgi apparatus and *trans*-Golgi Network (TGN) to the plasma membrane. Since CESA enzymes are only active in the plasma membrane, control of CSC secretion is a critical step in the regulation of cellulose synthesis. However, the regulatory framework for CSC secretion is not clarified. In this study, we identify members of a family of seven transmembrane domain-containing proteins (7TMs) as important for cellulose production during cell wall integrity stress. 7TM proteins are often associated with guanine nucleotide-binding protein (G) protein signalling and mutants in several of the canonical G protein complex components phenocopied the *7tm* mutant plants. Unexpectedly, the 7TM proteins localized to the Golgi apparatus/TGN where they interacted with the G protein complex. Here, the 7TMs and G proteins regulated CESA trafficking, but did not affect general protein secretion. Furthermore, during cell wall stress, 7TMs’ localization was biased towards small CESA-containing vesicles, specifically associated with CSC trafficking. Our results thus outline how a G protein-coupled module regulates CESA trafficking and reveal that defects in this process lead to exacerbated responses upon exposure to cell wall integrity stress.

## Introduction

The plant cell wall is a polysaccharide-based cellular exoskeleton that provides the basis for directed plant growth and protects the cell against the environment. Cell wall polysaccharides are typically divided into three classes: pectins, hemicelluloses and cellulose. Cellulose, the main component of the primary cell wall, consists of β-1,4-linked glucans that coalesce into microfibrils through hydrogen bonding (McFarlane et al., 2014). These microfibrils are the load-bearing structures of the plant cell wall. Therefore, cellulose microfibril abundance and orientation primarily dictate the mechanical properties of most growing cell walls.

Cellulose is synthesized by Cellulose Synthase (CESA) protein complexes (CSCs) at the plasma membrane of plant cells (McFarlane et al., 2014). The CSCs consist of three structurally related CESA subunits. In Arabidopsis, CESA1, 3 and 6-like CESAs contribute to synthesis of the primary walls that surround all plant cells (Desprez et al., 2007; Persson et al., 2007). The CSCs are thought to be assembled in the Golgi apparatus, or perhaps the endoplasmic reticulum, and then secreted to the plasma membrane. Since CESA enzymes are only active in the plasma membrane, control of CSC secretion and endocytosis is a critical step in regulating cellulose synthesis. Several factors directly or indirectly contribute to CSC secretion, including the actin cytoskeleton (Sampathkumar et al., 2013; Zhang et al., 2019), the lumenal pH of endomembrane compartments (Luo et al., 2015), PATROL1 via the exocyst complex (Zhu et al., 2018), SHOU4 (Polko et al 2018) and the kinesin FRA1 (Zhu et al., 2015). In addition, two proteins called STELLO (STL) 1 and 2 regulate the secretion of the CSC, perhaps by aiding in the assembly of the complex in the Golgi apparatus (Zhang et al., 2016). Several small molecules also affect CSC trafficking. For example, isoxaben causes rapid internalization of the CSCs into Small CESA Containing Compartments (SmaCCs; Gutierrez et al., 2009; Crowell et al., 2009), 2,6-dichlorobenzonitrile (DCB) impacts CSC activity at the plasma membrane (Debolt et al., 2007), and cestrin affects trafficking of CSCs and associated proteins (Worden et al., 2015).

The delivery of CSCs to the plasma membrane typically occurs at sites marked by cortical microtubules (Gutierrez et al., 2009). Once in the plasma membrane, the CSCs become activated through an unknown mechanism and laterally diffuse in the plasma membrane to synthesize cellulose (Paredez et al., 2006). CSC movement is likely driven by glucose addition to the growing cellulose chain. As the chain is extruded from the enzyme it becomes entangled within the cell wall network, presumably causing the CSCs to move in the plasma membrane due to the addition of new glucose residues. The direction of this CSC movement may be steered by cortical microtubules (Paradez et al., 2006; Bringmann et al., 2012; Li et al., 2012). Once cellulose microfibrils reach a certain length, the CSCs stall and are internalized into the cell via clathrin-mediated endocytosis (Bashline et al., 2015; Sanchez-Rodriguez et al., 2018).

Plant cell walls are dynamic structures that undergo changes in response to environmental and developmental signals. It is therefore clear that, similar to yeast, plant cells need mechanisms to perceive and respond to changes in cell wall integrity (CWI) (Vaahtera et al., 2019; Rodicio and Heinisch, 2010). CWI changes can be induced by short term cell wall inhibitor treatments (e.g. with the cellulose synthesis inhibitor isoxaben), or via chronic alterations to cell wall synthesis or remodelling (e.g. over-expression of pectin modifying enzymes (Denness et al., 2011; Wolf et al., 2012)). Several receptor-like kinases (RLKs) have been implicated in CWI sensing, including members of the *Catharanthus roseus* RLK-like family, or CrRLKLs (Hematy et al., 2007; Haruta et al., 2014; Vaahtera et al., 2019). The CrRLKL THESEUS1 (THE1) negatively regulates growth when cellulose synthesis, and thus CWI, is impaired (Hematy et al., 2007). Consequently, cellulose deficient mutant plants display increased growth when THE1 function is compromised. Some of the CrRLKL members, including FERONIA and THE1, can bind to rapid alkalinisation factors (RALFs), which are peptides that promote apoplastic alkalinisation, potentially sensing cell wall changes and altering wall extensibility, which is controlled in part through pH-regulated expansins (Haruta et al., 2014; Gonneau et al., 2018; Feng et al., 2018). Several other RLK-based signal transduction circuits are associated with CWI sensing. For example, the brassinosteroid receptor BRASSINOSTEROID INSENSITIVE 1 (BRI1) perceives the pectin-status of the wall in association with the RECEPTOR-LIKE Protein 44 (RLP44) and the BRI1-ASSOCIATED KINASE1 (BAK1) (Wolf et al., 2012; Wolf et al., 2014). In addition, both FERONIA and the family of WALL ASSOCIATED KINASEs (WAKs) can bind to short stretches of the pectic polymer homogalacturonan, which presumably enables them to sense changes in the cell wall structure (Kohorn and Kohorn, 2012; Feng et al., 2018). Since CWI is likely sensed at the interface between the cell wall and the plasma membrane, the search for components that monitor cell wall damage has largely focused on plasma membrane-based proteins. While some downstream responses have been identified, e.g. local Ca^2+^ influx (Feng et al., 2018) or changes in Reactive Oxygen Species (ROS), that could influence cell wall flexibility via oxidative cell wall cross-linking (Raggi et al., 2015), mechanisms that explain how CWI impacts cell wall synthesis, secretion and remodeling are largely lacking.

Another component that has been linked to CWI signalling is the heterotrimeric guanine nucleotide-binding protein (G) protein complex (Klopffleisch et al., 2011; Delgado-Cerezo et al., 2012). This complex, comprised of Gα, Gβ and Gγ subunits, is conserved across eukaryotes, though there are substantial differences in number of subunits and their activities in plants and animals (Urano et al., 2016; Maruta et al., 2019). In the textbook model of G protein signalling, a plasma membrane-localized G protein-coupled receptor (GPCR) with seven transmembrane domains perceives an extracellular signal, resulting in Gα exchange of GDP for GTP. This exchange results in the dissociation of the Gαβγ heterotrimer so that GTP-bound Gα and the Gβγ dimer can elicit different intracellular signalling responses (Jones & Assmann, 2004). In the context of CWI signalling, components of the G protein signalling pathway have been linked to cell proliferation and cell expansion in plants (Ullah et al., 2001; Ullah et al., 2003; Chen et al., 2006; Jaffé et al., 2012; Choudhury et al., 2019), as well as pathogen-associated molecular pattern (PAMP)-triggered immunity (Sánchez-Rodríguez et al., 2009; Aranda-Sicilia et al., 2015; Tunc-Ozdemir et al., 2016; Liang et al., 2016; Escudero et al., 2017), all of which require close communication between the plant cell and its cell wall. Furthermore, a predicted interactome of G protein components in Arabidopsis revealed strong links to cell wall synthesis and modification genes (Klopffleisch et al., 2011). That study also demonstrated qualitative differences in the plant cell wall in several canonical G protein complex mutants. Similarly, Delgado-Cerezo et al. (2012) demonstrated that transcriptomic changes in *Gβ* and *Gγ* mutants are highly similar and are enriched in cell wall synthesis and modification genes. Together, these studies make components of the G protein complex, and their putative receptors, exciting candidates for CWI signalling components.

## Results

### Seven-transmembrane proteins are required for seedling growth during cell wall stress

Cellulose synthesis-associated genes tend to be co-expressed with the *CESA* genes (Persson et al., 2005; Brown et al., 2005). We employed the tool FamNet (Ruprecht et al., 2016) to re-assess genes and gene families (pfams) that are co-expressed with the primary cell wall *CESAs* (Supplemental Figure 1). We obtained T-DNA lines that disrupted several of the co-expressed genes and grew seedlings on MS media, or MS media supplemented with isoxaben, a potent cellulose synthesis inhibitor (Heim et al., 1990). Two T-DNA lines that targeted At5g18520, which corresponds to a putative seven-transmembrane domain (7TM) containing protein, displayed reduced hypocotyl growth, thicker hypocotyls on isoxaben-supplemented media and reduced transcript levels (*7tm1-1*) or no detectable transcript (*7tm1-2*) for the gene (Figure 1A & B; Supplemental Figure 2A). The mutants did not show any phenotypic deviations from wild type on media without isoxaben or under un-stressed growth conditions (Supplemental Figure 2).

**Figure 1.**
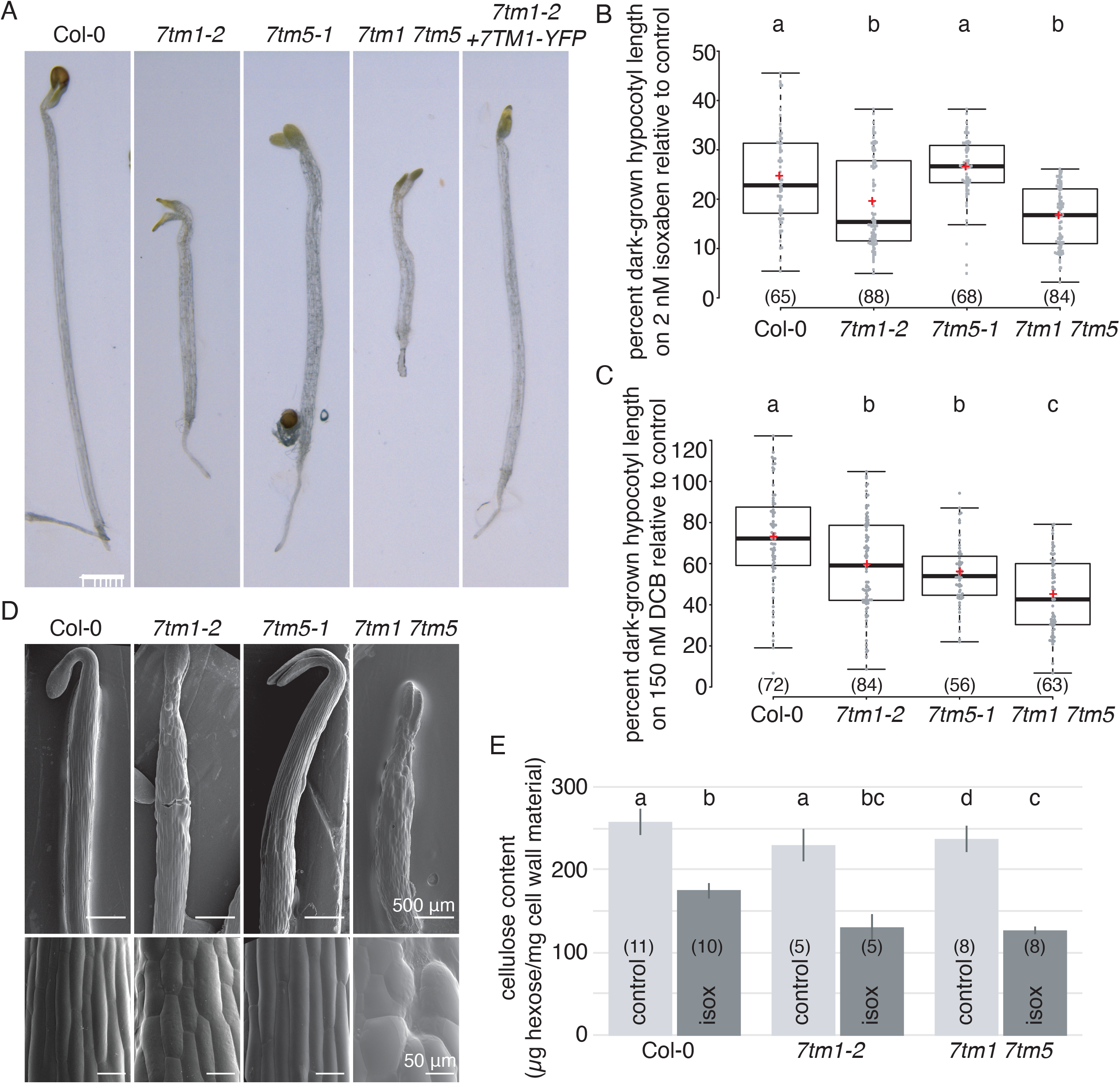
Mutations in 7TM family members cause increased sensitivity to cell wall stress. **A.** Representative images of seven-day-old etiolated seedlings grown on ½ MS media supplemented with 2 nM isoxaben. **B.** Quantification of etiolated hypocotyl lengths of seedlings represented in (A). **C.** Quantification of seven-day-old etiolated hypocotyl lengths of seedlings grown on ½ MS media supplemented with 150 nM DCB (see also Supplemental Figure 2). **D.** Cryo-scanning electron microscopy of three-day-old etiolated hypocotyls of seedlings grown on ½ MS media supplemented with 2 nM isoxaben. **E.** Cellulose content in different genotypes. All graphs summarize three independent experiments, n is indicated in parentheses; samples not sharing a common letter are significantly different (one-way ANOVA and Tukey HSD test, p<0.05), and in box plots, box limits indicate 25^th^ and 75^th^ percentiles, whiskers extend to 1.5 times the interquartile range, median is indicated by a line, mean by a red “+” and individual data points are shown. Scale bars represent 1 mm in A and 500 μm or 50 μm as indicated in B.

The 7TM protein is part of a small gene family in Arabidopsis (Supplemental Figure 2B) and we therefore refer to At5g18520 as 7TM1. We isolated T-DNA insertion mutants for each of the other closely related members of the 7TM clade, i.e. *7TM2* (At3g09570); *7TM3* (At5g02630) and *7TM4* (At5g42090). We also included T-DNA mutants for *7TM5* (At2g01070), which is in a second clade of 7TMs, as this gene was highly co-expressed with the *7TM1* gene (Supplemental Table 1). Of these mutants, only the *7tm1* seedlings displayed increased sensitivity to isoxaben as compared to wild type (Supplemental Figure 2C). To assess potential functional redundancy among the 7TM proteins, we generated double mutant combinations between *7tm1-2* and the other *7tm* mutants. Of the combinations we assayed, only *7tm1 7tm5* double mutant seedlings displayed an enhanced isoxaben sensitivity as compared to the *7tm1* seedlings (Figure 1A & B; Supplemental Figure 2C). The *7tm1* and *7tm1 7tm5* mutants also displayed increased sensitivity to the cellulose inhibitor DCB (Figure 1C; Supplemental Figure 3).

Cryo-scanning electron microscopy of etiolated seedlings grown on isoxaben-containing media revealed that the increased *7tm1* and *7tm1 7tm5* hypocotyl thickness was primarily due to epidermal cell swelling (Figure 1D), which is a common phenotype of cellulose synthesis deficiency (e.g. Arioli et al., 1998; Desprez et al., 2007; Persson et al., 2007). To further assess its role in cellulose synthesis, we introgressed *7tm1-2* into *procuste1-1* (*prc1-1*), a null mutation in *CESA6* (Fagard et al., 2000). The double homozygous progeny of *7tm1-2 prc1-1* displayed slightly decreased hypocotyl length as compared to the parent lines (Supplemental Figure 4). While disruption of cellulose synthesis thus resulted in growth defects of *7tm1* hypocotyls relative to wild type, other stress conditions, including sucrose, salinity, or oryzalin treatment, did not (Supplemental Figure 3). To assess whether the mutant seedlings were affected in their ability to produce cellulose, we measured cellulose content in etiolated seven-day old wild type and *7tm1* and *7tm1 7tm5* mutant seedlings. Under normal growth conditions, there was no significant difference in the level of cellulose between *7tm* mutants and wild type (Figure 1E). However, when grown on isoxaben, the *7tm1 7tm5* mutants had significantly less cellulose than wild type seedlings (Figure 1E). These results imply that the 7TMs play a role in cellulose production, particularly during cell wall stress.

### Components of the G protein signalling complex are required for seedling growth during cell wall stress

The 7TM family has previously been annotated as putative GPCRs based on *in silico* predictions from Arabidopsis, poplar and rice (Gookin et al., 2008). These analyses suggested that components of the G protein signalling pathway may also be required for seedling growth under cell wall stress. To this end, we assayed the growth of mutants of the canonical heterotrimeric G protein signalling components, including Gα (GPA1), Gβ (AGB1) and Gγ (AGG1, AGG2 and AGG3) (Ullah et al., 2003; Trusov et al., 2007; Fan et al., 2008; Chakravorty et al., 2011) on media containing isoxaben and other stress conditions (Figure 2; Supplemental Figure 2; Supplemental Figure 3). Similar to the *7tm1* and *7tm1 7tm5* mutants, single knockout mutants for the sole canonical Gβ subunit (*agb1-2*) and a triple mutant for all three Gγ subunits (*agg1-1c agg2-1 agg3-1)* displayed reduced hypocotyl growth and abnormal epidermal cell swelling when grown on isoxaben- and DCB-containing media (Figure 2A & 2B; Supplemental Figure 3). Furthermore, the *agb1-2* mutants displayed a similar reduction in cellulose as the *7tm1* and *7tm1 7tm5* mutant seedlings when grown on isoxaben, but wild type levels of cellulose under normal growth conditions (Figure 2C). These data support a role of the heterotrimeric G protein complex in cell wall signalling or responses to cell wall stress.

**Figure 2.**
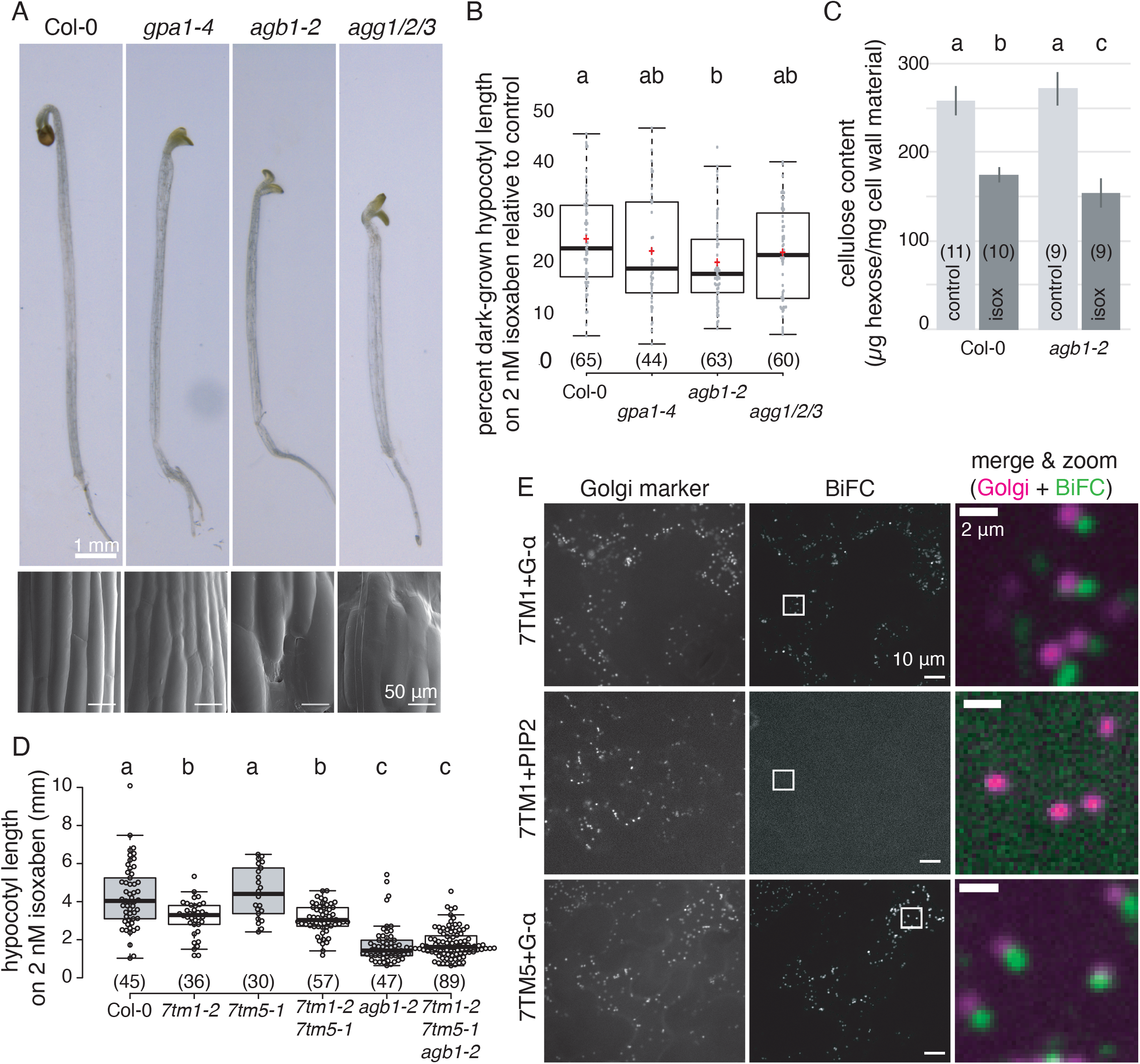
Mutations in components of the G protein signalling complex cause increased sensitivity to cell wall stress. **A.** Representative images of seven-day-old etiolated seedlings and cryo-scanning electron microscopy of 3-day-old etiolated seedlings grown on ½ MS media supplemented with 2 nM isoxaben. **B.** Quantification of etiolated hypocotyl lengths of seedlings represented in (A). **C.** Cellulose content in different genotypes. **D.** Quantification of etiolated hypocotyl lengths of combinatorial mutants between the *7tm* mutants and Gβ mutant *agb1-2.* **E.** Bimolecular Fluorescence Complementation (BiFC) assay in transiently infiltrated *Nicotiana benthamiana* epidermal leaf cells with the BiFC signal shown in green and Golgi-marker in magenta (Merge & Zoom). Experiment was repeated three times with similar results. All graphs summarize three independent experiments, n is indicated in parentheses, samples not sharing a common letter are significantly different (one-way ANOVA and Tukey HSD test, p<0.05), and in box plots, box limits indicate 25^th^ and 75^th^ percentiles, whiskers extend to 1.5 times the interquartile range, median is indicated by a line, mean by a red “+” and individual data points are shown. Data for Col-0 wild type in B & C are the same as in Figure 1, since these experiments were performed together. Scale bars represent 1 mm or 50 μm as indicated in A and 10 μm (two left panel columns) or 2 μm (merge & zoom column) as indicated in E.

To investigate whether the 7TMs and the G proteins work in a common pathway, we produced *7tm1 7tm5 agb1-2* triple mutants and assessed seedling growth on isoxaben-containing media. Consistent with their action in a linear pathway, the triple mutant seedlings displayed similar phenotypes to that of the most severely affected parent line; i.e. the *agb1-2* single mutant seedlings (Figure 2D; Supplemental Figure 5). 7TM1 was previously shown to physically interact with the Gα subunit in a yeast split-ubiquitin interaction system (Gookin et al., 2008). To corroborate these results, we used a modified bimolecular fluorescence complementation (BiFC) system in which both proteins of interest are on the same binary vector, which negates issues of different transformation efficiencies and expression levels of the two candidate interactors (Gookin & Assmann, 2014). We observed clear fluorescent signal when a vector containing both 7TM1-cYFP and Gα-nYFP, or both 7TM5-cYFP and Gα-nYFP, was transformed into *N. benthamiana* leaves (Figure 2E; Supplemental Figure 6). Surprisingly, the BiFC signals were associated with intracellular puncta that resembled the Golgi apparatus, the *trans-*Golgi network (TGN), or other secretory pathway compartments. When this BiFC vector was co-transformed with an RFP-Golgi marker, there was substantial colocalization between YFP and RFP (Figure 2E; Supplemental Figure 6). As a control, we used the modified BiFC system to transform a vector containing 7TM1-cYFP and PIP2-nYFP or 7TM5-cYFP and PIP2-nYFP, since PIP2 is a plasma membrane-localized protein with a similar localization to previous reports of Gα (Chen et al., 2003) and included a mTurquoise-Golgi marker on the same vector backbone as a positive control for transformation. Here, we did not detect any YFP BiFC signal in cells that expressed the Golgi marker (Supplemental Figure 6). Taken together, the common phenotypes, genetic interactions, split-ubiquitin (Gookin et al., 2008) and BiFC data support that 7TM1 and 7TM5 can interact with the G protein signalling complex through Gα, and that normal functions of the Gβ and Gγ subunits are crucial for tolerance to cell wall stress.

### A sub-population of G protein components associates with the Golgi apparatus and TGN

We next investigated the subcellular localization of G protein components. G protein localization data in plants are largely based on constitutive promoter-driven constructs in heterologous systems (Chen et al., 2003; Anderson & Botella, 2007; Zeng et al., 2007), which are prone to localization artefacts. Therefore, we generated native promoter-driven fluorescent protein fusions for Gα, Gβ and all three Gγ subunits and transformed these into their corresponding knockout mutants. Since no signal was detected in native promoter-driven Gα fluorescent protein fusions, we also constructed *Ubiquitin10* promoter-driven GFP fusions for Gα. These constructs partially or fully complemented the respective mutant phenotypes and are thus likely to indicate accurate subcellular localizations (Supplemental Figure 7; Supplemental Figure 8). For all lines, we detected stable fluorescent protein signal at the plasma membrane of young root epidermal cells, where signals were strongest (Supplemental Figure 9). We also detected some intracellular signals in several of the lines; however, this was difficult to visualize due to low fluorescent signal. Therefore, we treated these lines with Brefeldin A (BFA), a fungal toxin that causes aggregation of the TGN and post-Golgi compartments in Arabidopsis root cells (Richter et al., 2007). Treatment with 100 μM BFA for 2h resulted in signal aggregation of GFP-Gα, Gα-GFP and Gβ-mCherry (but not mCherry-Gβ) in intracellular BFA bodies (Supplemental Figure 9), indicating that these proteins can at least partially associate with the TGN or post-Golgi compartments. By contrast, we could not detect signal from the fluorescently tagged Gγ subunits at BFA bodies (Supplemental Figure 9), indicating that either the Gγ are not substantially present at the endomembrane system, or perhaps more likely given that Gβ and Gγ function as Gβγ dimers, that only minute amounts of the proteins associate with the bodies or that the fluorescent tag interfered with the endomembrane localization.

### 7TM1 and 7TM5 act at the Golgi apparatus and *trans-* Golgi network

The Golgi/TGN-localized BiFC signal from the interaction between 7TM1/7TM5 and GPA1 indicated that the 7TMs may be localized to the endomembrane system. To test this, we generated a genomic triple-YFP fusion construct of 7TM1 via recombineering (Zhou et al., 2011) and transformed the construct into *7tm1-2* mutant plants. This construct restored the growth of *7tm1-2* mutant seedlings on isoxaben to nearly wild type levels (Supplemental Figure 10), indicating that the 7TM1-3xYFP fusion was functional. We also generated a functional 7TM5-CFP fusion, driven by a 35S-promoter, and another genomic 7TM1-RFP construct via recombineering, for colocalization purposes. Time-lapse imaging in etiolated hypocotyl cells revealed that 7TM1-3xYFP, 7TM1-RFP and 7TM5-CFP were localized to intracellular puncta that rapidly streamed in the cytoplasm (Figure 3A; Supplemental Movie 1) and that resembled the pattern of proteins localized to the Golgi apparatus or TGN. These results are consistent with proteomic studies that have detected 7TM1 and 7TM5 in the Golgi or TGN-associated proteomes (Dunkley et al., 2006; Parsons et al., 2012; Groen et al., 2013).

**Figure 3.**
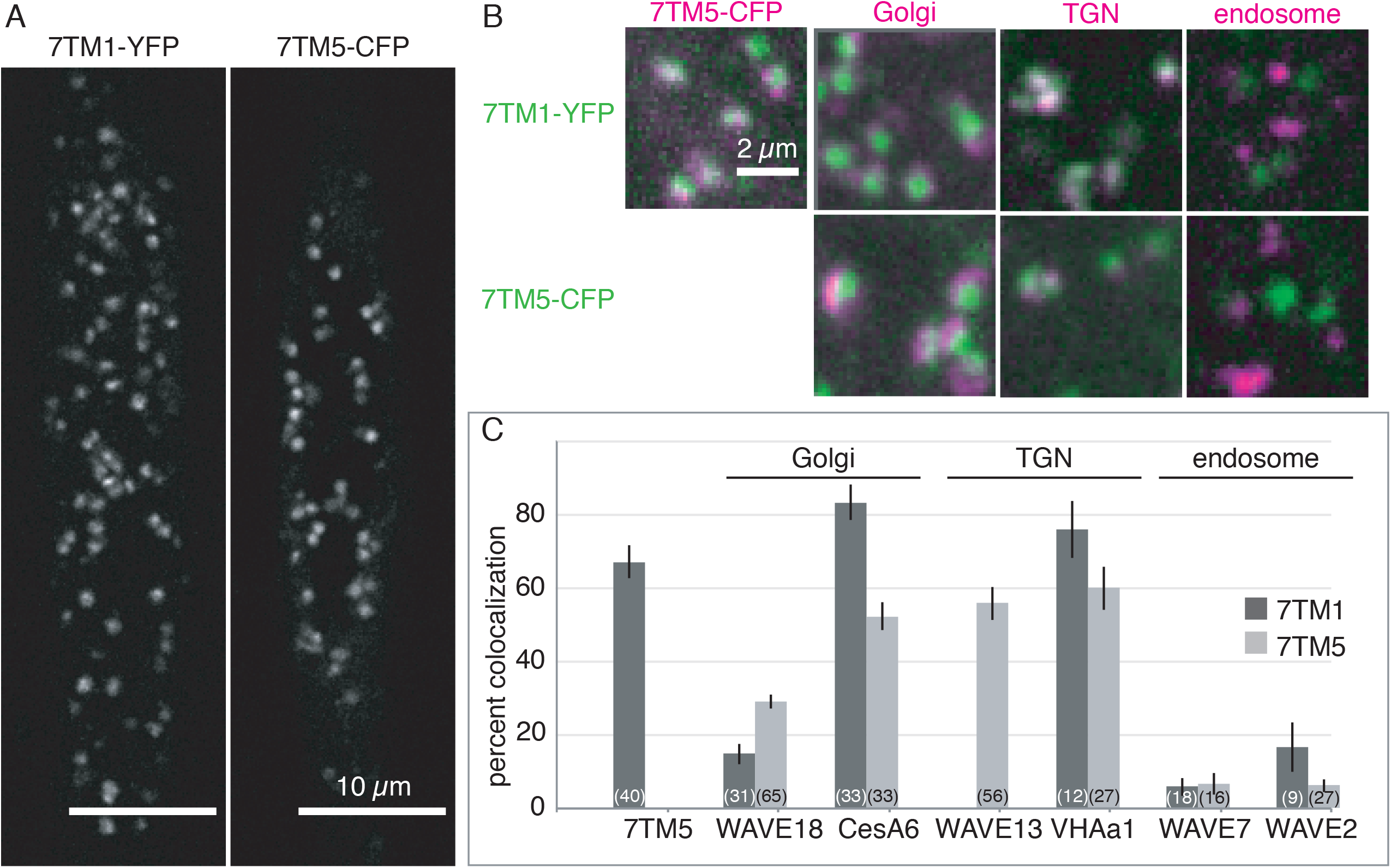
The 7TMs are localized to the Golgi apparatus and *trans*-Golgi network. **A.** Functional 7TM1-3xYFP (7TM1-YFP; native promoter-driven) and 7TM5-CFP (35S-driven) fusion proteins localize to intracellular compartments in 3-day-old etiolated hypocotyl cells. **B.** Representative images from colocalization experiments between 7TM1-3xYFP (green) and 7TM5-CFP (magenta) or between 7TM1-3xYFP or 7TM5-CFP (green) and different endomembrane markers (magenta) as indicated in 3-day-old seedling roots. **C.** Quantification of percent colocalization between 7TM1-3xYFP, 7TM1-RFP, or 7TM5-CFP and endomembrane markers. Graph summarizes three independent experiments, n (cells, no more than three cells imaged per seedling) is indicated in parentheses. Scale bars represent 10 μm in A and 2 μm in B.

To confirm 7TM1 and 7TM5 localization to the Golgi apparatus and TGN, we crossed the 7TM1-3xYFP, 7TM1-RFP and 7TM5-CFP lines to markers for different endomembrane compartments, including the Golgi apparatus (YFP-CESA6 and WAVE18-RFP), the TGN (WAVE13-RFP and VHAa1-mRFP) and the late endosome (WAVE2-RFP and WAVE7-RFP) (Paredez et al., 2006; Dettmer et al., 2006; Geldner et al., 2009). Object-based colocalization was equally high between 7TM1 and 7TM5 with markers for both the Golgi apparatus and the TGN (Figure 3B and C; Supplemental Figure 11), indicating that the steady-state localizations of the 7TM1 and 7TM5 protein fusions are at the Golgi and TGN.

### Overall structure and function of the Golgi apparatus and TGN are not affected in *7tm1 7tm5* and *agb1* mutants

The localization of 7TM1 and 7TM5 to the Golgi and TGN indicates that they may play a role in Golgi or TGN function. To assess this, we first crossed the Golgi marker, NAG-GFP (Grebe et al., 2003) into the *7tm1 7tm5* double mutants. We found no significant differences in NAG-GFP distribution or dynamics in live hypocotyl cells of etiolated *7tm1 7tm5* mutants compared to wild type (Supplemental Figure 12; Supplemental Movie 2). Neither short-term (200 nM for 2h) nor long-term (2 nM for 3 days) isoxaben treatment caused any major alterations to the distribution or behaviour of NAG-GFP in the *7tm1 7tm5* mutant etiolated hypocotyl cells relative to wild type (Supplemental Figure 12).

To determine whether secretion was affected, we crossed sec-GFP into the *7tm1 7tm5* and *agb1-2* mutants. Sec-GFP is a modified GFP with a signal peptide that directs the protein to the secretory pathway and ultimately to the apoplast, where the GFP fluorescence is quenched by the low pH. Because of the stochastic expression of sec-GFP, especially in epidermal cells, a further modified version of the protein was generated in which an endomembrane-targeted RFP is produced in equal amounts to sec-GFP; therefore, the ratio of GFP:RFP can be compared across different plants (Samalova et al., 2006). We found no difference in the ratiometric sec-GFP marker between *7tm1 7tm5*, *agb1-2* and wild type plants, indicating that secretion of this soluble protein is unaffected in the mutants (Supplemental Figure 13). To determine whether secretion of membrane protein cargo was affected in the *7tm1 7tm5* and *agb1-2* mutants, we crossed lines expressing the fluorescent plasma membrane protein GFP-LTI6b (Cutler et al., 2000) into the mutant plants. We found that the steady-state localization of GFP-LTI6b was similar in etiolated hypocotyls of *7tm1 7tm5*, *agb1-2* and wild type plants, and that this localization was unaffected by short-term isoxaben treatment (Figure 4A; Supplemental Figure 14).

**Figure 4:**
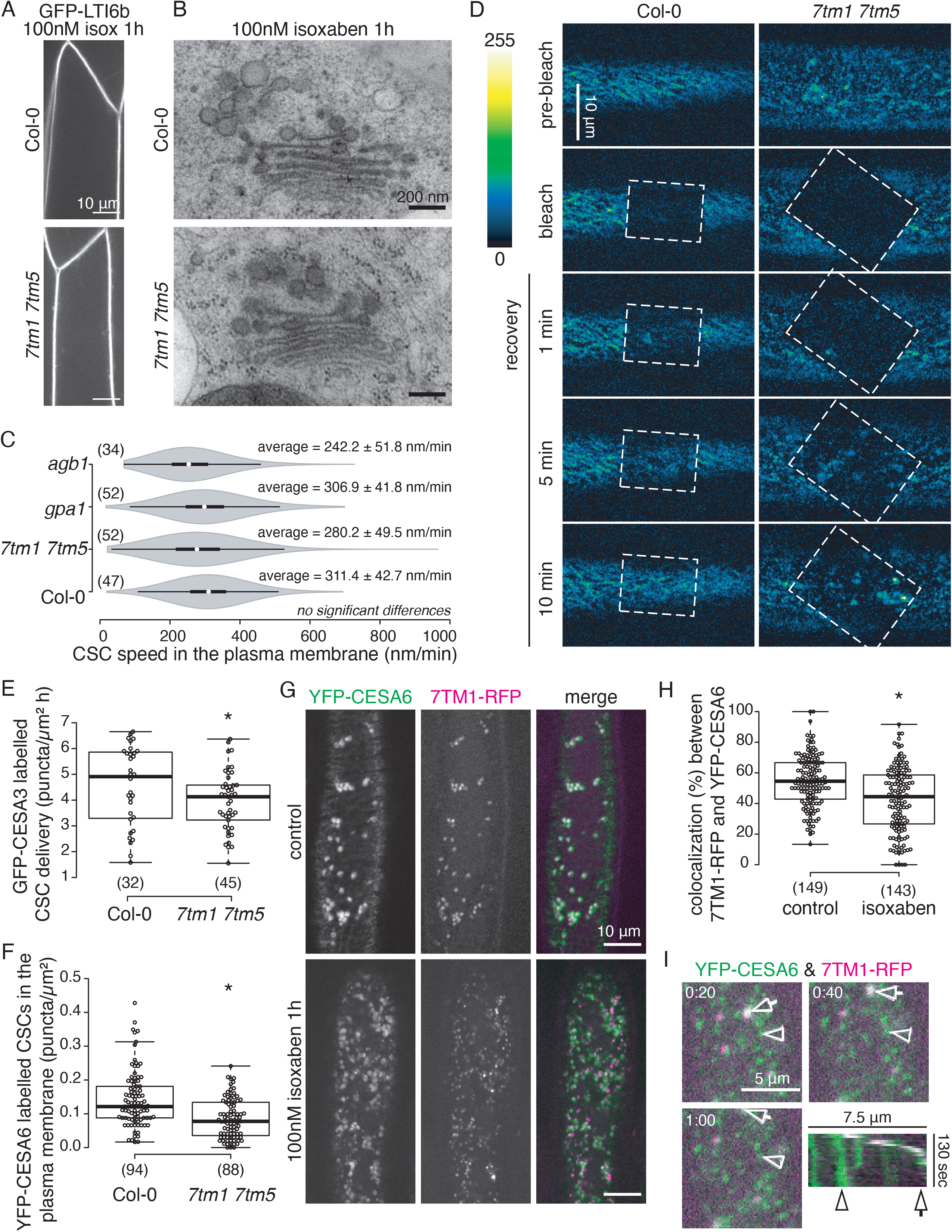
*7tm* mutants are defective in secretion of CSCs, but not other cargos, to the plasma membrane. **A.** Representative images of the plasma membrane marker, GFP-LTI6b in 3-day-old wild type and *7tm1 7tm5* mutant etiolated hypocotyl cells after short-term (100 nM for 1h) isoxaben treatment. **B.** Representative images of the Golgi apparatus in high-pressure frozen, freeze-substituted 3-day-old wild type and *7tm1 7tm5* mutant etiolated hypocotyl cells after short-term (100 nM for 1h) isoxaben treatment. **C.** Quantification of CSC speeds in the plasma membrane of 3-day-old wild type and mutant etiolated hypocotyl cells. **D.** Representative images of photobleaching time-course of GFP-CESA3 in 3-day-old wild type and *7tm1 7tm5* mutant 3-day-old etiolated hypocotyl cells. Bleached area is indicated by dashed box, images are false-coloured according to the scale indicated. **E.** Quantification of the rate of CSC recovery in the plasma membrane after photobleaching in wild type versus *7tm1 7tm5* mutant 3-day-old etiolated hypocotyl cells. **F.** Quantification of YFP-CESA6 labelled CSC particle density in the plasma membrane of 3-day-old etiolated hypocotyl cells in wild type versus *7tm1 7tm5* mutants after long-term isoxaben treatment (2 nM for 3 days). **G.** Representative images of YFP-CESA6 and 7TM1-RFP co-localization in 3-day-old etiolated hypocotyl cells after control or short-term (100 nM for 1h) isoxaben treatment. **H.** Quantification of percent co-localization between YFP-CESA6 and 7TM1-RFP in 3-day-old etiolated hypocotyl cells after control or short-term (100 nM for 1h) isoxaben treatment. **I.** Representative time-lapse images of YFP-CESA6 and 7TM1-RFP co-localization in 3-day-old etiolated hypocotyl cells after control or short-term (100 nM for 1h) isoxaben treatment and kymograph indicating dynamics of YGP-CESA6 and 7TM1-RFP labelled particles; timestamp is in minutes:seconds. All graphs summarize three independent experiments, n (cells, no more than three cells imaged per seedling) is indicated in parentheses and samples marked by an * are significantly different from control (one-way t-test, p<0.05). In violin plot, white circles show the medians, wide bar limits indicate the 25 ^th^ and 75^th^ percentiles, whiskers extend 1.5 times the interquartile range, polygons represent density estimates of data and extend to extreme values. In box plots, box limits indicate 25^th^ and 75^th^ percentiles, whiskers extend to 1.5 times the interquartile range, median is indicated by a line and individual data points are shown. Scale bars represent 10 μm as indicated in A, D and G, 200 nm in B and 5 μm in H.

Since the plant TGN also acts as an early endosome (Dettmer et al., 2006), we evaluated endocytic trafficking to the TGN by tracking the uptake of the fluorescent membrane dye, FM4-64, in seedling roots. There were no substantial differences in either the rate of uptake of FM4-64 or the number of fluorescent puncta at various time points (representing early endosomes/TGNs and subsequently late endosomes) between wild type and either the *7tm1 7tm5* or the *agb1-2* mutants (Supplemental Figure 15). Re-secretion or recycling of proteins to the plasma membrane is also an important function of the plant TGN/early endosome. Therefore, we next evaluated the steady-state localization and re-secretion of the polar auxin transport marker, PIN2-GFP, which we crossed into the *7tm1 7tm5* mutant. In root epidermal cells of wild type and *7tm1 7tm5* mutants, PIN2-GFP was stably localized to the apical surface of the plasma membrane, as previously reported (Xu & Scheres, 2005). Treatment with BFA resulted in PIN2-GFP aggregation in BFA bodies and these BFA bodies were no longer visible after 2 hours of BFA washout in both *7tm1 7tm5* and wild type roots (Supplemental Figure 16).

Finally, we examined the ultrastructure of the secretory pathway in high-pressure frozen, freeze-substituted etiolated seedling hypocotyls. The structures of the Golgi apparatus and TGN were comparable between wild type, *agb1-2* and *7tm1 7tm5* mutant plants (Figure 4B; Supplemental Figure 12). Neither short-term (200 nM for 2h) nor long-term (2 nM for 3 days) isoxaben treatment caused any substantial defects to Golgi or TGN structure in etiolated hypocotyl cells of the *7tm1 7tm5* or *agb1-2* mutants, relative to wild type (Figure 4B; Supplemental Figure 12). These results indicate that there are no large-scale defects in Golgi or TGN structure or function due to loss of 7TM1 and 7TM5 or the G protein complex (via AGB1). These data are consistent with the mostly normal overall growth of *agb1-2* and *7tm1 7tm5* plants under standard growth conditions (Supplemental Figure 2D & 2E), as mutant plants with large Golgi/TGN morphology or secretion defects are usually dwarf, bushy, semi-sterile and exhibit other pleiotropic phenotypes (Teh & Moore, 2007; Gendre et al., 2011; Gendre et al., 2013).

### CSC trafficking is defective in *7tm1 7tm5* and *agb1* mutants

To gain more mechanistic insight into the cellulose defect in *7tm1 7tm5* and G protein mutants, we crossed them with the CESA marker lines YFP-CESA6 (Paredez et al., 2006) or GFP-CESA3 (Desprez et al., 2007) to monitor the behaviour of primary cell wall CSCs. The rate of cellulose synthesis is thought to correspond to the speed of fluorescent CESA puncta movement in the plasma membrane. Consistent with our cellulose assays, we did not detect any significant differences in YFP-CESA6 puncta speed at the plasma membrane in the *7tm1 7tm5, agb1-2* or *gpa1* mutants compared to wild type under control conditions (Figure 4C; Supplemental Figure 17; Supplementary Movie 3). Similarly, we did not observe any differences in steady-state CSC density at the plasma membrane or in SmaCC density in the cortical cytoplasm between wild type and *7tm1 7tm5* mutants under control conditions (Supplemental Figure 17).

Because 7TM1-3xYFP and 7TM5-CFP are localized to the Golgi/TGN, it seemed likely that the Golgi distribution or trafficking of CESAs may be affected in the mutants. To investigate this, we tracked delivery of GFP-CESA3 labelled CSCs from the Golgi apparatus to the plasma membrane using a modified fluorescence recovery after photobleaching (FRAP) approach (Sampathkumar et al., 2013). In wild type etiolated hypocotyl cells, CSCs recovered to nearly normal density within 10 min post-bleach (Figure 4D; Supplemental Figure 18). By contrast, the recovery of CSC density took substantially longer in the *7tm1 7tm5* and *agb1-2* mutants (Figure 4D). Consequently, we found that the rate of CSC insertion into the plasma membrane was significantly lower in the *7tm1 7tm5* and *agb1-2* mutants, relative to wild type (Figure 4E; Supplemental Figure 18).

As short-term isoxaben treatment leads to substantial removal of CESAs from the plasma membrane (DeBoldt et al., 2007), it was intractable to monitor CESA delivery to, and behaviour at, the plasma membrane under those conditions. Therefore, we chose to assess CESA behaviour in seedlings grown on media supplemented with low concentrations of isoxaben. Here, we found substantially fewer GFP-CESA3 labelled CSCs at the plasma membrane in *7tm1 7tm5* double mutants, relative to wild type (Figure 4F). These data suggest that the mutations in the 7TMs and AGB1 impair cellulose synthesis through decreased CSC secretion, which is exacerbated under conditions of cell wall stress.

### 7TM1 localization is biased from Golgi/TGN to SmaCCs upon cell wall stress treatment

Isoxaben induces CSC internalization and/or reduces CSC secretion, resulting in an increase in intracellular CESAs in SmaCCs (Gutierrez et al., 2009), which have been proposed to be involved in either secretion, recycling or degradation of the CESAs. We reasoned that if the 7TMs are actively promoting CSC secretion to the plasma membrane, we may expect to see them colocalize at least in part with some of the isoxaben-induced SmaCCs. We therefore monitored dual-labelled YFP-CESA6 and 7TM1-RFP hypocotyl cells before and after short-term isoxaben treatment (100nM for 1 hour). Under control conditions, the two fluorescent proteins colocalized at the Golgi apparatus and TGN (Figure 2; Figure 4G) and very few SmaCCs were found in the cells (Figure 4G). After 1 h treatment with 100 nM isoxaben, we observed reduced colocalization between the YFP-CESA6 and the 7TM1-RFP signals (Figure 4G and H; Supplemental Figure 19; Supplemental Movie 4). While some the 7TM1-RFP maintained its localization at the Golgi apparatus and TGN, some SmaCCs were labelled by both 7TM1-RFP and YFP-CESA6, and time-lapse imaging revealed erratic movement of these dual-labelled particles, in agreement with previous reports of SmaCC behaviour (Figure 4I; Supplemental Figure 19; Supplemental Movie 4). Interestingly, whereas the highly motile SmaCCs contained both signals, the vast majority of SmaCCs that were stalled at the cell cortex no longer contained 7TM1-RFP signal (Supplemental Figure 19). The stalled SmaCCs are hypothesized to be a final step before CSC insertion into the plasma membrane (Gutierrez et al., 2009; Crowell et al., 2009). Together with our observations that there are no overall changes to Golgi and TGN structure or function in the *7tm1 7tm5* mutants (Figure 4; Supplemental Figures 12-16), these results imply that the 7TM proteins may change their localization upon CWI stress to promote early stages of CESA secretion, prior to CSC insertion to the plasma membrane.

## Discussion

Cellulose is made at the plasma membrane by CSCs and CSC trafficking to and from the plasma membrane represents a critical step in the regulation of cellulose synthesis. By contrast to most plasma membrane-localized proteins, the CESAs display a complex localization pattern, including association with the plasma membrane, Golgi apparatus, SmaCCs and clathrin coated vesicles (Paredez et al., 2006; Crowell et al., 2009; Gutierrez et al., 2009; Sanchez-Rodriguez et al., 2018). This complexity has prompted research into the mechanisms that govern CSC localization and trafficking as an important component of understanding cell wall synthesis. Here, we identified members of a putative GPCR-like protein family, the 7TMs, that are important for CSC secretion and for seedling growth during cell wall stress. Our data indicate that these proteins regulate CSC trafficking from the Golgi apparatus and TGN, and that this role becomes particularly important during cell wall stress.

Yeast assays (Gookin et al., 2008), BiFC data, phenotypic behaviour, cellulose defects in response to cell wall stress and genetic interactions all indicate that 7TM1 and 7TM5 engage with the G protein signalling components. We further found that the corresponding GPCR-like module is localized to the Golgi and TGN based on functional fluorescent protein fusions. Given that a Gβ subunit and a Gγ subunit typically signal together, the absence of a clear fluorescent signal of the mCherry-AGG1, AGG2 or AGG3 at the endomembrane system could indicate that the tag may interfere with some, but not all, of the AGG functions, or that the signal is too dim to reliably detect.

Interestingly, the Golgi/TGN localization of 7TM1 and 7TM5 is reminiscent of an “orphan” family of GPCRs that localize to the Golgi/TGN in animal cells and that may not play a canonical GPCR function (Tafesse et al., 2014). Indeed, the 7TM family are phylogenetically and structurally most similar to these “orphan” GPCR family members (Munk et al., 2019; Supplemental Figure 2B), which include the Golgi and TGN localized GPR107 (Tafesse et al., 2014). Although depletion of GPR107 in HeLa cells did not significantly alter Golgi structure or bulk anterograde trafficking (i.e. secretion) through the secretory pathway, it did substantially affect retrograde trafficking of several markers (Tafesse et al., 2014). Disrupting GPR107 in mouse fibroblast cells resulted in defects in both endocytosis and re-secretion of a subset of cargoes (Zohu et al., 2014). Similarly, overexpression of other members of this “orphan” family, TMEM87A or 87B, was sufficient to rescue trafficking defects in Golgi-associated retrograde protein (GARP) complex disrupted HEK293 cells. The authors reasoned that these Golgi localized human 7TM proteins therefore play a role in retrograde and anterograde trafficking from the Golgi (Hirata et al., 2015). Indeed, in the mammalian G protein signalling field, the notion that GPCRs can be active within the endomembrane system, e.g. at the Golgi apparatus and the TGN, is an exciting and emerging topic (Lobingier & von Zastrow, 2018).

By contrast to GPR107, 7TM1 and 7TM5 do not appear to generally affect Golgi/TGN to plasma membrane trafficking but rather control trafficking of the CESAs between these compartments. Indeed, because *7tm1 7tm5* mutants displayed reduced CSC secretion under normal growth conditions (Figure 4), we propose that the 7TMs are important components of a G protein-mediated mechanism that maintains CSC secretion to the plasma membrane (Figure 5). Interestingly, a sub-population of the isoxaben-induced SmaCCs contained both CESAs and 7TM1. We envision that the SmaCCs consist of a heterogenous population of compartments, which is consistent with observations that different subpopulations of SmaCCs colocalize with different TGN markers (Gutierrez et al., 2009). A progressive change in the SmaCC content may represent a “maturation” of the SmaCCs to prepare to deliver CESAs to the plasma membrane (Figure 5). Consistent with this hypothesis, we found that many of the rapidly moving SmaCCs contained both 7TM1-RFP and YFP-CESA6 signal, while stalled cortical SmaCCs only contained YFP-CESA6. It is plausible that the SmaCCs could form large vesicle “clusters” similar to secretory vesicle clusters (Toyooka et al., 2009) or TGN remnants (Staehelin & Kang, 2008), that can exchange proteins and other materials that may contribute to SmaCC maturation. While the loss of the GPCR-like maintenance mechanism may not be detrimental to plant growth during optimal growth conditions (i.e. seedlings growing on media plates), once challenged with cell wall stress, *7tm1*, *7tm1 7tm5* and several G protein mutant seedlings displayed reduced growth (Figure 1; Figure 2). The sensitivity of the *7tm1 7tm5* and *agb1-2* mutants to isoxaben and DCB, and their reduced capacity to synthesize cellulose under these conditions, indicate that these components are important for modulating CSC secretion during CWI stress. These results imply that the wild type role of the GPCR-like module is to regulate CSC secretion, and therefore contribute to cell wall fortification, especially under CWI stress conditions. Hence, the 7TMs, as part of a potential G protein module, may contribute to maintaining CSC density at the plasma membrane during fluctuating environmental conditions.

**Figure 5:**
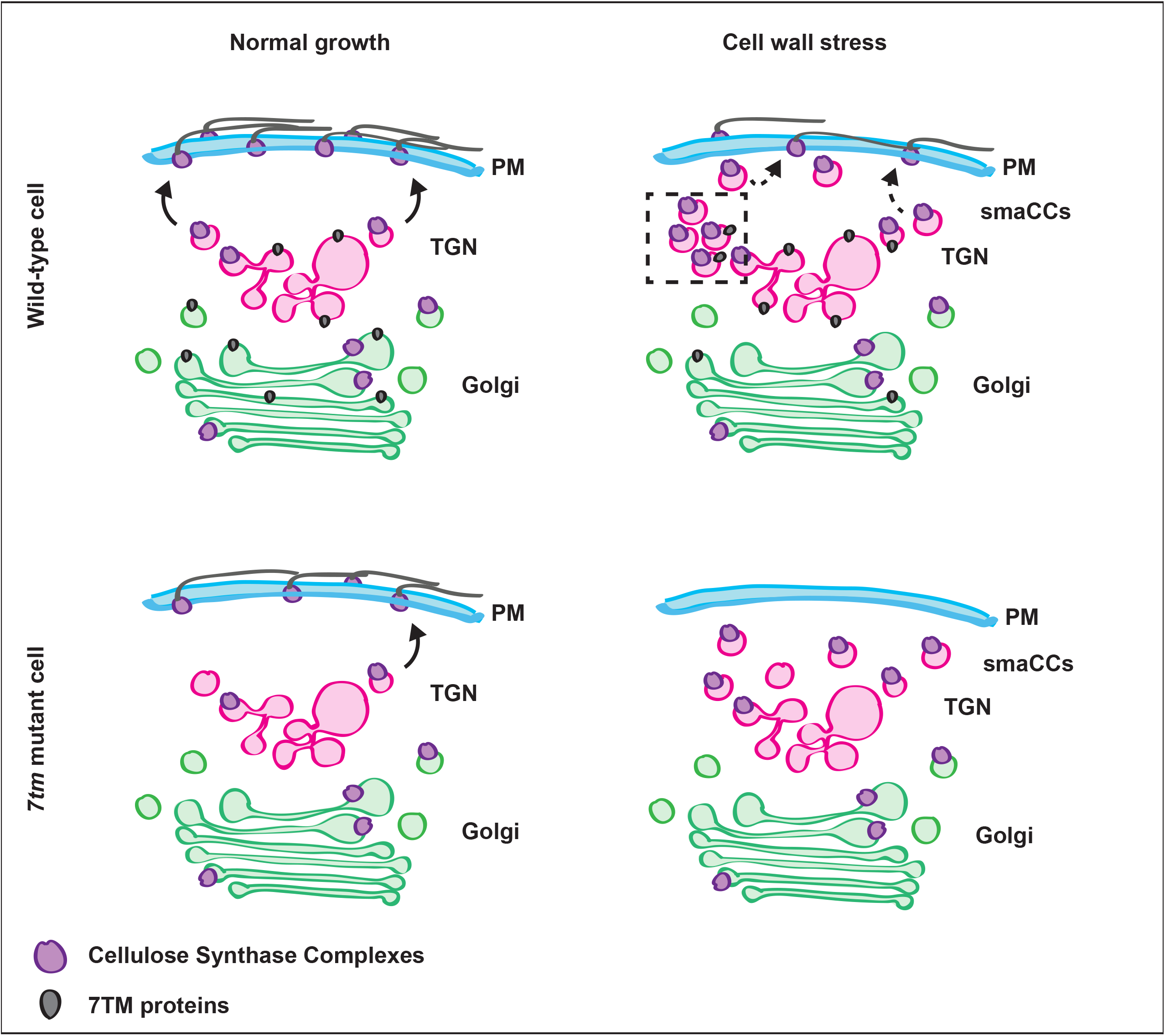
Proposed model for 7TM cellular function. Under normal growth CSCs are actively secreted to the plasma membrane in growing cells. In *7tm1 7tm5* (*7tm*) mutant cells, CSC secretion is reduced. When grown on media supplemented with low concentrations of isoxaben, i.e. cell wall stress, SmaCCs are induced and CSC secretion is maintained at a reduced level in wild type cells, possibly by relocalization of the 7TM proteins to a SmaCC sub-population. Under these conditions, the *7tm* mutant cells are unable to cope with the stress and CSC secretion is substantially reduced. PM; plasma membrane, TGN; trans-Golgi Network, SmaCCs; Small CESA Containing Compartments. Arrows indicate secretion of CSCs, which is reduced in *7tm* mutant cells. Dashed line box indicates a potential vesicle cluster of maturing SmaCCs.

Members of the *Catharanthus roseus* receptor-like kinase-like (CrRLKL) family are associated with CWI signalling in Arabidopsis (reviewed by Engelsdorf et al., 2014) and a number of studies have linked RLK activity to G protein signaling in plants (e.g. Bommert et al., 2013; Aranda-Sicilia et al., 2015; Liang et al., 2016; Tunc-Ozdemir et al., 2016). Notably, FERONIA senses cell wall softening and can engage with cell wall pectins (Feng et al., 2018). FERONIA can interact with the Gβ subunit, AGB1, to control stomatal movement (Yu et al., 2018), which is related to pectin status (Amsbury et al., 2016; Rui et al., 2017). Therefore, FERONIA and its associated signalling components and/or other similar CrRLKLs, are likely to play important roles in the perception of CWI stress at the plasma membrane. The G protein complex may thus constitute an interesting framework for integrating CWI signals and responses. They potentially sense cell wall status via their association with FERONIA at the plasma membrane and regulate CSC secretion via the 7TMs at the endomembrane system.

In summary, we propose that the 7TMs are important maintenance components of plant responses to CWI stress, rather than part of the initial signal perception machinery. Indeed, the *7tm1 7tm5* mutants are hypersensitive to isoxaben, DCB and other cellulose-related defects (Figure 1), which stands in contrast to other CWI signal perception mutants, such as *the1*, which are relatively insensitive to CWI stress (Hematy et al., 2007). In the future it will be interesting to further investigate the extent to which these 7TM-mediated responses interact with previously characterized CWI sensing pathways.

## Methods

### Plant Material and Growth Conditions

*Arabidopsis thaliana* seeds were surface sterilized and sown on ½ MS media (Duchefa) with 2.5 mM MES (Sigma; unless otherwise indicated, all chemicals were from Sigma) at pH 5.8 and 1% sucrose, stratified for 2-3 days in the dark at 4°C, then moved to environmental growth chambers or growth rooms under long-day conditions: 18 h light at ~120 μmol m^−2^ s^−1^ at 21°C and 6 h dark at 18°C. After 7-14 days, seedlings were transferred to soil and returned to the same conditions. For etiolated hypocotyl growth, seeds were exposed to white light (~120 μmol m^−2^ s^−1^ 21°C) for 3 hours, then plates were wrapped in foil and returned to the long-day growth conditions until required for experiments.

For long-term inhibitor experiments, media was supplemented with 0.5-2 nM isoxaben from a 20 μM stock in ethanol, or 200 nM 2,6-Dichlorobenzonitrile (DCB) from a 20 μM stock in ethanol, as indicated. Plates were scanned using an Epson 1000XL flat-bed scanner and hypocotyl lengths were measured using a segmented line in Fiji. Individual hypocotyls were photographed using a Leica M205FA microscope with a DMC4500 camera.

*7tm1-1* (SAIL_701_G12), *7tm1-2* (Salk_134021) and *7tm5-1* (Salk_044297) were obtained from the Arabidopsis Biological Resource Centre. Complete details of all mutants and marker lines are included in Supplemental Table 2. Plants were genotyped via PCR using primers indicated in Supplemental Table 3. For BiFC assays, *Nicotiana benthamiana* seeds were sown directly on soil and grown under the same conditions as Arabidopsis.

### RT-PCR

Seven-day-old light-grown seedlings were flash-frozen in liquid nitrogen and RNA was extracted using the Qiagen Plant RNeasy Mini kit according to the manufacturer’s instructions. RNA integrity was verified via gel electrophoresis. DNA was removed by treatment with DnaseI (Amp-grade, Invitrogen) and cDNA was generated using the SSVI first strand synthesis kit (Invitrogen) with the oligo dT_20_ primer provided according to the manufacturer’s instructions. RT-PCR was performed using primers indicated in Supplemental Table 3.

### Cloning and Plant Transformation

Fragments were amplified from genomic DNA (*7TM1, ABG1, AGG1, AGG2,* AGG3) or cDNA (*7TM5, GPA1, PIP2*) using the primers indicated in Supplemental Table 3.

Whole gene translational fusions between the *7TM1* and the reporters mCherry and 3xYPET were generated by recombineering as described in Brumos et al. (2020). Briefly, the JATY clone JATY76F23 carrying the *7TM1* gene was transferred to the *E. coli* recombineering strain SW105 using the procedures described in Brumos et al. (2020). The mCherry and 3xYPET cassettes (Brumos et al., 2020) were PCR amplified using the primers At5g18520CF and At5g18520CR, inserted in the *At5g18520* gene by recombineering to create in-frame C-terminus translational fusions and the recombinant products confirmed using the primers At5g18520CTF and At5g18520CTR. Finally, these DNA constructs were trimmed using the primers At5g18520delRB and At5g18520delLB and the deletions confirmed using the primers At5g18520delTRB and At5g18520delTLB. All primers are indicated in Supplemental Table 3.

For *35S* promoter-driven 7TM5-CFP, *7TM5* coding sequence was amplified without the stop codon using the primers indicated in Supplemental Table 3 and transferred into the pEZR(K)-LNC binary vector via the EcoRI and BamHI sites.

For *Ubiquitin10* promoter-driven GFP-GPA1 and GPA1-GFP, *GPA1* coding sequence was amplified with or without the stop codon, respectively, from cDNA using the primers indicated in Supplemental Table 3 and transferred into pENTR-D-TOPO (Invitrogen) via Gibson assembly using NEB Gibson Assembly Master Mix according to the manufacturer’s instructions. The GPA1 cassettes with and without the stop codon were then transferred to the pUBN-eGFP and pUBC-eGFP binary vectors, respectively (Grefen et al., 2010), via a Gateway LR reaction using LR Clonase II (Invitrogen) according to the manufacturer’s instructions.

For native promoter-driven AGB1-mCherry, the *AGB1* genomic region from the start codon to immediately before the stop codon was inserted into pENTR-D-TOPO (Invitrogen) via TOPO cloning according to the manufacturer’s instructions. Then the *AGB1* upstream regulatory region (~1.6 kb upstream of ATG, including the 5` UTR) was inserted into the Not1 site of pENTR-D-TOPO and mCherry, including a stop codon, was inserted into the AscI site while also introducing a downstream KpnI site, and the downstream *AGB1* regulatory region (~400bp of 3` UTR) was inserted into this KpnI site.

For native promoter-driven mCherry-AGB1, the coding sequence of *mCherry* without a stop codon was inserted into pENTR-D-TOPO (Invitrogen) via TOPO cloning according to the manufacturer’s instructions. Then, the upstream *AGB1* regulatory region (~1.6 kb upstream of ATG, including the 5` UTR) was inserted into the Not1 site of pENTR-D-TOPO and the genomic sequence of AGB1 (from start codon to ~400bp downstream of the stop codon) was inserted into the AscI site.

For native promoter-driven mCherry-AGG1, mCherry-AGG2 and mCherry-AGG3, the same strategy was taken as for mCherry-AGB1; see Supplemental Table 3 for primers.

All of the *AGB1* and *AGG* cassettes were transferred to the pMDC99 binary vector (Curtis & Grossniklaus 2003) via a Gateway LR reaction using LR Clonase II (Invitrogen) according to the manufacturer’s instructions.

All constructs were verified via Sanger sequencing. Vectors were electroporated into *Agrobacterium tumefaciens* strain GV3101 and selected in half-salt LB media with appropriate antibiotics.

For stable Arabidopsis transformations, Agrobacteria from an overnight culture were harvested by centrifugation, then resuspended in a solution of 5% sucrose and 0.05% silwet L-77 (PhytoTech Labs). Young (3-4 week-old) Arabidopsis plants with floral buds were submerged in this solution for one minute, then returned to growth conditions. Seedlings were selected for positive transformants on 50 μg/mL kanamycin, 15 μg/mL glufosinate ammonium, or 30 μg/mL hygromycin, as appropriate. T3 lines that were likely homozygous for single insertions were selected based on segregation of the selection marker.

For BiFC, Agrobacteria from an overnight culture were harvested by centrifugation, then resuspended in a solution of 10 mM MES, 10 mM MgSO_4_ and 75 μM acetosyringone (from a 100 mM stock in DMSO) to a final concentration of OD_600_ = 0.03 for each construct. *N. benthamiana* plants (4-6 week-old) were infiltrated by leaf injection. Duplicate infiltrations were performed on two separate plants per assay in each independent experiment. 48h post-infiltration, small regions of the leaf were excised with a razor blade and mounted in water for imaging via spinning disk microscopy.

### Live Cell Imaging and Image Analysis

Live cell imaging for all figures was conducted using a Nikon Ti-E equipped with an Andor Revolution CSU-W1 spinning disk, a Borealis homogeneous illumination system, an Andor FRAPPA photobleaching unit, an Andor Ixon Ultra 888 EM-CCD and a 100x or 60x N.A. 1.49 Apo TIRF oil-immersion objectives. GFP was excited with a 488 nm laser and emission collected with a 525/50 nm band pass filter, YFP was excited with a 515 nm laser and emission collected with a 535/30 nm filter, CFP and mTurquoise were excited with a 445 nm laser and emission collected with a 470/24 nm filter, tdTomato, mCherry and RFP were excited with a 561 nm laser and emission collected with a 610/40 nm filter. Alternatively, some images for quantification were collected with a Nikon Ti-E equipped with an Andor CSU-X1 spinning disk, a BioVision iLAS FRAP photobleaching unit, a Photometrics Evolve EM-CCD and a 100x N.A. 1.49 Apo TIRF oil-immersion objective. GFP and YFP were excited with a 491 nm laser and emission collected with a 525/50 nm filter; tdTomato and RFP were excited with a 561 nm laser and emission collected with a 595/50 nm filter.

For seedling imaging, 3-day-old etiolated hypocotyls or roots were mounted in water under a pad of 0.8% agarose (Bioline). To limit the time that seedlings spent mounted, no more than three cells per seedling were imaged in any experiment. For short-term inhibitor treatments, seedlings were incubated for the time indicated, gently shaking in a 6-well plate containing ½ MS media with 2.5 mM MES pH 5.8 and 1% sucrose, supplemented with 100 nm isoxaben (from a 20 μM stock in ethanol) or 5 μm 2,6-Dichlorobenzonitrile (DCB) (from a 20 μM stock in ethanol), or 100 μM Brefeldin A (BFA) (from a 100 mM stock in DMSO), then mounted in the same medium under an agarose pad as described above.

For FM4-64 uptake assays, seedlings for short time points were directly mounted in water containing 4 μM FM4-64 (from a 4 mM stock in DMSO), while seedlings for long time points were incubated in 4 μM FM4-64 in water for 5 minutes, then transferred to a 6-well plate containing ½ MS media with 2.5 mM MES pH 5.8 and 1% sucrose for the remainder of the time point before mounting and imaging. FM4-64 signal was excited with a 488 nm laser and emission collected with a 625/90 nm filter.

CESAs were monitored using 10 second time-lapse images for 10 minutes and speeds were determined in Fiji. If required, time-lapse images were aligned using the StackReg plugin (Thevenaz et al., 1998) and background was subtracted from images using a 50 pixel rolling ball radius. Sum projections of time-lapse images were used to select CSC tracks, then kymographs (3 pixel line width) were generated from the time series using the MultiKymograph tool. CSC speeds were calculated from the displacement over time of individual particles identified in the kymographs (Paredez et al., 2006). CSC density at the plasma membrane and SmaCCs in the cell cortex were identified using the ThunderSTORM plugin (Ovesný et al., 2014) with the microscope hardware information, but otherwise using default settings. Golgi bodies in the cell cortex were manually excluded by their large size, SmaCCs were identified as high-intensity particles and the remainder of the particles were considered to be CSCs.

Photobleaching was conducted with an Andor FRAPPA unit on the microscope described above, using the 488 nm laser, a 20 ms dwell time and 80% laser power; otherwise imaging conditions were as described above for CESA monitoring. New CSC delivery events were identified according to Sampathkumar et al. (2013).

Colocalization was quantified from z-stacks (with 0.2 μm spacing using the 100x N.A. 1.49 objective) using the object-based (centre-particle) method in the JaCOP plugin for Fiji (Bolte & Cordelieres, 2006) with the microscope hardware information, but otherwise default settings.

### Transmission Electron Microscopy

Etiolated 3-day-old seedlings were cryofixed using a Leica EM-ICE high pressure freezer using B-type carriers and 1-hexadecene as a cryoprotectant, according to McFarlane et al. (2008) with modifications. Samples were freeze-substituted in a Leica AFS2 automatic freeze substitution unit at −85°C for 4 days in 2% (w/v) osmium tetroxide (Electron Microscopy Sciences) and 8% 2,2-dimethoxypropane (w/v) in anhydrous acetone, after which the temperature was gradually raised to room temperature over 2 days. Samples were washed 5 times with anhydrous acetone, then infiltrated with Spurr’s Resin (Electron Microscopy Sciences) over the course of 4 days. Samples in resin were polymerized at 65°C for 36 hours in BEEM capsules (Electron Microscopy Sciences). Silver (~80 nm thick) sections were cut using a Leica UCT R or a UC7 Ultramicrotome and a DiATOME diamond knife, placed on Gilder fine bar hexagonal 200 mesh grids coated with 0.3% formvar (Electron Microscopy Sciences). Grids were post-stained with 1% aqueous uranyl acetate (Polysciences) and Sato’s triple lead (Sato, 1968; sodium citrate, lead acetate, lead citrate from BDH, lead nitrate from Fisher) and imaged with a Phillips CM120 BioTWIN transmission electron microscope with a Gatan MultiScan 791 CCD camera and a tungsten filament at an accelerating voltage of 120 kV.

### Scanning Electron Microscopy

Etiolated 3-day-old seedlings were mounted on a sample holder using Tissue-Tek (Sakura-Finetek), plunge-frozen in a slush of liquid nitrogen (~-210°C), then transferred to a Gatan cryostage. Ice crystals were evaporated at −95°C for 2.5 minutes, then samples were coated with 60:40 gold-palladium alloy for 120 sec (∼6 nm) under argon at −120°C, before being transferred into the FEI Quanta cryo scanning electron microscope. Stage temperatures were maintained below −120°C and images were collected at a 5 kV accelerating voltage and a 10 mm working distance using the E-T detector.

### Cell Wall Analysis

Cell wall analysis was conducted according to the protocol described in Sanchez-Rodriguez et al., 2012 without any modifications.

### Phylogenetic Analysis

Full length protein sequences were analyzed using MEGA X (Kumar et al., 2018) and included software packages. Sequences were aligned using MUSCLE and unrooted phylogenetic trees were constructed via the Maximum Likelihood method using 391 positions after complete deletion of gaps and missing data. The reliability of the inferred tree was evaluated using 100 bootstrap replicates.

## Supporting information

McFarlane et al Supplemental Data

## Acknowledgements

Live cell imaging was conducted using instruments that are part of the Biological Optical Microscopy Platform (BOMP) at University of Melbourne and electron microscopy was conducted using instruments that are part of the Melbourne Advanced Microscopy Facility. S.P. acknowledges the financial aid of an ARC Future Fellow Grant (FT160100218), an ARC Discovery grant (DP19001941) and an IRRTF grant from UoM. H.E.M. acknowledges an EMBO-LTF (1246-2013), NSERC PDF (454454-2014) and an ARC Discovery Early Career Researcher Award (DE170100054). T.E.G. and S.M.A. acknowledge support from the U.S. National Science Foundation (MCB-1121612, with additional support from MCB-1715826), JMA acknowledges support from NSF grant IOS1444561 and D.M-A. acknowledges financial support from Deutsche Forschungsgemeinschaft.

## Supplemental Figure Legends

**Supplemental Figure 1: *7TM1* is co-expressed with primary wall *CESA* genes. A.** Gene family-based co-expression network. Nodes represent gene families (pfam) and edges (lines) connect co-expressed gene families. Edge colors indicate number of species where gene families are co-expressed: blue line for 2 species, green line for 3 species, yellow line for 4 species, red line for 5 species, black line for > 5 species. Species as per Ruprecht *et al.* (2016). The Lung_7TM-R pfam includes the *7TM* genes and is highlighted in red. **B.** Gene co-expression network centred around Arabidopsis primary cell wall *CESA1*. Cellulose-associated gene names are indicated as acronyms: *KOB* (*Kobito*), *CSI1* (*Cellulose Synthase Interacting1*), *CTL1* (*Chitinase-like1*), *KOR* (*Korrigan*), *CC1* (*Companion of Cellulose synthase1*). Node colors represent different gene families as per FamNet using *CESA1* as search (http://www.gene2function.de/famnet.html). *7TM1* (At5g18520) is highlighted.

**Supplemental Figure 2: Mutations in 7TM family members do not cause general plant growth phenotypes as compared to wild type. A.** RT-PCR of *7TM1* and *7TM5,* and *ACT2* as a control in wild type Col-0 compared to corresponding mutants. **B.** Phylogenetic assessment of full-length Arabidopsis and human protein sequences. Evolutionary analysis shows a bifurcation of the LUSTR family. Sequences were aligned using MUSCLE and the tree was constructed using Maximum Likelihood with 100 bootstrap replicates. Tree branch lengths are drawn to scale (substitutions per site). *A. thaliana* sequences: 7TM1 (At5g18520.1), 7TM2 (At3g09570.1), 7TM3 (At5g02630.1), 7TM4 (At5g42090.1), 7TM5 (At2g01070.1). *Homo sapiens* protein sequences: GPR107 (NP_001130029.1), GPR108 (NP_001073921.1), TMEM87A isoform 1 (NP_056312.2), TMEM87B isoform 1 (NP_116213.1). **C.** Quantification of hypocotyl lengths of 7-day-old etiolated seedlings grown on ½ MS media supplemented with 2 nM isoxaben. Box limits indicate 25^th^ and 75^th^ percentiles, whiskers extend to 1.5 times the interquartile range, median is indicated by a line, mean by a red “+”, individual data points are shown, n is indicated in parentheses, and samples marked by an * are significantly different from wild type (one-way t-test, p<0.05). **D.** Representative images of 7-day-old etiolated seedlings grown on ½ MS media. **E.** Representative images of 4-week-old plants grown under unstressed, long-day conditions.

**Supplemental Figure 3 *7tm* mutants are sensitive to cell wall synthesis inhibitors, but not to other stresses.** Representative images of 7-day-old etiolated seedlings grown on ½ MS media supplemented with various inhibitors as indicated and quantification of hypocotyl length; n is indicated in parentheses, samples not sharing a common letter are significantly different (one-way ANOVA and Tukey HSD test, p<0.05).

**Supplemental Figure 4: *7tm1 prc1-1* double mutants show slight decrease in growth as compared to *prc1-1*.** Representative images of 7-day-old etiolated seedlings grown on ½ MS media and quantification of etiolated hypocotyl lengths; n is indicated in parentheses, samples not sharing a common letter are significantly different (one-way ANOVA and Tukey HSD test, p<0.05), white circles show the medians, wide bar limits indicate the 25^th^ and 75^th^ percentiles, whiskers extend 1.5 times the interquartile range, polygons represent density estimates of data and extend to extreme values.

**Supplemental Figure 5: Genetic evidence indicates that 7TM1/7TM5 and AGB1 work in a linear pathway.** Representative images of 7-day-old etiolated seedlings grown on ½ MS media supplemented with 2 nM isoxaben and quantification of etiolated hypocotyl lengths; n is indicated in parentheses, samples not sharing a common letter are significantly different (one-way ANOVA and Tukey HSD test, p<0.05), box limits indicate 25^th^ and 75^th^ percentiles, whiskers extend to 1.5 times the interquartile range, median is indicated by a line, and individual data points are shown. The same data for Col-0 wild type, *7tm1-2*, *7tm5-1* and *7tm1 7tm5* double mutants is shown in both graphs since these controls were grown together with the G protein and triple mutants.

**Supplemental Figure 6: 7TMs and Gα interact in BiFC assays.** Bimolecular Fluorescence Complementation (BiFC) assay in transiently infiltrated *Nicotiana benthamiana* leaf epidermal cells. PIP2, which, similar to the G proteins primarily localizes to the plasma membrane, was substituted for Gα as a negative control. BiFC signal is shown in green and Golgi-marker in magenta in the merge and zoom images; white boxes in the merge images indicate the region in the zoom images.

**Supplemental Figure 7: Fluorescently tagged G protein constructs complement corresponding G protein mutant plants.** Representative images of 7-day-old etiolated seedlings from homozygous, single insertion T3 lines (identified based on segregation of the selection marker) in the corresponding mutant background grown on ½ MS media supplemented with 2 nM isoxaben, and quantification of etiolated hypocotyl lengths. Independent transformants are separated, n is indicated in parentheses, samples not sharing a common letter are significantly different (one-way ANOVA and Tukey HSD test, p<0.05), box limits indicate 25^th^ and 75^th^ percentiles, whiskers extend to 1.5 times the interquartile range, median is indicated by a line, and individual data points are shown.

**Supplemental Figure 8: Fluorescently tagged G protein constructs complement corresponding G protein mutant plants.** Representative images of 7-day-old etiolated seedlings from homozygous, single insertion T3 lines (identified based on segregation of the selection marker) in the corresponding mutant background grown on ½ MS media supplemented with 2 nM isoxaben and quantification of etiolated hypocotyl lengths. Independent transformants are separated, n is indicated in parentheses, samples not sharing a common letter are significantly different (one-way ANOVA and Tukey HSD test, p<0.05), box limits indicate 25^th^ and 75^th^ percentiles, whiskers extend to 1.5 times the interquartile range, median is indicated by a line, and individual data points are shown, green line indicates wild type median, magenta line indicates *agg1-1c agg2-1 agg3-1* triple mutant median.

**Supplemental Figure 9: G protein components are localized to the plasma membrane and intracellular compartments.** Representative images from 3-day-old seedling root cells of GPA1-GFP, GFP-GPA1 (both *Ubiquitin10* promoter-driven), AGB1-mCherry, mCherry-AGB1, mCherry-AGG1, mCherry-AGG2, mCherry-AGG3 (all native promoter-driven) single insertion T3 lines (identified based on segregation of the selection marker) in the corresponding mutant background under control conditions, or after 2 hour treatment with 100 μM brefeldin A (BFA).

**Supplemental Figure 10: Fluorescently tagged 7TM constructs complement corresponding** *7tm* **mutant plants.** Representative images of 7-day-old etiolated seedlings from homozygous, single insertion T3 lines (identified based on segregation of the selection marker) in the corresponding mutant background grown on ½ MS media supplemented with 2 nM isoxaben and quantification of etiolated hypocotyl lengths. Independent transformants are separated, n is indicated in parentheses, samples not sharing a common letter (lowercase for 7TM1, uppercase for 7TM5) are significantly different (one-way ANOVA and Tukey HSD test, p<0.05), box limits indicate 25^th^ and 75^th^ percentiles, whiskers extend to 1.5 times the interquartile range, median is indicated by a line, mean by a red “+”, and individual data points are shown.

**Supplemental Figure 11: The 7TMs are localized to the Golgi apparatus and the *trans*-Golgi network.** Representative images from colocalization experiments between 7TM1-YFP and 7TM5-CFP or between 7TM1-YFP, 7TM1-RFP, or 7TM5-CFP, and different endomembrane markers as indicated in 3-day-old seedling roots.

**Supplemental Figure 12: Golgi apparatus organization and structure are not affected *7tm1 7tm5* or *agb1* mutants.** Representative images of the NAG-GFP Golgi marker and of the Golgi apparatus in high-pressure frozen, freeze-substituted transmission electron microscopy samples from wild type, *7tm17tm5* double mutant and *agb1-2* single mutant 3-day-old etiolated hypocotyls under control conditions compared to short-term (200 nM for 2 hours) and long-term (2 nM for 3 days) isoxaben treatment.

**Supplemental Figure 13: Localization of sec-GFP is not affected in *7tm1 7tm5* or *agb1* mutants.** Representative images of the ratiometric sec-GFP marker (35S:stN-RM-2A-sec-GFP) crossed into and *7tm1 7tm5* and *agb1-2* mutants, compared to wild type 3-day-old etiolated hypocotyls. Green channel represents sec-GFP signal as it is transported through the secretory pathway, magenta channel represents RFP that is expressed at the same level and targeted to the vacuole via the Golgi apparatus.

**Supplemental Figure 14: Localization of GFP-LTI6b is not affected in *7tm1 7tm5* or *agb1* mutants.** Additional representative images of the plasma membrane marker 35S:GFP-LTI6b in 3-day-old etiolated hypocotyl cells of *7tm1 7tm5*, *gpa1-4*, and *agb1-2* mutants, compared to wild type under control conditions or after short-term (200 nM for 1 hour) isoxaben treatment.

**Supplemental Figure 15: Time course of FM4-64 uptake reveals that uptake is not affected in *7tm1 7tm5* and *agb1* mutants.** Representative time course images of membrane dye (FM4-64) uptake into in root cells of 3-day-old seedlings of *7tm1 7tm5* and *agb1-2* mutants, compared to wild type. FM4-64 initially labels the plasma membrane, then travels to the early endosome, late endosome, and the tonoplast (vacuole membrane) over 3 hours. Images are false-coloured according to the scale.

**Supplemental Figure 16: Time course of PIN2-GFP localization after BFA treatment and washout reveals that localization is not affected in *7tm1 7tm5* mutants.** Representative time course images of PIN2-GFP localization before and after 2 hour treatment with 100 μM brefeldin A (BFA), and after a time course of BFA washout in root cells of 3-day-old *7tm1 7tm5* mutant seedling roots, compared to wild type.

**Supplemental Figure 17: YFP-CESA6 movement and density at the plasma membrane are unaffected in *7tm 7tm5* and G protein mutants.** Representative images of YFP-CESA6 crossed into *7tm1 7tm5*, *gpa1-4* and *agb1-2* mutants, compared to wild type in in 3-day-old etiolated hypocotyl cells. Data are represented as single frames from the time lapse, a sum projection of the data (from 10 minute time lapse with 10 second intervals) and a kymograph from the magenta line indicated in the single frame. CESA speeds in the plasma membrane were quantified using the kymographs while CESA density at the plasma membrane and SmaCC density in the cortical cytoplasm were quantified from single frames. Graphs represent data from three independent experiments, n (cells, no more than three cells imaged per seedling) is indicated in parentheses, samples not sharing a common letter are significantly different (one-way ANOVA and Tukey HSD test, p<0.05), box limits indicate 25^th^ and 75^th^ percentiles, whiskers extend to 1.5 times the interquartile range, median is indicated by a line, and individual data points are shown.

**Supplemental Figure 18: Photobleaching experiments reveal reduced GFP-CESA3 delivery to the plasma membrane in *7tm1 7tm5* and *agb1* mutants compared to wild type.** Representative images of GFP-CESA3 crossed into *7tm1 7tm5*, *gpa1-4* and *agb1-2* mutants, compared to wild type in 3-day-old etiolated hypocotyl cells. Bleached area is indicated by dashed box, scale bars represent 10 μm. Images are single frames from the time lapse or a sum projection of the data (from 10 minute time lapse with 10 second intervals), as indicated. CESA-labelled CSC particles secreted to the plasma membrane were quantified according to Sampathkumar et al. (2013). Graphs represent data from two (*gpa1-4* and *agb1-2*) or three (Col-0 and *7tm1 7tm5*) independent experiments, n (cells, no more than three cells imaged per seedling) is indicated in parentheses, box limits indicate 25^th^ and 75^th^ percentiles, whiskers extend to 1.5 times the interquartile range, median is indicated by a line, individual data points are shown, and samples marked by an * are significantly different from control (one-way t-test, p<0.05).

**Supplemental Figure 19: YFP-CESA6 and 7TM1-RFP co-localize in rapidly moving SmaCCs.** Montage image of YFP-CESA6 and 7TM1-RFP colocalization in wild type 3-day-old etiolated hypocotyl cells treated with 100 nM isoxaben for 1 hour. Arrow indicates a fast-moving SmaCC containing both YFP-CESA6 and 7TM1-RFP signal; arrowhead indicates a stalled SmaCC that contains only YFP-CEAA6. Scale bar represents 2 μm.

## Supplemental Movies

**Supplemental Movie 1: Time-lapse imaging of 7TM1-YFP in etiolated hypocotyl cells**.

**Supplemental Movie 2: Time-lapse imaging of NAG-GFP in wild type and *7tm1 7tm5* etiolated hypocotyl cells**.

**Supplemental Movie 3: Time-lapse imaging of YFP-CESA6 in wild type and***7tm1 7tm5*

**etiolated hypocotyl cells**.

**Supplemental Movie 4: Time-lapse imaging of YFP-CESA6 and 7TM1-RFP in control and isoxaben-treated etiolated hypocotyl cells.**

## Supplemental Tables

**Supplemental Table 1: Co-expression gene list using***7TM1* **as bait (from ATTED-II; Obayashi et al., (2018)**.

**Supplemental Table 2: A list of seed lines employed in this study**.

**Supplemental Table 3: A list of primers employed in this study**.

