## Supplementary material for "A G protein-coupled receptor-like module regulates Cellulose Synthase secretion from the endomembrane system in Arabidopsis": McFarlane et al Supplemental Data

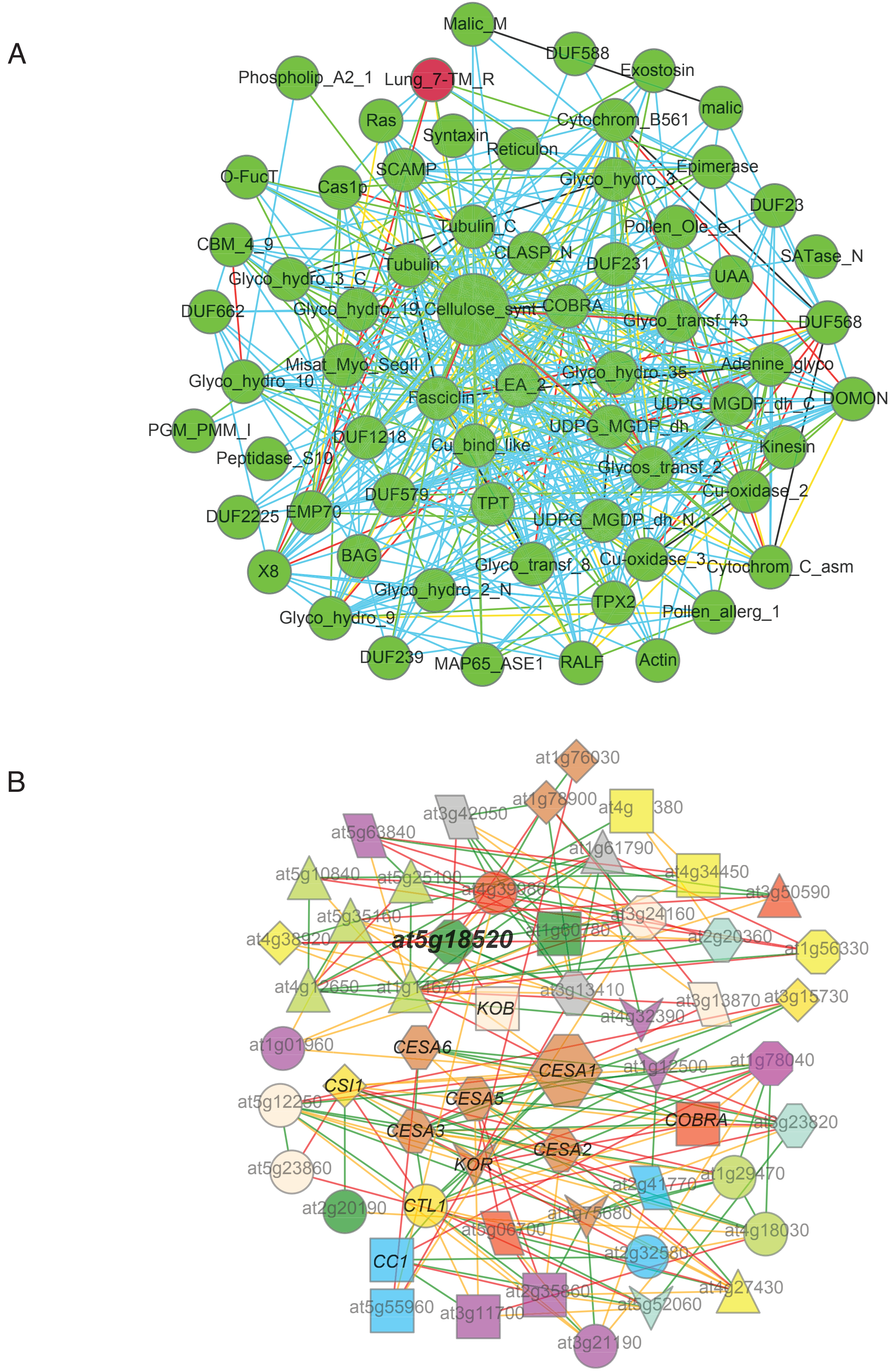


**Supplemental Figure 1: *7TM1* is coexpressed with primary wall *CESA* genes. A.** Gene family-based coexpression network. Nodes represent gene families (pfam) and edges (lines) connect coexpressed gene families. Edge colors indicate number of species where gene families are coexpressed: blue line for 2 species, green line for 3 species, yellow line for 4 species, red line for 5 species, black line for > 5 species. Species as per Ruprecht et al. (2016). The Lung_7TM-R pfam includes the *7TM* genes and is highlighted in red. **B.** Gene co-expression network centred around Arabidopsis primary cell wall CESA1. Cellulose-associated gene names are indicated as acronyms *KOB* (*Kobito*), *CSI1* (*Cellulose Synthase Interacting1*), *CTL1* (*Chitinase-like1*), *KOR* (*Korrigan*), *CC1* (*Companion of Cellulose synthase1*). Node colors represent different gene families as per FamNet using *CESA1* as search (http://www.gene2function.de/famnet.html). *7TM1* (At5g18520) is highlighted.

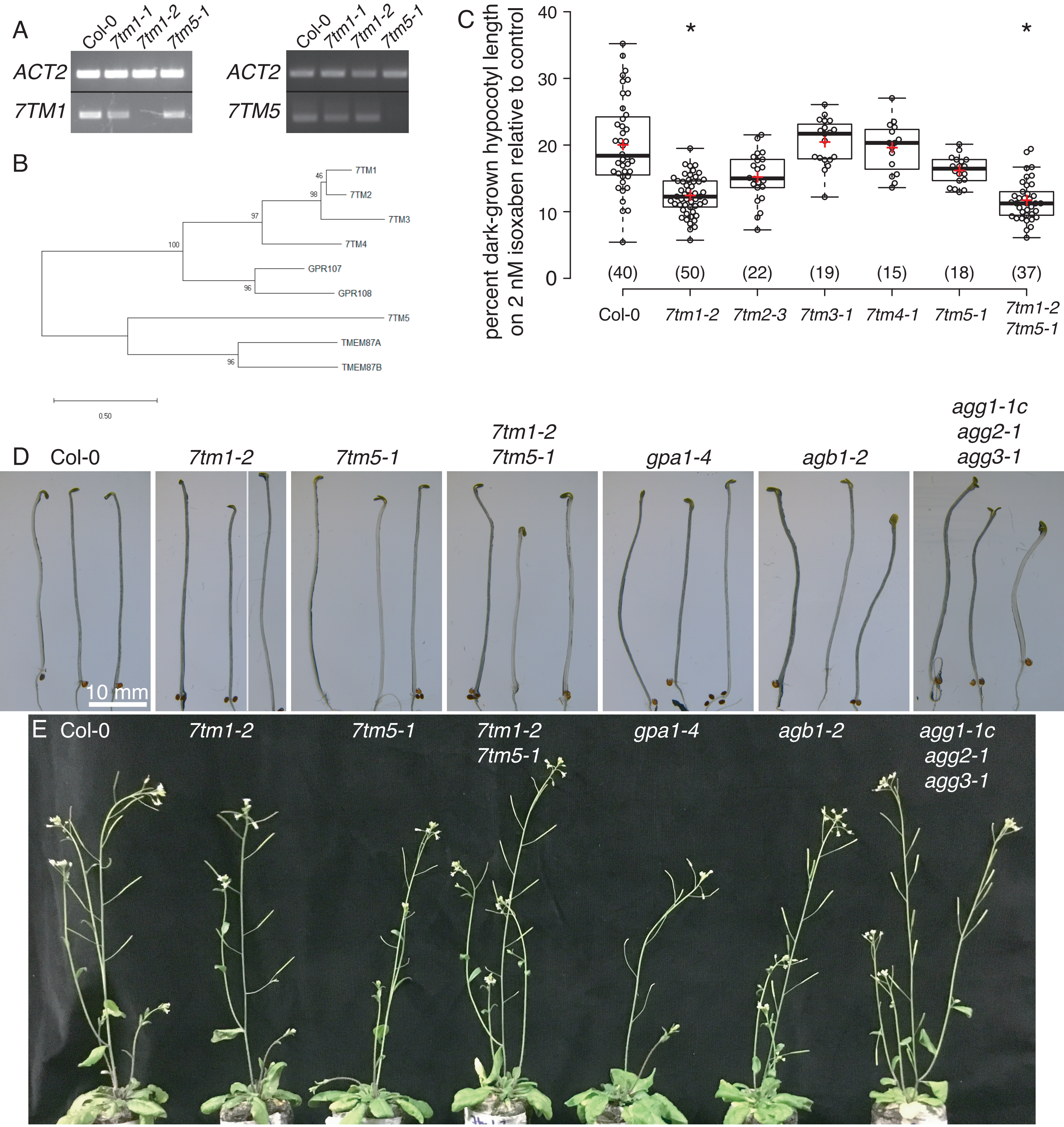


**Supplemental Figure 2: Mutations in 7TM family members do not cause general plant growth phenotypes compared to wild type.** **A.** RT-PCR of *7TM1* and *7TM5*, and *ACT2* as a control in wild type Col-0 compared to corresponding mutants. **B.** Phylogenetic assessment of full-length Arabidopsis and human protein sequences. Evolutionary analysis shows a bifurcation of the LUSTR family. Sequences were aligned using MUSCLE and the tree was constructed using Maximum Likelihood with 100 bootstrap replicates. Tree branch lengths are drawn to scale (substitutions per site). *A. thaliana* sequences: 7TM1 (At5g18520.1), 7TM2 (At3g09570.1), 7TM3 (At5g02630.1), 7TM4 (At5g42090.1), 7TM5 (At2g01070.1). *Homo sapiens* protein sequences: GPR107 (NP_001130029.1), GPR108 (NP_001073921.1), TMEM87A isoform 1 (NP_056312.2), TMEM87B isoform 1 (NP_116213.1). **C.** Quantification of hypocotyl lengths of 7-day-old etiolated seedlings grown on ½ MS media supplemented with 2 nM isoxaben. Box limits indicate 25th and 75th percentiles, whiskers extend to 1.5 times the interquartile range, median is indicated by a line, mean by a red "+", individual data points are shown, n is indicated in parentheses, and samples marked by an * are significantly different from wild type (one-way t-test, $p<0.05$). **D.** Representative images of 7-day-old etiolated seedlings grown on ½ MS media. **E.** Representative images of 4-week-old plants grown under unstressed, long-day conditions.

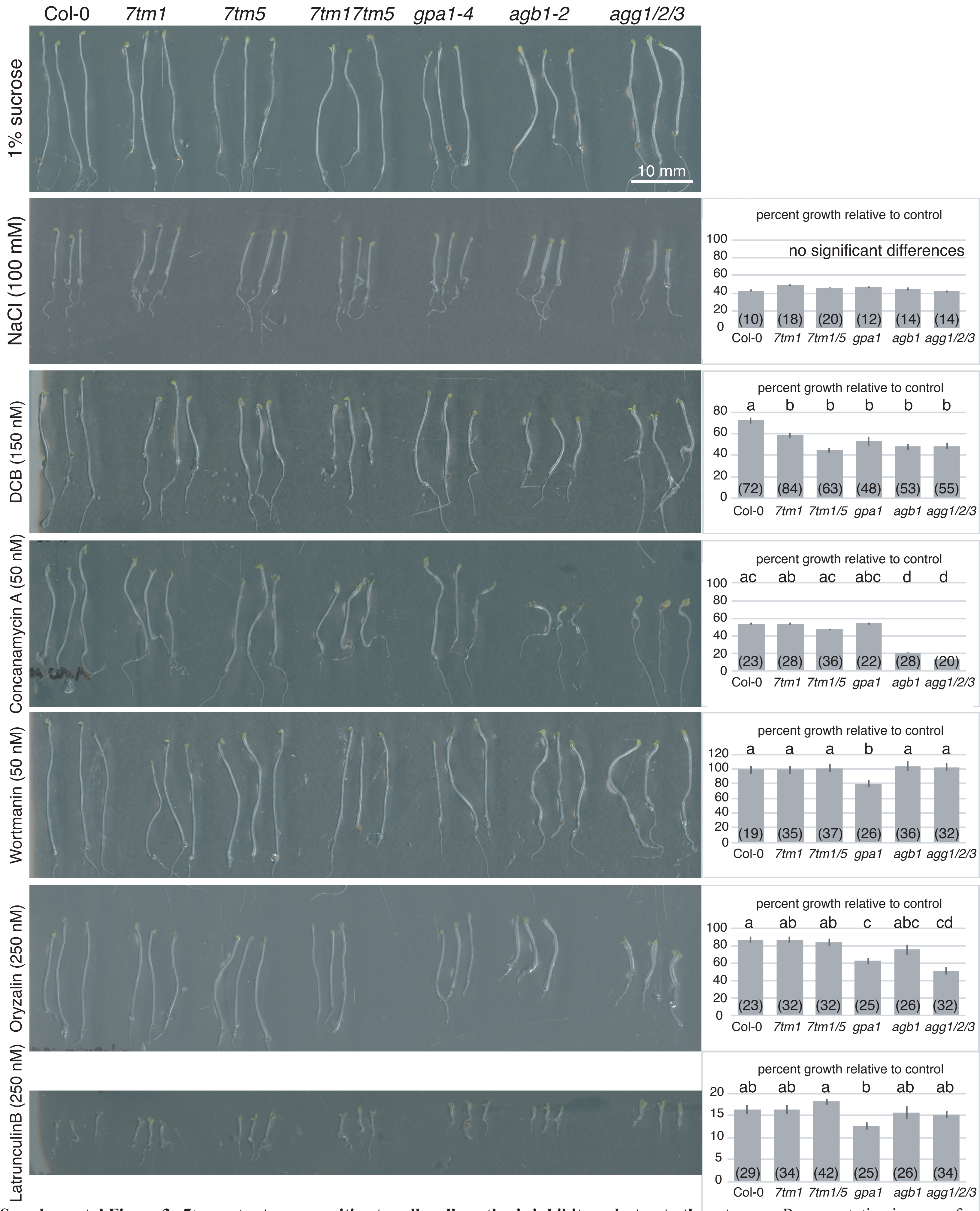


**Supplemental Figure 3: *7tm* mutants are sensitive to cell wall synthesis inhibitors, but not other stresses.** Representative images of 7-day-old etiolated seedlings grown on ½ MS media supplemented with inhibitors and quantification of hypocotyl lengths; n is indicated in parentheses, samples not sharing a common letter are significantly different (one-way ANOVA and Tukey HSD test, $p<0.05$).

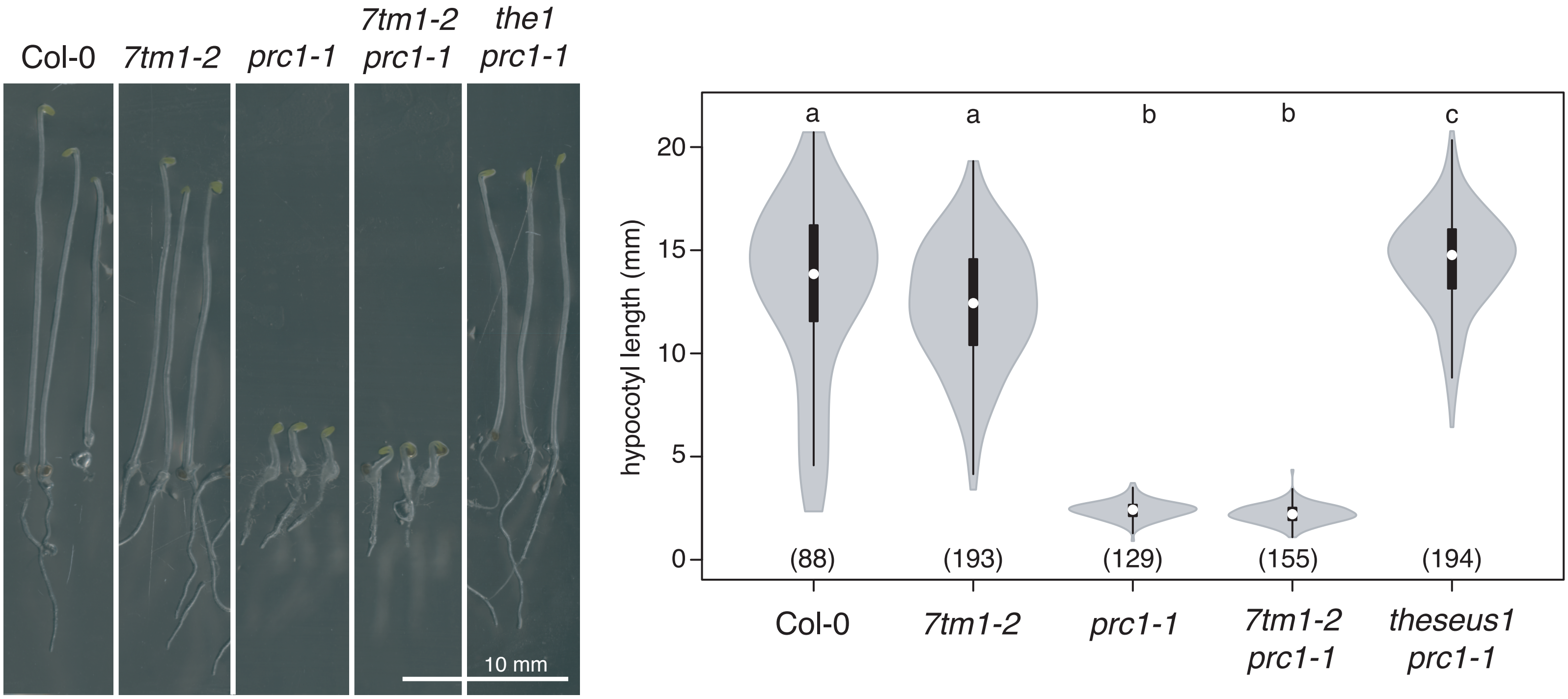


**Supplemental Figure 4: *7tm1 prc1-1* double mutants show slight decrease in growth as compared to *prc1-1*.** Representative images of 7-day-old etiolated seedlings grown on ½ MS media and quantification of etiolated hypocotyl lengths; n is indicated in parentheses, samples not sharing a common letter are significantly different (one-way ANOVA and Tukey HSD test, $p<0.05$), white circles show the medians, wide bar limits indicate the 25th and 75th percentiles, whiskers extend 1.5 times the interquartile range, polygons represent density estimates of data and extend to extreme values.

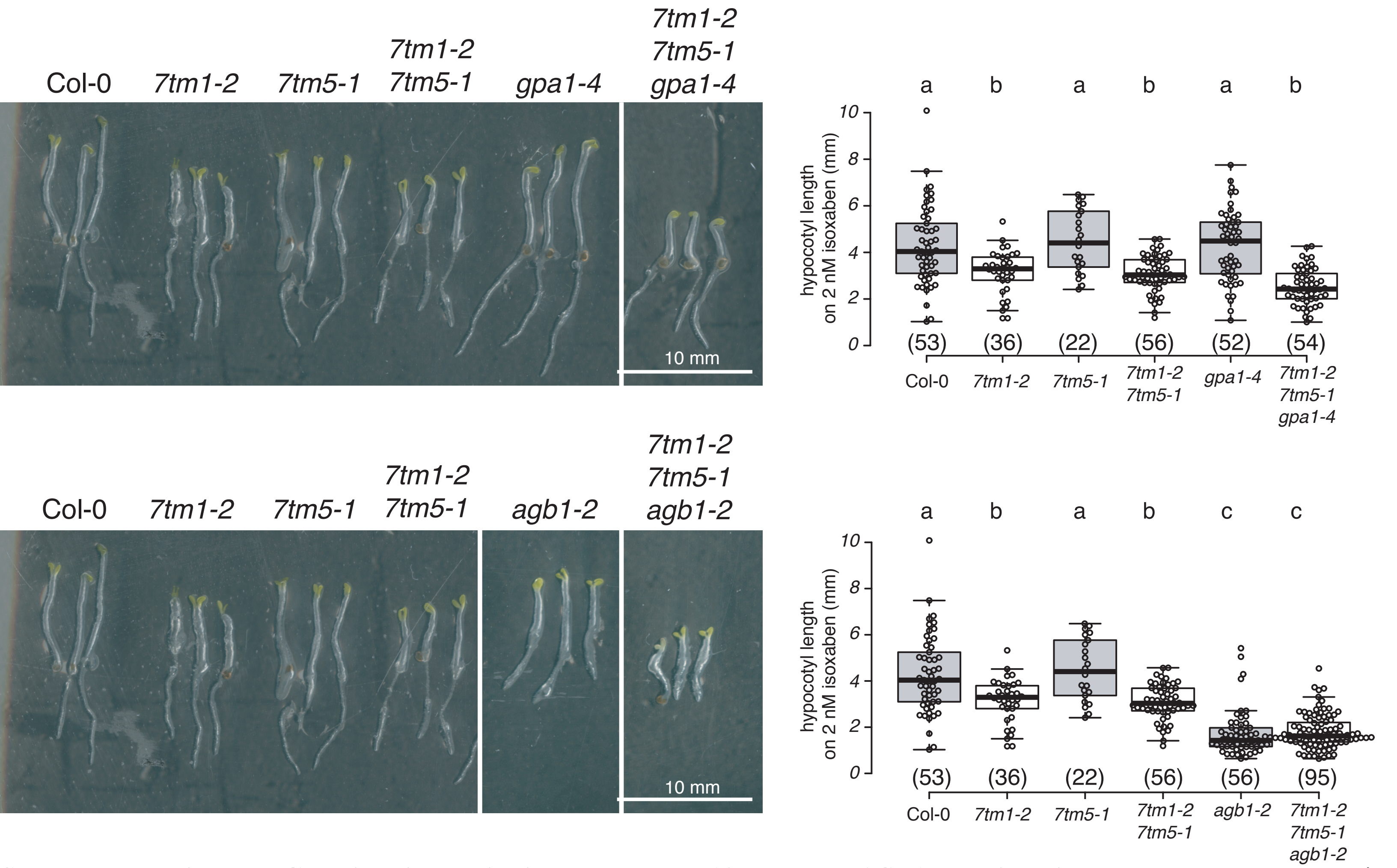


**Supplemental Figure 5: Genetic evidence indicates that *7TM1/7TM5* and *AGB1* work in a linear pathway.** Representative images of 7-day-old etiolated seedlings grown on ½ MS media supplemented with 2 nM isoxaben and quantification of etiolated hypocotyl lengths; n is indicated in parentheses, samples not sharing a common letter are significantly different (one-way ANOVA and Tukey HSD test, $p<0.05$), box limits indicate 25th and 75th percentiles, whiskers extend to 1.5 times the interquartile range, median is indicated by a line, and individual data points are shown. The same data for Col-0 wild type, *7tm1-2*, *7tm5-1* and *7tm1 7tm5* double mutants is shown in both graphs since these controls were grown together with the G protein and triple mutants.

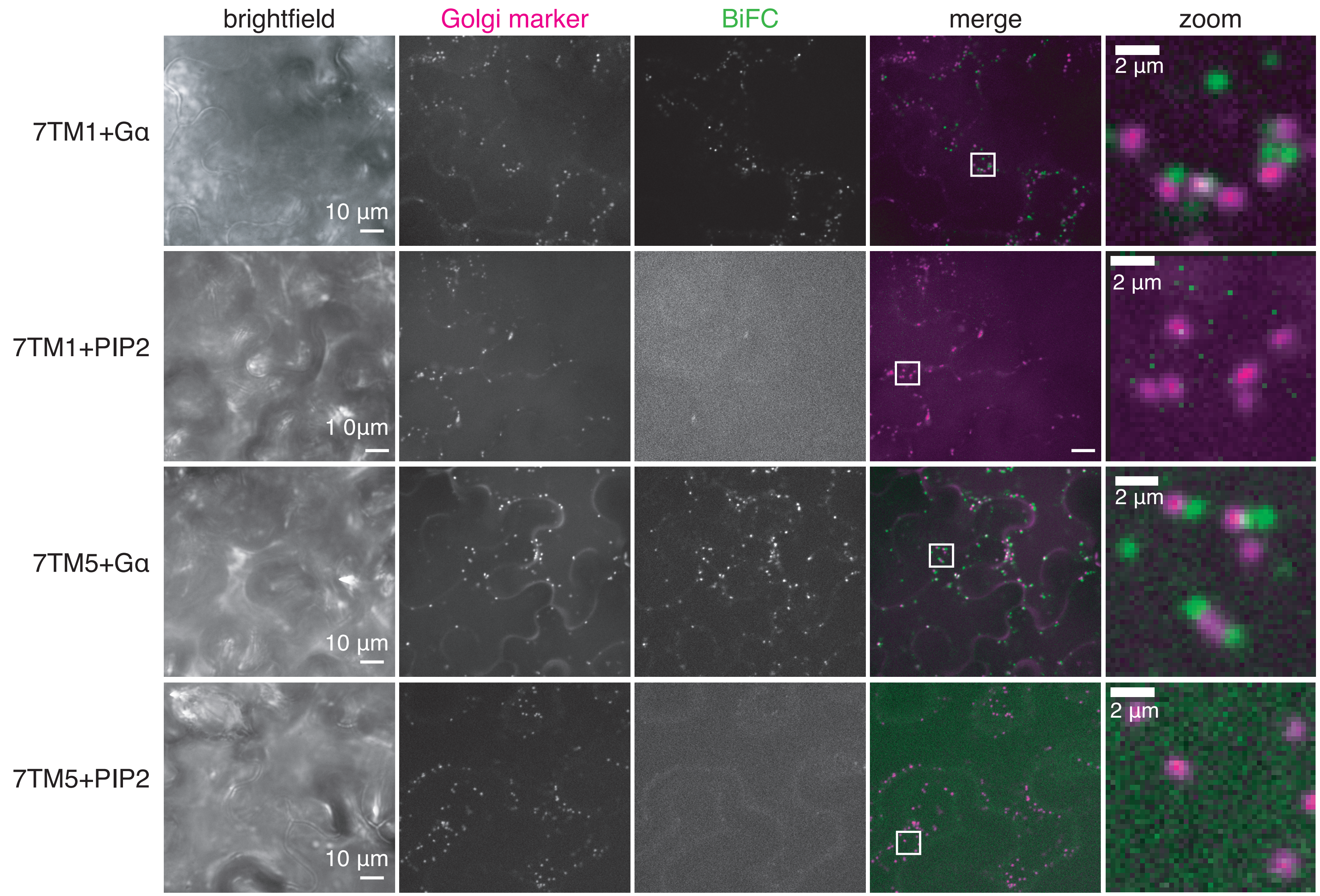


**Supplemental Figure 6: 7TMs and Gα interact in BiFC assays.** Bimolecular Fluorescence Complementation (BiFC) assay in transiently infiltrated Nicotiana benthamiana leaf epidermal cells. PIP2, which, similar to the G proteins primarily localizes to the plasma membrane, was substituted for Gα as a negative control. BiFC signal is shown in green and Golgi-marker in magenta in the merge and zoom images; white boxes in the merge images indicate the region in the zoom images.

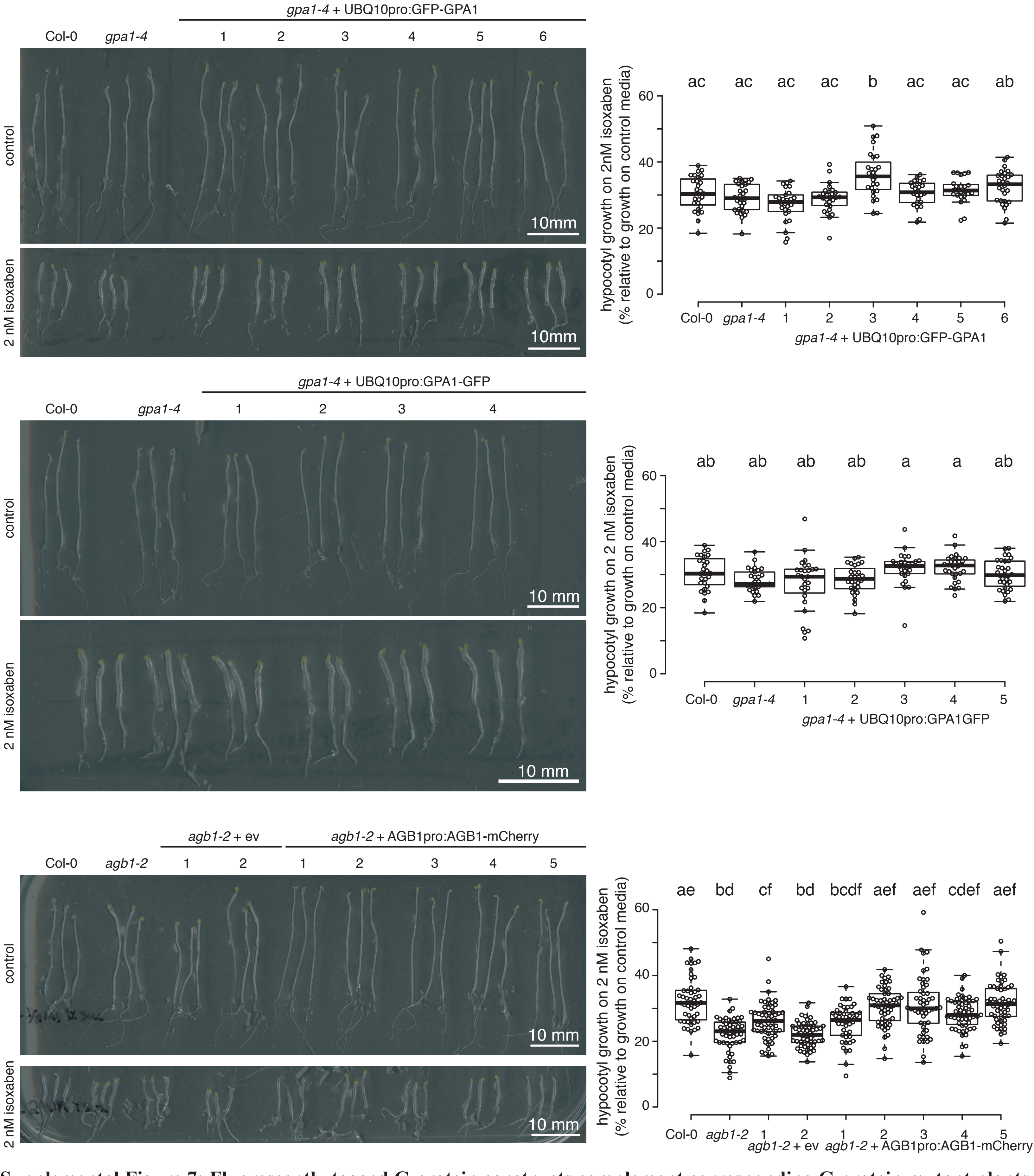


**Supplemental Figure 7: Fluorescently tagged G protein constructs complement corresponding G protein mutant plants.** Representative images of 7-day-old etiolated seedlings from homozygous, single insertion T3 lines (identified based on segregation of the selection marker) in the corresponding mutant background grown on ½ MS media supplemented with 2 nM isoxaben, and quantification of etiolated hypocotyl lengths. Independent transformants are separated, n is indicated in parentheses, samples not sharing a common letter are significantly different (one-way ANOVA and Tukey HSD test, $p<0.05$), box limits indicate 25th and 75th percentiles, whiskers extend to 1.5 times the interquartile range, median is indicated by a line, and individual data points are shown.

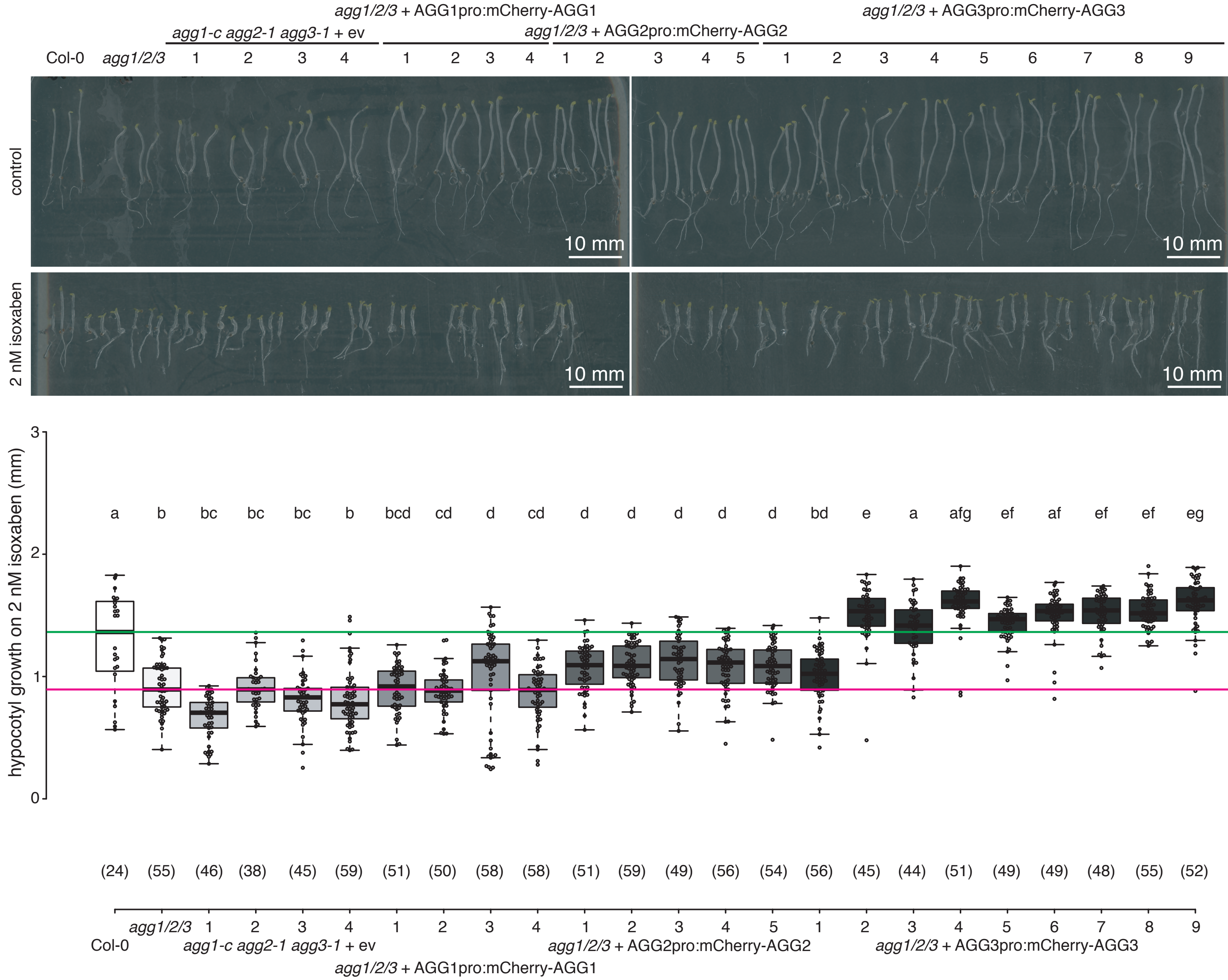


**Supplemental Figure 8: Fluorescently tagged G protein constructs complement corresponding G protein mutant plants.** Representative images of 7-day-old etiolated seedlings from homozygous, single insertion T3 lines (identified based on segregation of the selection marker) in the corresponding mutant background grown on ½ MS media supplemented with 2 nM isoxaben and quantification of etiolated hypocotyl lengths. Independent transformants are separated, n is indicated in parentheses, samples not sharing a common letter are significantly different (one-way ANOVA and Tukey HSD test, $p<0.05$), box limits indicate 25th and 75th percentiles, whiskers extend to 1.5 times the interquartile range, median is indicated by a line, and individual data points are shown, green line indicates wild type median, magenta line indicates *agg1-1c agg2-1 agg3-1* triple mutant median.

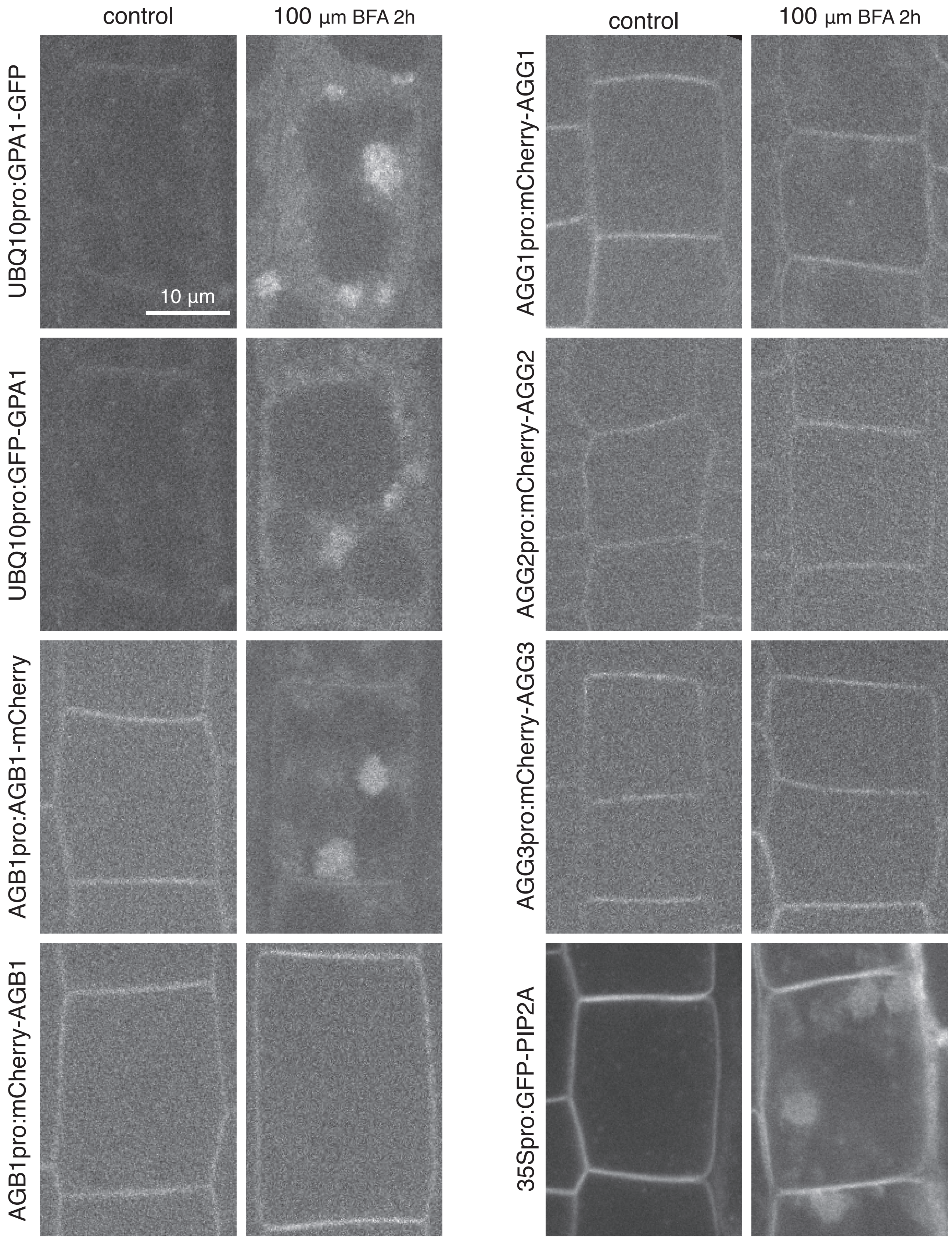


**Supplemental Figure 9: G protein components are localized to the plasma membrane and intracellular compartments.** Representative images from 3-day-old seedling root cells of GPA1-GFP, GFP-GPA1 (both Ubiquitin10 promoter-driven), AGB1-mCherry, mCherry-AGB1, mCherry-AGG1, mCherry-AGG2, mCherry-AGG3 (all native promoter-driven) single insertion T3 lines (identified based on segregation of the selection marker) in the corresponding mutant background under control conditions, or after 2 hour treatment with 100 μM brefeldin A (BFA).

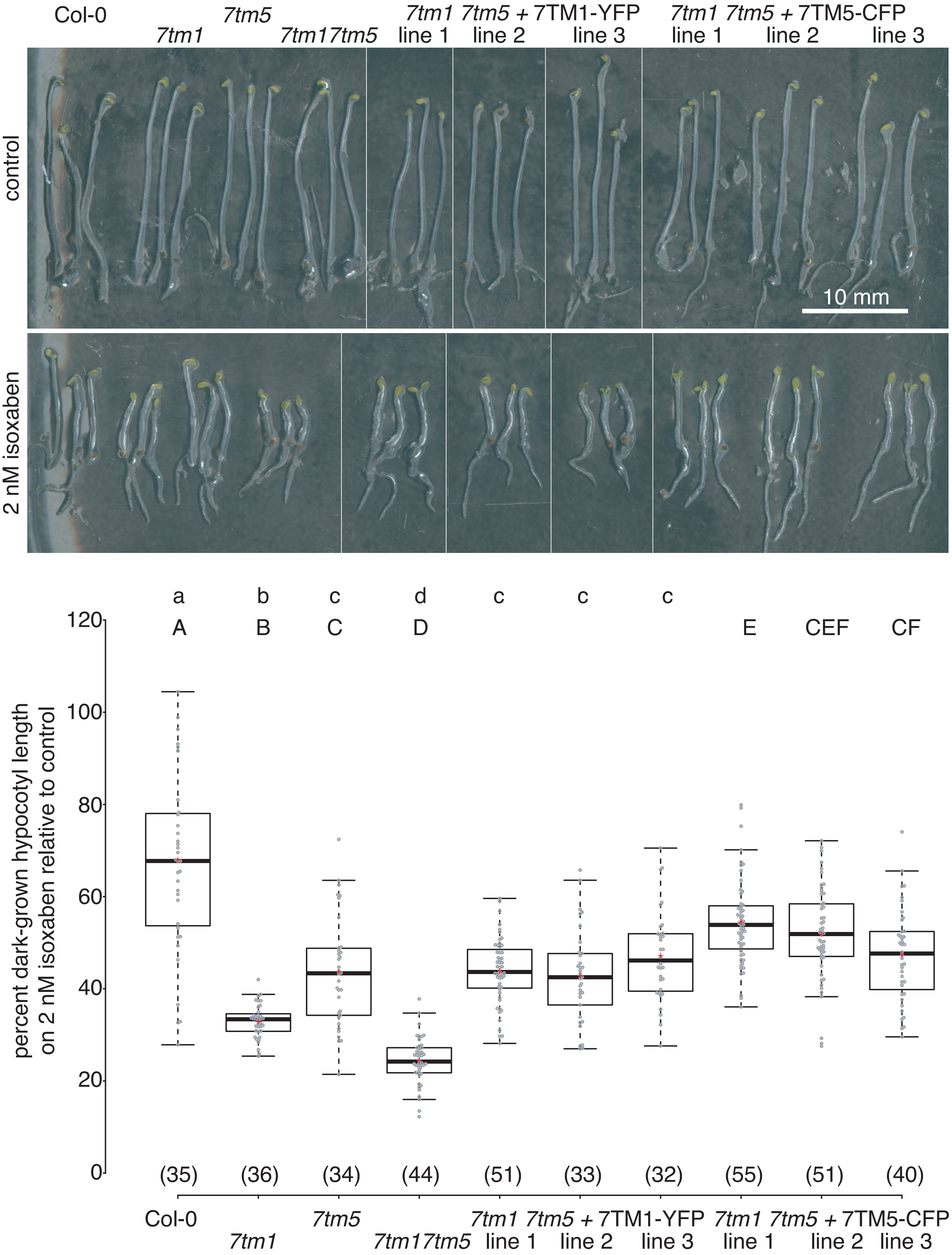


**Supplemental Figure 10: Fluorescently tagged 7TM constructs complement corresponding *7tm* mutant plants.** Representative images of 7-day-old etiolated seedlings from homozygous, single insertion T3 lines (identified based on segregation of the selection marker) in the corresponding mutant background grown on ½ MS media supplemented with 2 nM isoxaben and quantification of etiolated hypocotyl lengths. Independent transformants are separated, n is indicated in parentheses, samples not sharing a common letter (lowercase for 7TM1, uppercase for 7TM5) are significantly different (one-way ANOVA and Tukey HSD test, p<0.05), box limits indicate 25th and 75th percentiles, whiskers extend to 1.5 times the interquartile range, median is indicated by a line, mean by a red "+", and individual data points are shown.

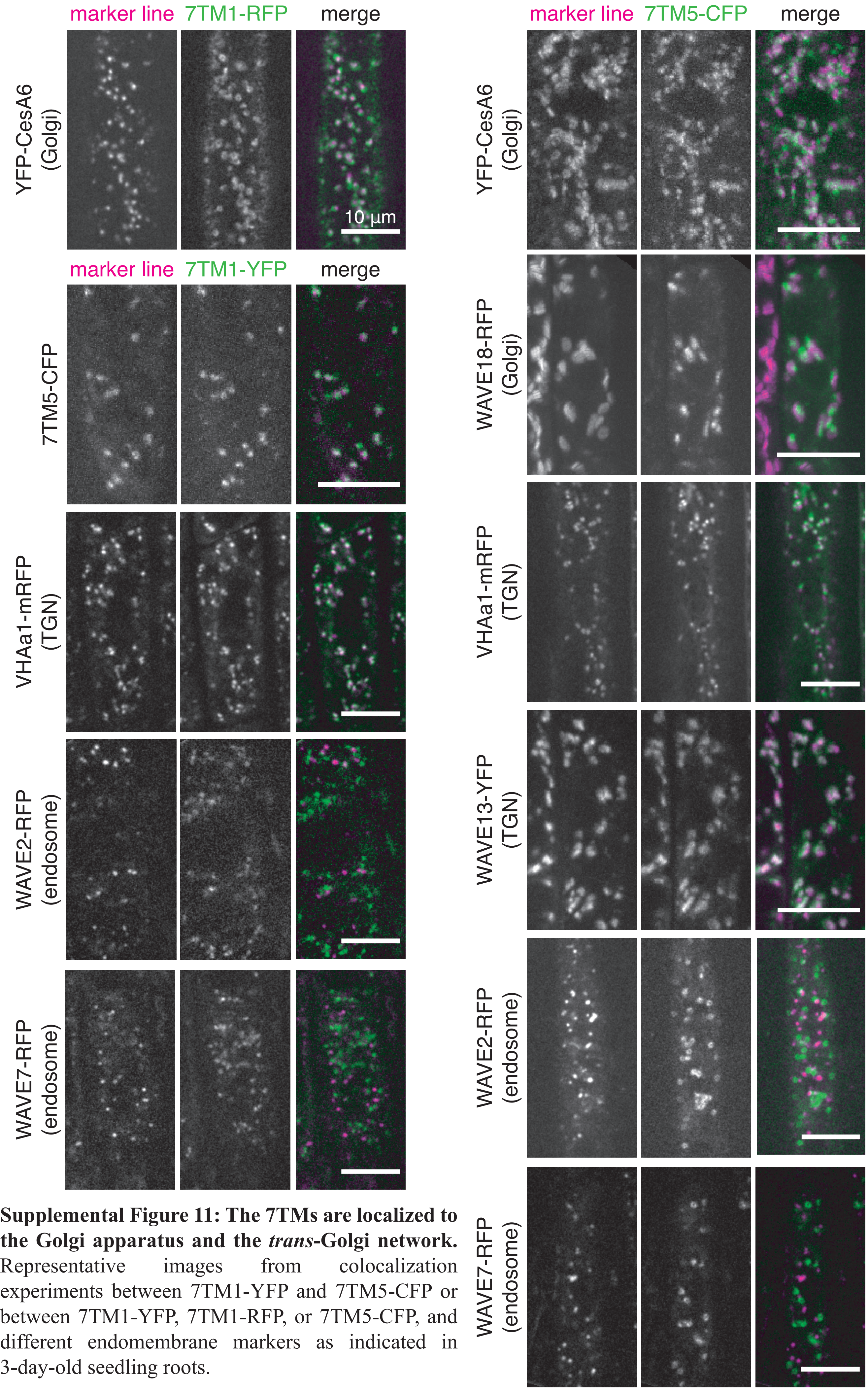


**Supplemental Figure 11: The 7TMs are localized to the Golgi apparatus and the *trans*-Golgi network.** Representative images from colocalization experiments between 7TM1-YFP and 7TM5-CFP or between 7TM1-YFP, 7TM1-RFP, or 7TM5-CFP, and different endomembrane markers as indicated in 3-day-old seedling roots.

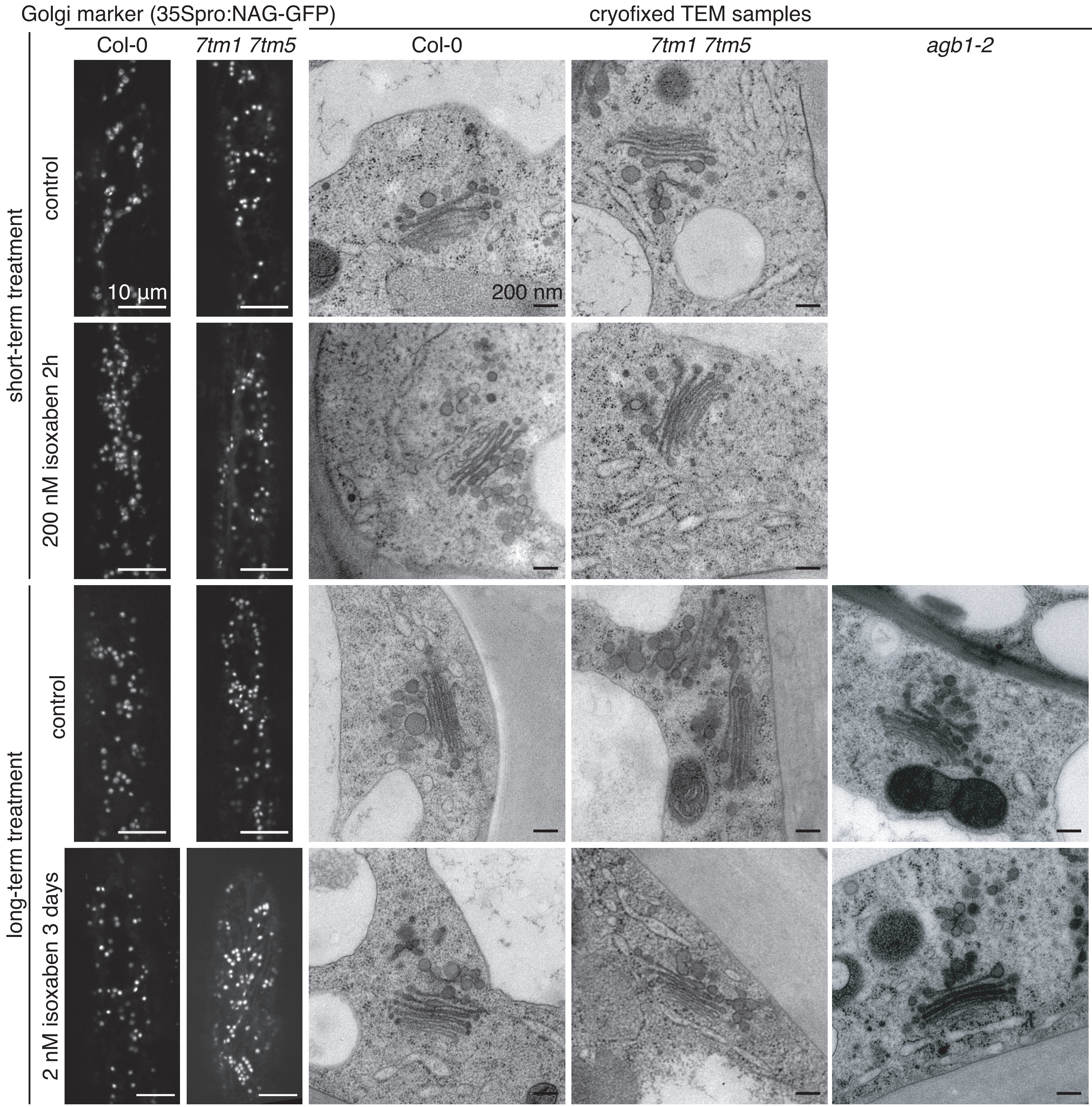


**Supplemental Figure 12: Golgi apparatus organization and structure are not affected *7tm1 7tm5* or *agb1* mutants.** Representative images of the NAG-GFP Golgi marker and of the Golgi apparatus in high-pressure frozen, freeze-substituted transmission electron microscopy samples from wild type, *7tm17tm5* double mutant and *agb1-2* single mutant 3-day-old etiolated hypocotyls under control conditions compared to short-term (200 nM for 2 hours) and long-term (2 nM for 3 days) isoxaben treatment.

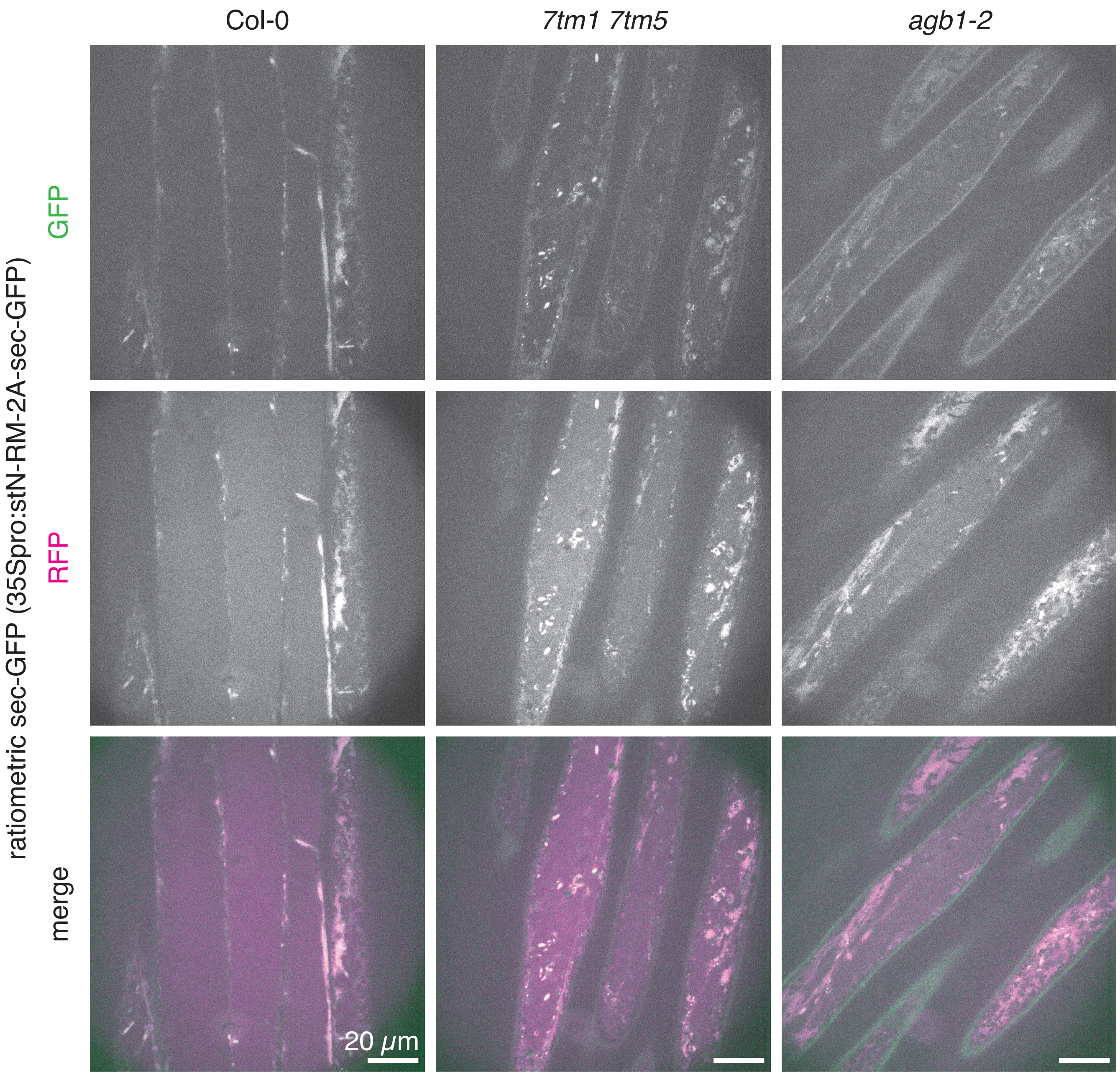


**Supplemental Figure 13: Localization of sec-GFP is not affected in *7tm1 7tm5* or *agb1* mutants.** Representative images of the ratiometric sec-GFP marker (35S:stN-RM-2A-sec-GFP) crossed into and *7tm1 7tm5* and *agb1-2* mutants, compared to wild type 3-day-old etiolated hypocotyls. Green channel represents sec-GFP signal as it is transported through the secretory pathway, magenta channel represents RFP that is expressed at the same level and targeted to the vacuole via the Golgi apparatus.

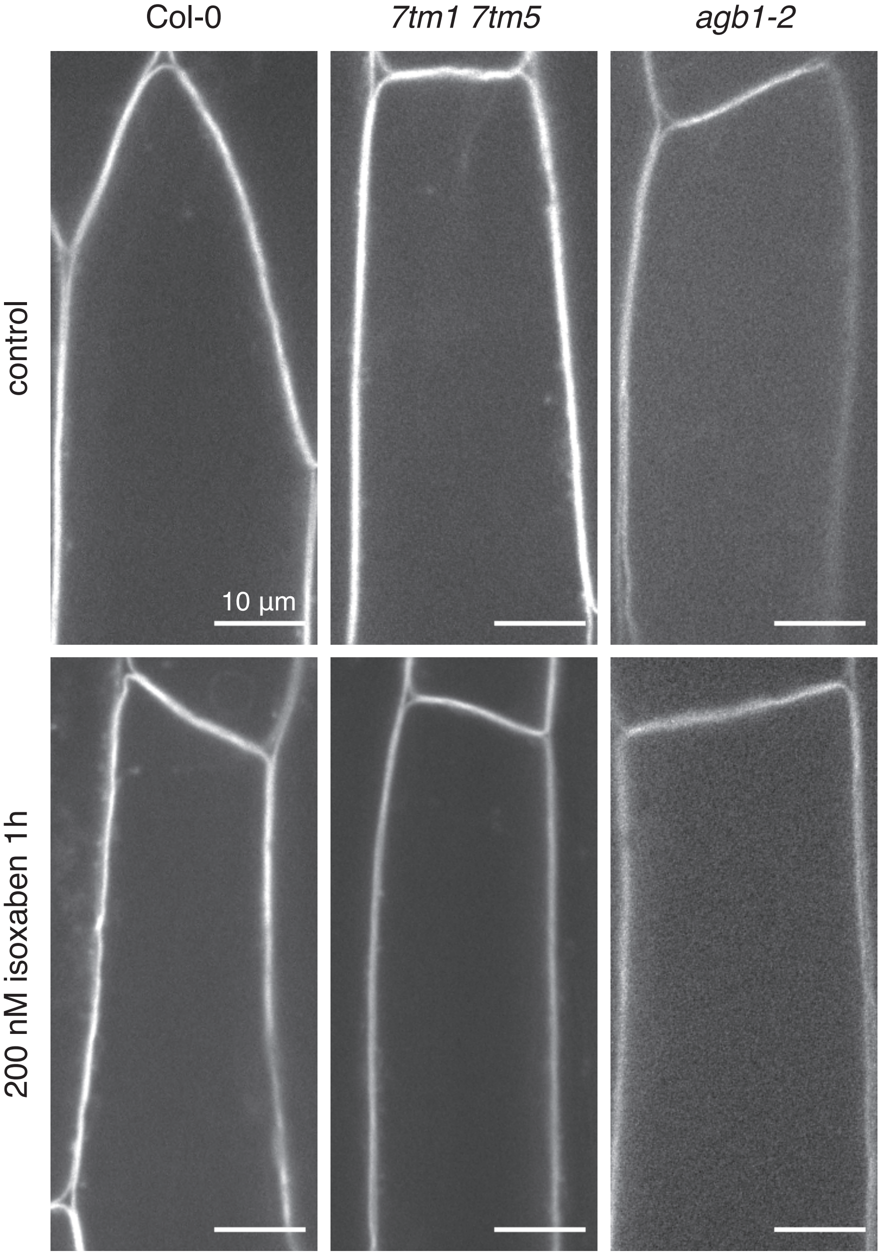


**Supplemental Figure 14: Localization of GFP-LTI6b is not affected in *7tm1 7tm5* or *agb1* mutants.** Additional representative images of the plasma membrane marker 35S:GFP-LTI6b in 3-day-old etiolated hypocotyl cells of *7tm1 7tm5* and *agb1-2* mutants, compared to wild type under control conditions or after short-term (200 nM for 1 hour) isoxaben treatment.

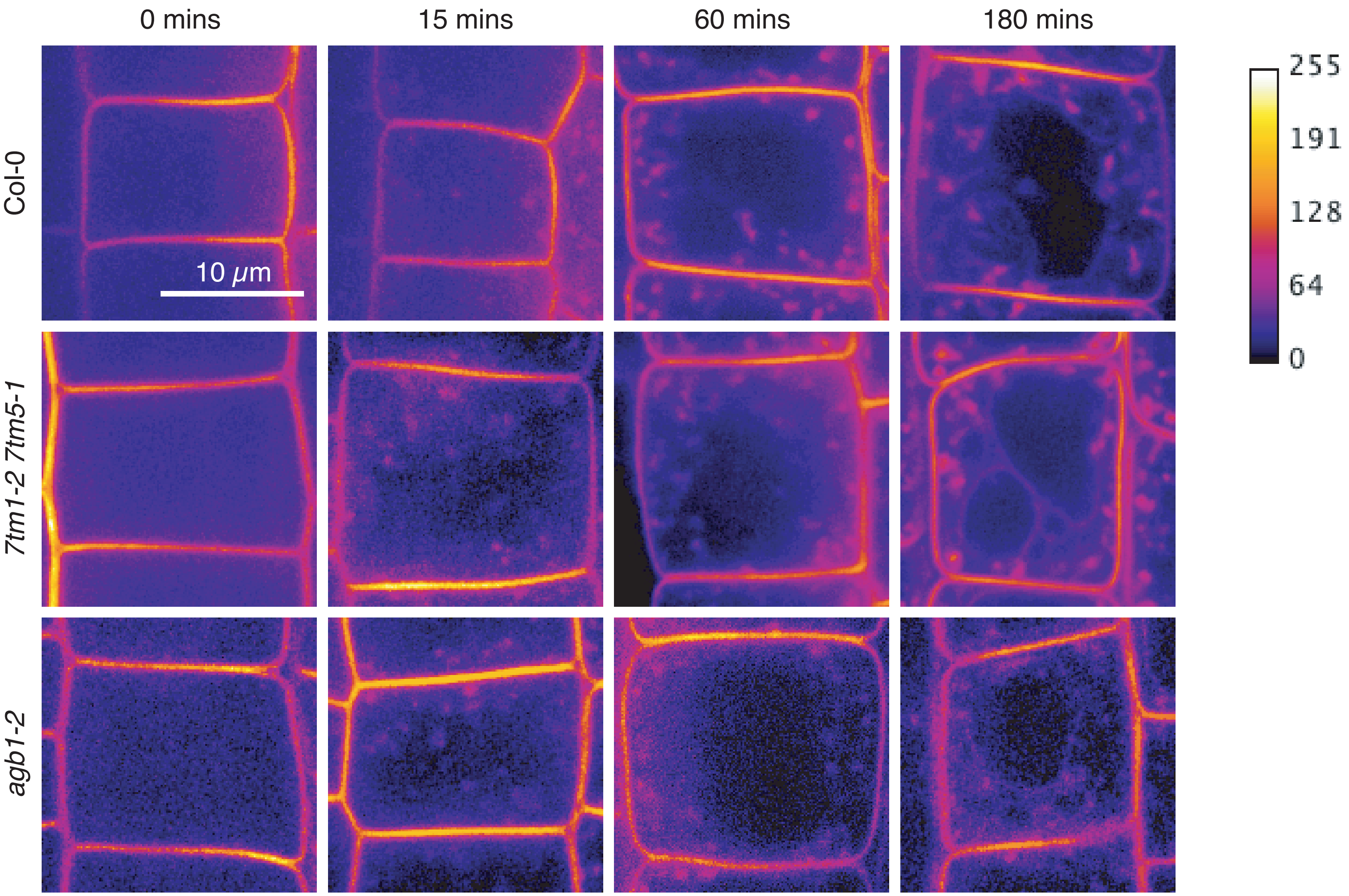


**Supplemental Figure 15: Time course of FM4-64 uptake reveals that uptake is not affected in *7tm1 7tm5* and *agb1* mutants.** Representative time course images of membrane dye (FM4-64) uptake into in root cells of 3-day-old seedlings of *7tm1 7tm5* and *agb1-2* mutants, compared to wild type. FM4-64 initially labels the plasma membrane, then travels to the early endosome, late endosome, and the tonoplast (vacuole membrane) over 3 hours. Images are false-coloured according to the scale

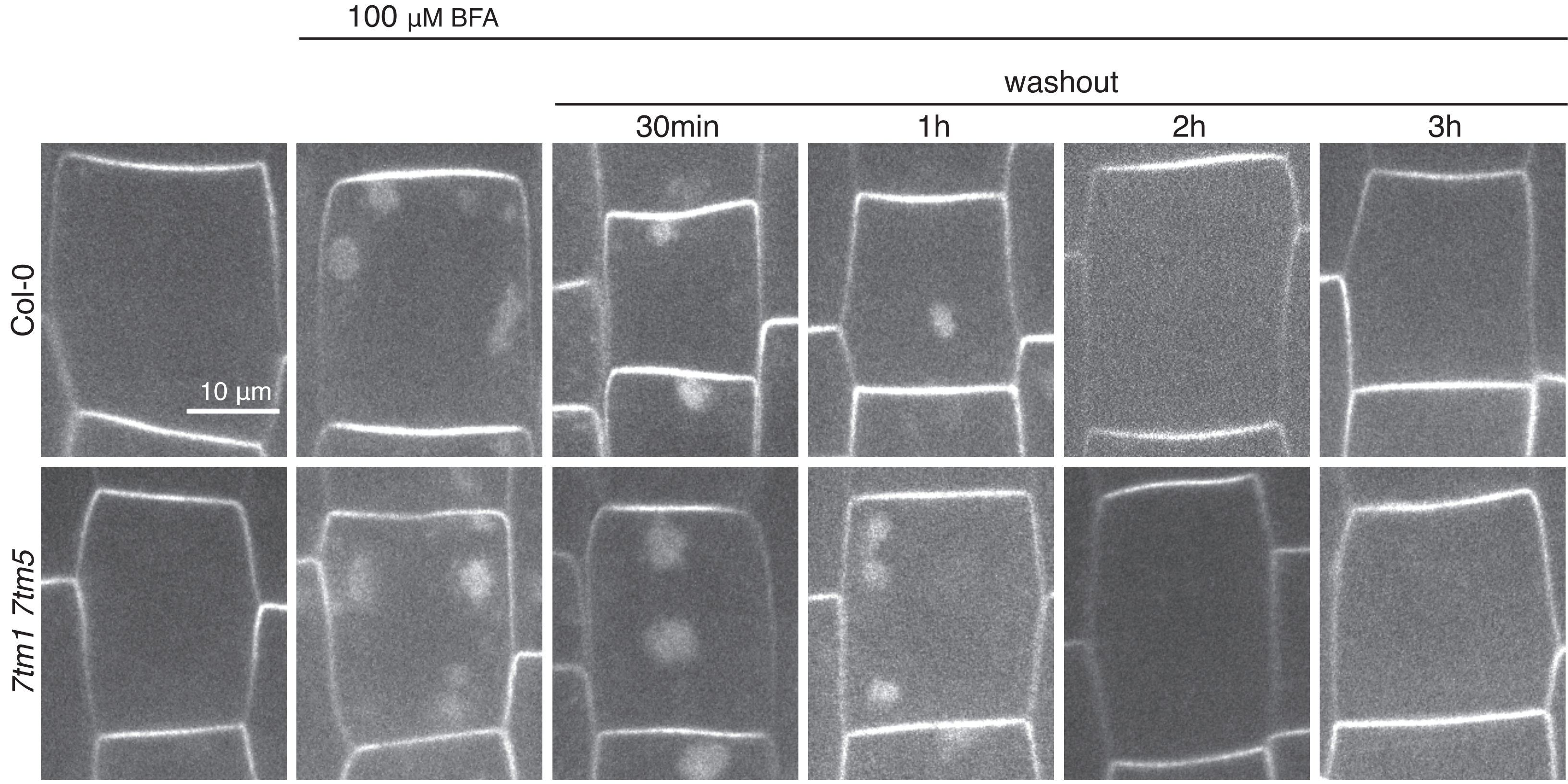


**Supplemental Figure 16: Time course of PIN2-GFP localization after BFA treatment and washout reveals that localization is not affected in *7tm1 7tm5* mutants.** Representative time course images of PIN2-GFP localization before and after 2 hour treatment with 100 µM brefeldin A (BFA), and after a time course of BFA washout in root cells of 3-day-old *7tm1 7tm5* mutant seedling roots , compared to wild type.

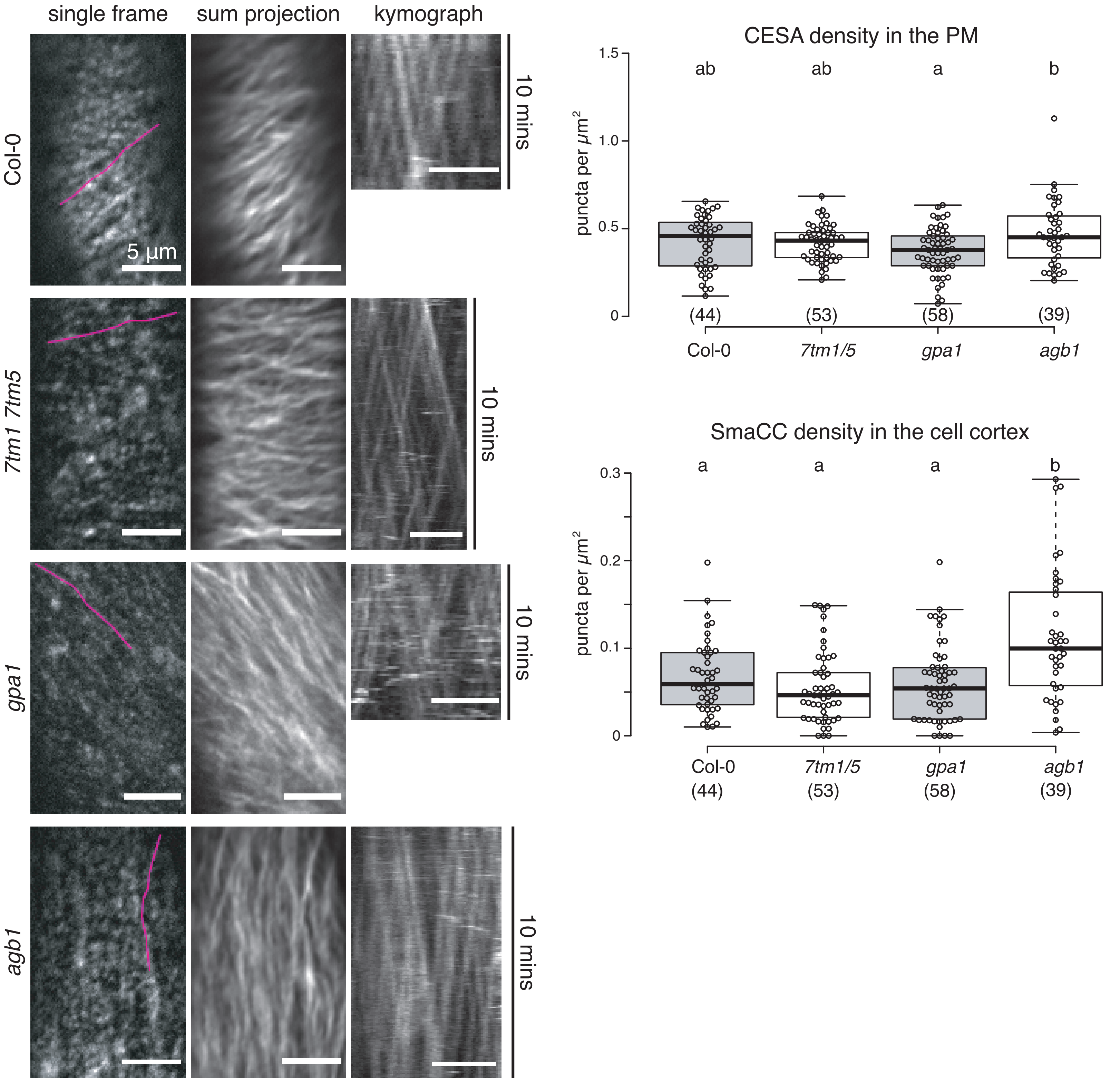


**Supplemental Figure 17: YFP-CESA6 movement and density at the plasma membrane are unaffected in *7tm 7tm5* and G protein mutants.** Representative images of YFP-CESA6 crossed into *7tm1 7tm5*, *gpa1-4* and *agb1-2* mutants, compared to wild type in in 3-day-old etiolated hypocotyl cells. Data are represented as single frames from the time lapse, a sum projection of the data (from 10 minute time lapse with 10 second intervals) and a kymograph from the magenta line indicated in the single frame. CESA speeds in the plasma membrane were quantified using the kymographs while CESA density at the plasma membrane and SmaCC density in the cortical cytoplasm were quantified from single frames. Graphs represent data from three independent experiments, n (cells, no more than three cells imaged per seedling) is indicated in parentheses, samples not sharing a common letter are significantly different (one-way ANOVA and Tukey HSD test, $p<0.05$), box limits indicate 25th and 75th percentiles, whiskers extend to 1.5 times the interquartile range, median is indicated by a line, and individual data points are shown.

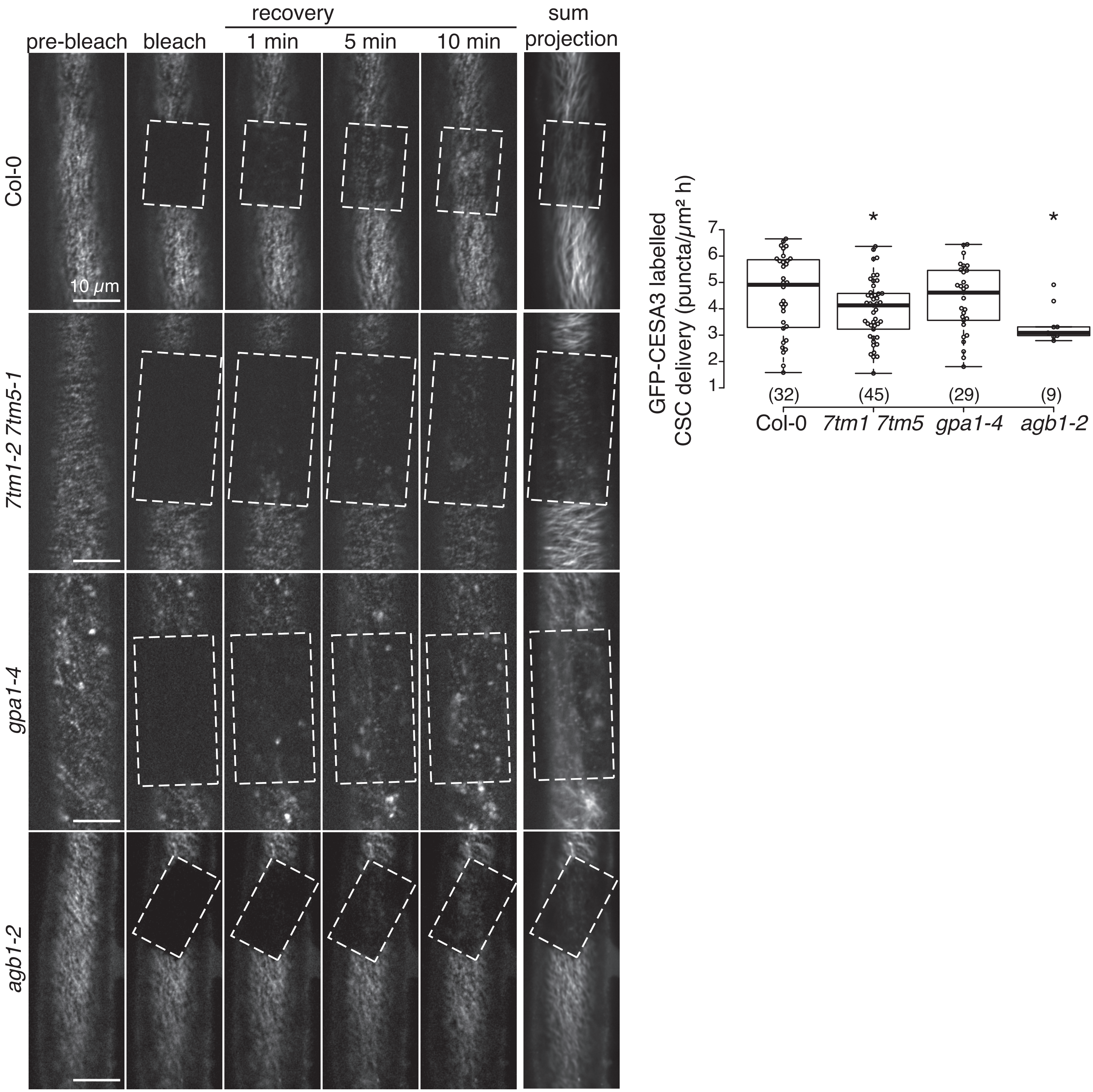


**Supplemental Figure 18: Photobleaching experiments reveal reduced GFP-CESA3 delivery to the plasma membrane in *7tm1 7tm5* and *agb1* mutants compared to wild type.** Representative images of GFP-CESA3 crossed into *7tm1 7tm5*, *gpa1-4* and *agb1-2* mutants, compared to wild type in 3-day-old etiolated hypocotyl cells. Bleached area is indicated by dashed box, scale bars represent 10 µm. Images are single frames from the time lapse or a sum projection of the data (from 10 minute time lapse with 10 second intervals), as indicated. CESA-labelled CSC particles secreted to the plasma membrane were quantified according to Sampthkumar et al. (2013). Graphs represent data from two (*gpa1-4* and *agb1-2*) or three (Col-0 and *7tm1 7tm5*) independent experiments, n (cells, no more than three cells imaged per seedling) is indicated in parentheses, box limits indicate 25th and 75th percentiles, whiskers extend to 1.5 times the interquartile range, median is indicated by a line, individual data points are shown, and samples marked by an * are significantly different from control (one-way t-test, $p<0.05$).

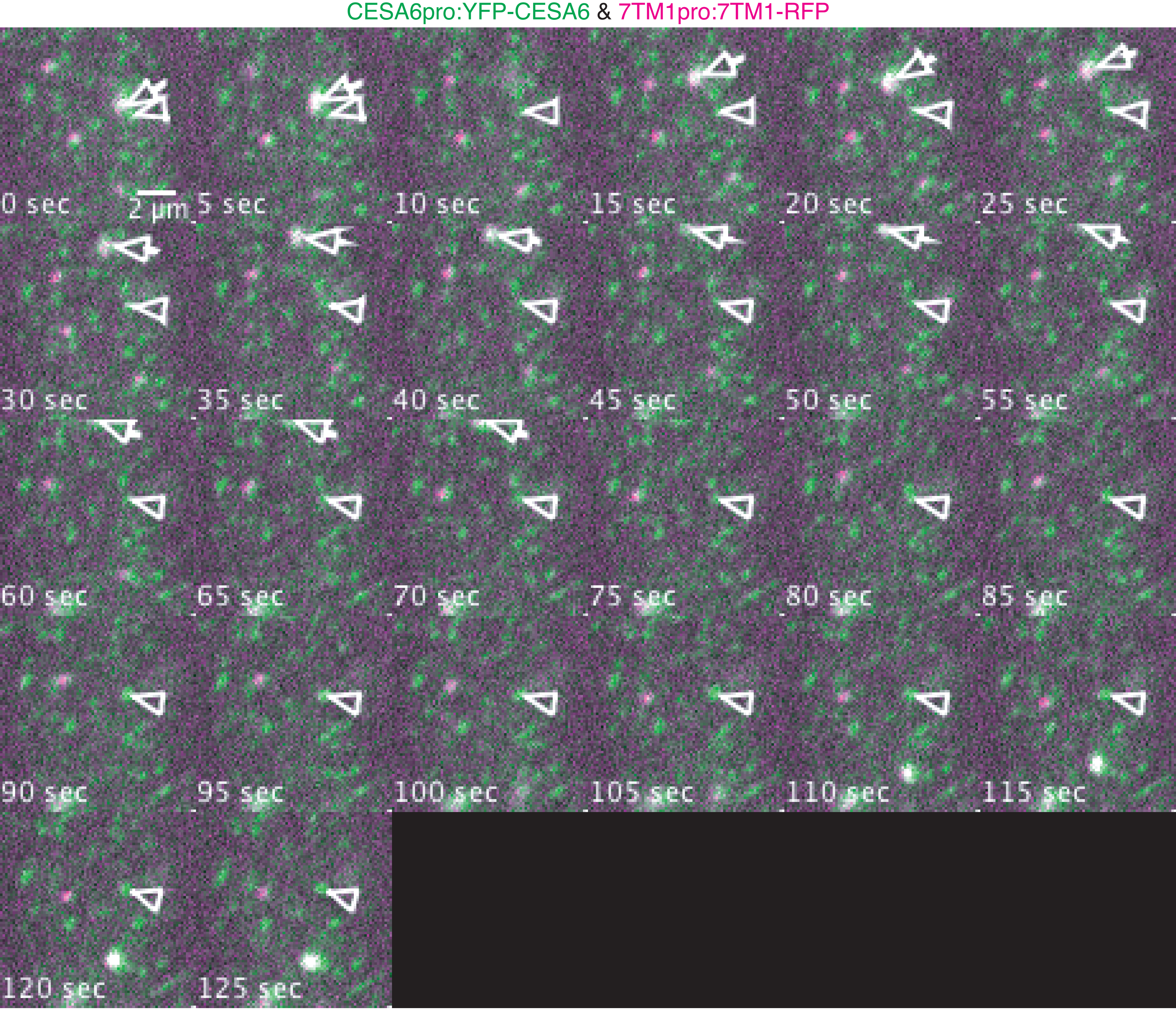


**Supplemental Figure 19: YFP-CESA6 and 7TM1-RFP co-localize in rapidly moving SmaCCs.** Montage image of YFP-CESA6 and 7TM1-RFP colocalization in wild type 3-day-old etiolated hypocotyl cells treated with 100 nM isoxaben for 1 hour. Arrow indicates a fast-moving SmaCC containing both YFP-CESA6 and 7TM1-RFP signal; arrowhead indicates a stalled SmaCC that contains only YFP-CEAA6. Scale bar represents 2 µm.

**Supplemental Table 1: Co-expression gene list using *7TM1* (At5g18520) as bait (from ATTED-II; Obayashi et al., 2008)**

| Co-expression rank | Locus | Alias | Function | Entrez Gene ID | Supportability | Ath-m | Ath-r At5g18520 |
|---|---|---|---|---|---|---|---|
| **0** | **At5g18520** | **Cand7 (Lung seven transmembrane receptor)** | **Lung seven transmembrane receptor family protein** | **831970** | **3** | **0** | **0** |
| 1 | At4g32390 | transporter | Nucleotide-sugar transporter family protein | 829374 | 3 | 1.2 | 8.7 |
| 2 | At5g35160 | Endomembrane 70 | Endomembrane protein 70 protein family | 833470 | 3 | 1.3 | 1.8 |
| 3 | At2g01970 | Endomembrane 70 | Endomembrane protein 70 protein family | 814728 | 3 | 4.2 | 9.7 |
| 4 | At4g12650 | Endomembrane 70 | Endomembrane protein 70 protein family | 826878 | 3 | 5 | 3.7 |
| 5 | At4g22580 | Exostosin | Exostosin family protein | 828354 | 3 | 6.5 | 27.9 |
| 6 | At3g56110 | PRA1.B1 | prenylated RAB acceptor 1.B1 | 824777 | 3 | 8 | 136 |
| 7 | At3g09570 | Lung seven transmembrane receptor | Lung seven transmembrane receptor family protein | 820114 | 3 | 9.1 | 16 |
| 8 | At3g51160 | MUR_1 | NAD(P)-binding Rossmann-fold superfamily protein | 824280 | 3 | 9.2 | 8.4 |
| 9 | At5g19980 | GONST4 | golgi nucleotide sugar transporter 4 | 832120 | 3 | 9.5 | 13.8 |
| 10 | At1g14670 | Endomembrane 70 | Endomembrane protein 70 protein family | 838029 | 3 | 10.2 | 9.5 |
| 11 | At2g16510 | ATPase | ATPase, F0/V0 complex, subunit C protein | 816150 | 3 | 11.7 | 183.5 |
| 12 | At5g25100 | Endomembrane 70 | Endomembrane protein 70 protein family | 832581 | 3 | 12.8 | 10 |
| 13 | At4g14160 | Sec23/Sec24 transport | Sec23/Sec24 protein transport family protein | 827055 | 3 | 14.6 | 579.2 |
| 14 | At5g10840 | EMP1 | Endomembrane protein 70 protein family | 830951 | 3 | 14.8 | 11.6 |
| 15 | At5g57460 | | | 835851 | 3 | 18.5 | 17.9 |
| 16 | At2g28520 | VHA-A1 | vacuolar proton ATPase A1 | 817400 | 3 | 21 | 49.2 |
| 17 | At4g01220 | MGP4 | Nucleotide-diphospho-sugar transferase family protein | 828028 | 3 | 22.5 | 159.1 |
| 18 | At1g64650 | Major facilitator | Major facilitator superfamily protein | 842773 | 3 | 23.4 | 90.7 |
| 19 | At1g70490 | ARFA1D | Ras-related small GTP-binding family protein | 843385 | 3 | 23.6 | 333.1 |
| 20 | At3g25540 | LAG1 | TRAM, LAG1 and CLN8 (TLC) lipid-sensing domain containing protein | 822140 | 3 | 23.7 | 853.1 |
| 21 | At5g14040 | PHT3;1 | phosphate transporter 3;1 | 831252 | 3 | 24.2 | 76.2 |
| 22 | At5g13890 | DUF716 | Family of unknown function (DUF716) | 831236 | 3 | 28.4 | 26.3 |
| 23 | At5g42090 | Lung seven transmembrane receptor | Lung seven transmembrane receptor family protein | 834214 | 3 | 28.4 | 487.9 |
| 24 | At1g61790 | transferase | Oligosaccharyltransferase complex/magnesium transporter family protein | 842476 | 3 | 28.9 | 76.5 |
| 25 | At5g15490 | UGD3 | UDP-glucose 6-dehydrogenase family protein | 831402 | 3 | 29.2 | 43.3 |
| 26 | At3g24160 | PMP | putative type 1 membrane protein | 822002 | 3 | 29.5 | 88.6 |
| 27 | At5g60570 | Galactose oxidase/kelch repeat | Galactose oxidase/kelch repeat superfamily protein | 836178 | 3 | 29.6 | 73.2 |

| | | | | | | | |
|---|---|---|---|---|---|---|---|
| 28 | At2g38360 | PRA1.B4 | prenylated RAB acceptor 1.B4 | 818416 | 3 | 29.9 | 16.9 |
| 29 | At2g36880 | MAT3 | methionine adenosyltransferase 3 | 818260 | 3 | 30.3 | 356.9 |
| 30 | At1g77610 | transporter | EamA-like transporter family protein | 844096 | 3 | 30.4 | 241.4 |
| 31 | At3g07330 | CSLC6 | Cellulose-synthase-like C6 | 819921 | 3 | 31.1 | 29.6 |
| 32 | At4g11380 | Adaptin | Adaptin family protein | 826741 | 3 | 31.2 | 97.8 |
| 33 | At4g16360 | 5'-AMP-activated kinase beta-2 subunit | 5'-AMP-activated protein kinase beta-2 subunit protein | 827331 | 3 | 31.2 | 2946.1 |
| 34 | At3g07950 | rhomboid | rhomboid protein-related | 819986 | 3 | 31.4 | 128.8 |
| 35 | At3g02350 | GAUT9 | galacturonosyltransferase 9 | 820474 | 3 | 31.6 | 63.7 |
| 36 | At3g29360 | UGD2 | UDP-glucose 6-dehydrogenase family protein | 822594 | 3 | 31.9 | 75.7 |
| 37 | At4g02500 | XXT2 | UDP-xylosyltransferase 2 | 827940 | 3 | 32 | 758.1 |
| 38 | At5g54830 | N-terminal | DOMON domain-containing protein / dopamine beta-monooxygenase N-terminal domain-containing protein | 835573 | 3 | 32.5 | 104.6 |
| 39 | At3g59360 | UTR6 | UDP-galactose transporter 6 | 825105 | 3 | 32.6 | 691.7 |
| 40 | At1g13900 | PAP2 | Purple acid phosphatases superfamily protein | 837949 | 3 | 33 | 66.9 |
| 41 | At5g61240 | LRR | Leucine-rich repeat (LRR) family protein | 836245 | 3 | 33.5 | 81.5 |
| 42 | At2g34250 | SecY transport | SecY protein transport family protein | 817986 | 3 | 34.6 | 45.9 |
| 43 | At2g25520 | transporter | Drug/metabolite transporter superfamily protein | 817090 | 3 | 35.5 | 61.7 |
| 44 | At2g36300 | Yip1 | Integral membrane Yip1 family protein | 818202 | 3 | 36.4 | 76.5 |
| 45 | At1g12840 | DET3 | vacuolar ATP synthase subunit C (VATC) / V-ATPase C subunit / vacuolar proton pump C subunit (DET3) | 837840 | 3 | 36.6 | 262.1 |
| 46 | At3g04080 | APY1 | apyrase 1 | 819563 | 3 | 36.8 | 281.5 |
| 47 | At2g29900 | PS2 | Presenilin-2 | 817540 | 3 | 36.9 | 115.2 |
| 48 | At5g39990 | transferase | Core-2/I-branching beta-1,6-N-acetylglucosaminyltransferase family protein | 833996 | 3 | 37.9 | 82.2 |
| 49 | At5g09810 | ACT7 | actin 7 | 830841 | 3 | 38.8 | 437.4 |
| 50 | At2g01070 | Lung seven transmembrane receptor | Lung seven transmembrane receptor family protein | 814637 | 3 | 39.8 | 1259.6 |
| 51 | At3g52760 | Yip1 | Integral membrane Yip1 family protein | 824442 | 2 | 40 | 562.2 |
| 52 | At1g70330 | ENT1,AT | equilibrative nucleotide transporter 1 | 843369 | 3 | 40.5 | 36.2 |
| 53 | At1g78300 | GRF2 | general regulatory factor 2 | 844165 | 3 | 41 | 29.4 |
| 54 | At3g23300 | transferase | S-adenosyl-L-methionine-dependent methyltransferases superfamily protein | 821910 | 3 | 41.2 | 25.9 |
| 55 | At1g78900 | VHA-A | vacuolar ATP synthase subunit A | 844228 | 3 | 42 | 374.1 |
| 56 | At2g24170 | Endomembrane 70 | Endomembrane protein 70 protein family | 816951 | 3 | 43.5 | 109.1 |
| 57 | At5g05820 | transporter | Nucleotide-sugar transporter family protein | 830467 | 3 | 43.6 | 31.9 |
| 58 | At5g14790 | ARM repeat | ARM repeat superfamily protein | 831331 | 3 | 43.8 | 119.3 |
| 59 | At1g10950 | TMN1 | transmembrane nine 1 | 837638 | 3 | 44.3 | 76.3 |

| | | | | | | | |
|---|---|---|---|---|---|---|---|
| 60 | At3g53520 | UXS1 | UDP-glucuronic acid decarboxylase 1 | 824520 | 3 | 44.3 | 65.1 |
| 61 | At2g41770 | DUF288 | Protein of unknown function (DUF288) | 818776 | 3 | 44.8 | 81 |
| 62 | At1g56330 | SAR1B | secretion-associated RAS 1B | 842086 | 3 | 46.8 | 373.3 |
| 63 | At3g01930 | Major facilitator | Major facilitator superfamily protein | 821218 | 2 | 47.9 | 2047.7 |
| 64 | At5g37310 | Endomembrane 70 | Endomembrane protein 70 protein family | 833705 | 3 | 48.6 | 602.9 |
| 65 | At1g05520 | Sec23/Sec24 transport | Sec23/Sec24 protein transport family protein | 837054 | 3 | 48.8 | 331.6 |
| 66 | At5g13390 | NEF1 | no exine formation 1 | 831180 | 3 | 48.9 | 435.5 |
| 67 | At4g27320 | PHOS34 | Adenine nucleotide alpha hydrolases-like superfamily protein | 828840 | 3 | 50.1 | 318.8 |
| 68 | At5g18280 | APY2 | apyrase 2 | 831946 | 3 | 51.8 | 78.8 |
| 69 | At5g44370 | PHT4;6 | phosphate transporter 4;6 | 834463 | 3 | 52.1 | 230 |
| 70 | At4g24330 | DUF1682 | Protein of unknown function (DUF1682) | 828536 | 3 | 52.7 | 23.6 |
| 71 | At5g55630 | TPK1 | Outward rectifying potassium channel protein | 835657 | 3 | 53.3 | 3430 |
| 72 | At3g13410 | | | 820541 | 3 | 53.8 | 125.5 |
| 73 | At3g58730 | VATPD | vacuolar ATP synthase subunit D (VATD) / V-ATPase D subunit / vacuolar proton pump D subunit (VATPD) | 825042 | 3 | 54.4 | 867.6 |
| 74 | At2g26680 | | | 817209 | 3 | 55.4 | 656.3 |
| 75 | At4g34490 | CAP1 | cyclase associated protein 1 | 829600 | 3 | 55.5 | 639 |
| 76 | At3g18060 | transducin | transducin family protein / WD-40 repeat family protein | 821331 | 3 | 57 | 268.4 |
| 77 | At2g39130 | transporter | Transmembrane amino acid transporter family protein | 818499 | 2 | 57.1 | 1500 |
| 78 | At4g38510 | VAB2 | ATPase, V1 complex, subunit B protein | 830008 | 3 | 58 | 276.8 |
| 79 | At4g38500 | DUF616 | Protein of unknown function (DUF616) | 830007 | 3 | 59.7 | 172.6 |
| 80 | At1g10630 | ARFA1F | ADP-ribosylation factor A1F | 837606 | 3 | 60 | 52.1 |
| 81 | At3g14790 | RHM3 | rhamnose biosynthesis 3 | 820707 | 3 | 60 | 35.2 |
| 82 | At5g38460 | transferase | ALG6, ALG8 glycosyltransferase family | 833834 | 3 | 60.8 | 127.7 |
| 83 | At5g11230 | transporter | Nucleotide-sugar transporter family protein | 830993 | 3 | 61.3 | 309.7 |
| 84 | At1g09020 | SNF4 | homolog of yeast sucrose nonfermenting 4 | 837423 | 2 | 61.4 | 218.7 |
| 85 | At4g32410 | RSW1 | cellulose synthase 1 | 829376 | 3 | 63.5 | 2221.1 |
| 86 | At3g44340 | CEF | clone eighty-four | 823559 | 3 | 64 | 190.8 |
| 87 | At1g25490 | REGA | ARM repeat superfamily protein | 839135 | 3 | 65.2 | 492.9 |
| 88 | At1g04820 | TUA4 | tubulin alpha-4 chain | 839405 | 3 | 66 | 356.3 |
| 89 | At1g09270 | IMPA-4 | importin alpha isoform 4 | 837448 | 3 | 67.2 | 158.6 |
| 90 | At4g30190 | PMA2 | H(+)-ATPase 2 | 829142 | 3 | 72 | 197.5 |
| 91 | At5g40830 | transferase | S-adenosyl-L-methionine-dependent methyltransferases superfamily protein | 834083 | 3 | 74.4 | 1727.7 |
| 92 | At1g27930 | DUF579 | Protein of unknown function (DUF579) | 839686 | 3 | 75.4 | 1263.4 |

| | | | | | | | |
|---|---|---|---|---|---|---|---|
| 93 | At1g30890 | HRF1 | Integral membrane HRF1 family protein | 839973 | 3 | 76.3 | 365.9 |
| 94 | At3g28720 | | | 822504 | 3 | 78.7 | 48.4 |
| 95 | At3g07810 | RNA-binding | RNA-binding (RRM/RBD/RNP motifs) family protein | 819972 | 3 | 78.8 | 58.5 |
| 96 | At5g53140 | Protein phosphatase 2C | Protein phosphatase 2C family protein | 835395 | 3 | 78.9 | 728.6 |
| 97 | At1g26850 | transferase | S-adenosyl-L-methionine-dependent methyltransferases superfamily protein | 839229 | 3 | 79.7 | 147.8 |
| 98 | At3g52930 | FBA8 | Aldolase superfamily protein | 824459 | 3 | 79.9 | 434.5 |
| 99 | At1g71070 | transferase | Core-2/I-branching beta-1,6-N-acetylglucosaminyltransferase family protein | 843447 | 3 | 80.2 | 205.9 |
| 100 | At4g29260 | HAD | HAD superfamily, subfamily IIIB acid phosphatase | 829047 | 2 | 80.7 | 3908.4 |
| 101 | At4g26940 | transferase | Galactosyltransferase family protein | 828801 | 3 | 83.6 | 376.5 |
| 102 | At5g02410 | ALG10 | DIE2/ALG10 family | 831764 | 3 | 83.9 | 420 |
| 103 | At4g32640 | Sec23/Sec24 transport | Sec23/Sec24 protein transport family protein | 829399 | 3 | 85 | 1106.5 |
| 104 | At2g46170 | Reticulon | Reticulon family protein | 819224 | 3 | 85.3 | 353.2 |
| 105 | At1g34020 | transporter | Nucleotide-sugar transporter family protein | 840299 | 3 | 86 | 292.1 |
| 106 | At3g21220 | MKK5 | MAP kinase kinase 5 | 821676 | 3 | 87 | 289.4 |
| 107 | At5g59160 | TOPP2 | type one serine/threonine protein phosphatase 2 | 836034 | 3 | 87.3 | 128.5 |
| 108 | At3g05280 | Yip1 | Integral membrane Yip1 family protein | 819692 | 3 | 87.6 | 75.2 |
| 109 | At2g30060 | PH | Pleckstrin homology (PH) domain superfamily protein | 817557 | 3 | 87.7 | 308.2 |
| 110 | At4g21150 | HAP6 | ribophorin II (RPN2) family protein | 827863 | 3 | 93.3 | 111.3 |
| 111 | At5g55610 | | | 835655 | 3 | 93.4 | 54 |
| 112 | At5g52180 | | | 835294 | 3 | 94.8 | 844.5 |
| 113 | At2g14835 | RING/U-box | RING/U-box superfamily protein | 815972 | 3 | 97.3 | 703.5 |
| 114 | At3g23660 | Sec23/Sec24 transport | Sec23/Sec24 protein transport family protein | 821946 | 3 | 98 | 206.8 |
| 115 | At5g01460 | LMBR1-like membrane | LMBR1-like membrane protein | 831815 | 3 | 98 | 242.1 |
| 116 | At1g29790 | transferase | S-adenosyl-L-methionine-dependent methyltransferases superfamily protein | 839857 | 3 | 98.1 | 788.8 |
| 117 | At1g12850 | mutase | Phosphoglycerate mutase family protein | 837842 | 3 | 99.2 | 279.6 |
| 118 | At5g59290 | UXS3 | UDP-glucuronic acid decarboxylase 3 | 836047 | 3 | 100.1 | 599.4 |
| 119 | At4g04910 | NSF | AAA-type ATPase family protein | 825830 | 3 | 102.9 | 118.5 |
| 120 | At4g39840 | | | 830143 | 3 | 103.2 | 182.1 |
| 121 | At3g46830 | RABA2c | RAB GTPase homolog A2C | 823836 | 3 | 105.2 | 371.9 |
| 122 | At4g00110 | GAE3 | UDP-D-glucuronate 4-epimerase 3 | 828145 | 2 | 106.8 | 114.1 |
| 123 | At3g19980 | STPP | flower-specific, phytochrome-associated protein phosphatase 3 | 821536 | 2 | 107.5 | 894.5 |
| 124 | At2g47170 | ARF1A1C | Ras-related small GTP-binding family protein | 819330 | 3 | 108 | 360.1 |
| 125 | At3g01360 | DUF716 | Family of unknown function (DUF716) | 821158 | 3 | 108.3 | 595.6 |

| | | | | | | | |
|---|---|---|---|---|---|---|---|
| 126 | At1g27190 | kinase | Leucine-rich repeat protein kinase family protein | 839608 | 3 | 108.3 | 5403.3 |
| 127 | At1g63000 | UER1 | nucleotide-rhamnose synthase/epimerase-reductase | 842603 | 3 | 108.5 | 434 |
| 128 | At4g36890 | IRX14 | Nucleotide-diphospho-sugar transferases superfamily protein | 829842 | 3 | 109.4 | 148 |
| 129 | At4g34720 | VHA-C1 | ATPase, F0/V0 complex, subunit C protein | 829624 | 3 | 109.6 | 1018.2 |
| 130 | At3g10370 | SDP6 | FAD-dependent oxidoreductase family protein | 820199 | 3 | 111.8 | 1037.4 |
| 131 | At1g62020 | Coatomer, | Coatomer, alpha subunit | 842497 | 3 | 112 | 70.8 |
| 132 | At3g55360 | TSC13 | 3-oxo-5-alpha-steroid 4-dehydrogenase family protein | 824702 | 3 | 112.1 | 1282.3 |
| 133 | At3g08580 | AAC1 | ADP/ATP carrier 1 | 820005 | 3 | 112.1 | 142.8 |
| 134 | At2g21600 | RER1B | endoplasmatic reticulum retrieval protein 1B | 816698 | 2 | 113.5 | 310.1 |
| 135 | At2g35610 | XEG113 | xyloglucanase 113 | 818128 | 3 | 113.5 | 39.7 |
| 136 | At5g44860 | | | 834516 | 2 | 114.1 | 167.2 |
| 137 | At4g08960 | PTPA | phosphotyrosyl phosphatase activator (PTPA) family protein | 826474 | 3 | 114.5 | 209.2 |
| 138 | At1g09330 | | | 837454 | 3 | 115.7 | 207.7 |
| 139 | At2g30050 | transducin | transducin family protein / WD-40 repeat family protein | 817556 | 3 | 116.4 | 145.1 |
| 140 | At5g19130 | Gaa1-like | GPI transamidase component family protein / Gaa1-like family protein | 832033 | 3 | 116.8 | 674.5 |
| 141 | At3g16760 | Tetratricopeptide repeat (TPR)-like | Tetratricopeptide repeat (TPR)-like superfamily protein | 820928 | 3 | 116.9 | 20.1 |
| 142 | At5g03080 | LPPgamma | Phosphatidic acid phosphatase (PAP2) family protein | 831801 | 3 | 117.3 | 1132.4 |
| 143 | At5g19690 | STT3A | staurosporin and temperature sensitive 3-like A | 832089 | 3 | 118.4 | 56.3 |
| 144 | At3g03250 | UGP1 | UDP-GLUCOSE PYROPHOSPHORYLASE 1 | 821313 | 3 | 118.4 | 893.7 |
| 145 | At5g21070 | | | 832231 | 3 | 118.8 | 485.3 |
| 146 | At1g18150 | MPK8 | Protein kinase superfamily protein | 838394 | 3 | 121 | 429.2 |
| 147 | At3g08930 | LMBR1-like membrane | LMBR1-like membrane protein | 820042 | 3 | 123.2 | 841.4 |
| 148 | At4g00740 | QUA3 | S-adenosyl-L-methionine-dependent methyltransferases superfamily protein | 828020 | 3 | 123.7 | 543.8 |
| 149 | At3g46290 | HERK1 | hercules receptor kinase 1 | 823774 | 3 | 124 | 156.2 |
| 150 | At4g35260 | IDH1 | isocitrate dehydrogenase 1 | 829679 | 3 | 125.4 | 130.1 |
| 151 | At1g08960 | CCX5 | cation exchanger 11 | 837416 | 3 | 126.1 | 77.9 |
| 152 | At4g14960 | TUA6 | Tubulin/FtsZ family protein | 827154 | 3 | 127.4 | 377.6 |
| 153 | At2g32380 | Transmembrane 97 | Transmembrane protein 97, predicted | 817799 | 3 | 129.3 | 282.9 |
| 154 | At1g20050 | MAD4 | C-8,7 sterol isomerase | 838593 | 3 | 129.4 | 1394.6 |
| 155 | At5g42420 | transporter | Nucleotide-sugar transporter family protein | 834248 | 3 | 131.4 | 1285.7 |
| 156 | At3g25040 | ERD2B | endoplasmic reticulum retention defective 2B | 822095 | 3 | 131.6 | 625.1 |
| 157 | At2g46505 | SDH4 | succinate dehydrogenase subunit 4 | 819261 | 3 | 131.8 | 220.4 |

| | | | | | | | |
|---|---|---|---|---|---|---|---|
| 158 | At5g53480 | ARM repeat | ARM repeat superfamily protein | 835429 | 2 | 133.1 | 415.1 |
| 159 | At5g05170 | MRE1 | Cellulose synthase family protein | 830399 | 3 | 133.5 | 795.3 |
| 160 | At1g18580 | GAUT11 | galacturonosyltransferase 11 | 838439 | 2 | 133.9 | 111.6 |
| 161 | At1g67930 | Golgi transport complex | Golgi transport complex protein-related | 843121 | 3 | 136.2 | 32.4 |
| 162 | At1g16560 | Per1-like | Per1-like family protein | 838226 | 3 | 137 | 519.4 |
| 163 | At1g15710 | dehydrogenase | prephenate dehydrogenase family protein | 838140 | 3 | 137.2 | 2275.3 |
| 164 | At4g23640 | TRH1 | Potassium transporter family protein | 828464 | 3 | 138.1 | 215.1 |
| 165 | At4g38920 | VHA-C3 | vacuolar-type H(+)-ATPase C3 | 830047 | 3 | 140.8 | 713.1 |
| 166 | At1g74380 | XXT5 | xyloglucan xylosyltransferase 5 | 843779 | 3 | 140.8 | 204 |
| 167 | At1g41830 | SKS6 | SKU5-similar 6 | 840792 | 3 | 141.2 | 4191.8 |
| 168 | At2g37550 | ASP1 | ARF-GAP domain 7 | 818331 | 3 | 141.7 | 164.1 |
| 169 | At5g39320 | UDG4 | UDP-glucose 6-dehydrogenase family protein | 833928 | 3 | 143.1 | 126.9 |
| 170 | At3g04090 | SIP1A | small and basic intrinsic protein 1A | 819564 | 3 | 143.7 | 212.1 |
| 171 | At5g64740 | PRC1 | cellulose synthase 6 | 836595 | 3 | 144.9 | 792.3 |
| 172 | At3g11800 | | | 820353 | 3 | 150.9 | 251 |
| 173 | At5g39500 | GNL1 | GNOM-like 1 | 833946 | 3 | 151 | 124.6 |
| 174 | At5g38480 | RCI1 | general regulatory factor 3 | 833836 | 3 | 154.8 | 1325.1 |
| 175 | At5g58060 | YKT61 | SNARE-like superfamily protein | 835918 | 3 | 156.2 | 653.8 |
| 176 | At5g36290 | UPF0016 | Uncharacterized protein family (UPF0016) | 833627 | 3 | 156.3 | 341.5 |
| 177 | At5g60920 | COB | COBRA-like extracellular glycosyl-phosphatidyl inositol-anchored protein family | 836213 | 3 | 157.5 | 357.6 |
| 178 | At3g17390 | SAMS3 | S-adenosylmethionine synthetase family protein | 821003 | 3 | 158.6 | 915.3 |
| 179 | At4g12110 | SMO1-1 | sterol-4alpha-methyl oxidase 1-1 | 826819 | 2 | 159 | 3867.3 |
| 180 | At2g03120 | SPP | signal peptide peptidase | 814841 | 3 | 162.7 | 194.8 |
| 181 | At4g39820 | Tetratricopeptide repeat (TPR)-like | Tetratricopeptide repeat (TPR)-like superfamily protein | 830141 | 3 | 163.4 | 35.6 |
| 182 | At1g67730 | YBR159 | beta-ketoacyl reductase 1 | 843098 | 3 | 164.9 | 402.2 |
| 183 | At5g63220 | | | 836442 | 3 | 165.2 | 481 |
| 184 | At1g50010 | TUA2 | tubulin alpha-2 chain | 841425 | 3 | 166.9 | 621.3 |
| 185 | At5g61780 | Tudor2 | TUDOR-SN protein 2 | 836300 | 3 | 168 | 204.5 |
| 186 | At4g00660 | RH8 | RNAhelicase-like 8 | 828042 | 3 | 168.5 | 595.8 |
| 187 | At1g75500 | WAT1 | Walls Are Thin 1 | 843886 | 3 | 169.7 | 3676.1 |
| 188 | At5g35730 | ERD1/XPR1/SYG1 | EXS (ERD1/XPR1/SYG1) family protein | 833549 | 3 | 169.9 | 334.5 |
| 189 | At1g72470 | EXO70D1 | exocyst subunit exo70 family protein D1 | 843579 | 3 | 172.7 | 110.6 |
| 190 | At5g11890 | EMB3135 | | 831062 | 3 | 173.4 | 569.3 |
| 191 | At5g04160 | transporter | Nucleotide-sugar transporter family protein | 830294 | 3 | 173.9 | 376.4 |

| | | | | | | | |
|---|---|---|---|---|---|---|---|
| 192 | At1g23490 | ARFA1A | ADP-ribosylation factor 1 | 838957 | 3 | 174 | 989.8 |
| 193 | At3g03050 | RHD7 | cellulose synthase-like D3 | 821148 | 3 | 176.1 | 195.2 |
| 194 | At5g17630 | transporter | Nucleotide/sugar transporter family protein | 831629 | 3 | 177.2 | 526.4 |
| 195 | At1g32050 | SCAMP5 | SCAMP family protein | 840097 | 1 | 177.7 | 7295.2 |
| 196 | At2g01410 | NHL | NHL domain-containing protein | 814669 | 2 | 179 | 3489.2 |
| 197 | At4g34960 | isomerase | Cyclophilin-like peptidyl-prolyl cis-trans isomerase family protein | 829648 | 3 | 179.6 | 513.5 |
| 198 | At2g26260 | 3BETAHSD/D2 | 3beta-hydroxysteroid-dehydrogenase/decarboxylase isoform 2 | 817166 | 3 | 180.3 | 1256.7 |
| 199 | At2g43130 | RABA5C | P-loop containing nucleoside triphosphate hydrolases superfamily protein | 818915 | 2 | 180.4 | 2783.8 |
| 200 | At1g17890 | GER2 | NAD(P)-binding Rossmann-fold superfamily protein | 838368 | 3 | 181.2 | 221.1 |
| 201 | At2g40730 | CTEXP | Protein kinase family protein with ARM repeat domain | 818668 | 2 | 181.4 | 150.9 |
| 202 | At4g01810 | Sec23/Sec24 transport | Sec23/Sec24 protein transport family protein | 828105 | 3 | 181.6 | 110.1 |
| 203 | At4g33530 | KUP5 | K+ uptake permease 5 | 829492 | 3 | 181.8 | 653 |
| 204 | At5g41950 | Tetratricopeptide repeat (TPR)-like | Tetratricopeptide repeat (TPR)-like superfamily protein | 834200 | 3 | 182.4 | 698.5 |
| 205 | At3g61130 | LGT1 | galacturonosyltransferase 1 | 825285 | 3 | 183.4 | 787.3 |
| 206 | At1g19440 | KCS4 | 3-ketoacyl-CoA synthase 4 | 838528 | 3 | 183.6 | 145.2 |
| 207 | At4g13940 | SAHH1 | S-adenosyl-L-homocysteine hydrolase | 827028 | 3 | 184.1 | 438.7 |
| 208 | At5g56040 | kinase | Leucine-rich receptor-like protein kinase family protein | 835702 | 3 | 184.8 | 3558.6 |
| 209 | At4g29120 | dehydrogenase | 6-phosphogluconate dehydrogenase family protein | 829033 | 3 | 185.1 | 1485.4 |
| 210 | At4g32530 | ATPase | ATPase, F0/V0 complex, subunit C protein | 829388 | 3 | 185.7 | 467 |
| 211 | At3g08550 | KOB1 | elongation defective 1 protein / ELD1 protein | 820002 | 3 | 185.9 | 108.9 |
| 212 | At1g26650 | | | 839205 | 3 | 186.2 | 2357.2 |
| 213 | At5g15470 | GAUT14 | galacturonosyltransferase 14 | 831400 | 3 | 186.7 | 100.9 |
| 214 | At1g60780 | HAP13 | Clathrin adaptor complexes medium subunit family protein | 842372 | 3 | 186.7 | 76.4 |
| 215 | At2g44520 | COX10 | cytochrome c oxidase 10 | 819059 | 3 | 186.7 | 635.2 |
| 216 | At3g25140 | QUA1 | Nucleotide-diphospho-sugar transferases superfamily protein | 822105 | 3 | 187.7 | 574.3 |
| 217 | At1g71900 | DUF803 | Protein of unknown function (DUF803) | 843521 | 3 | 187.7 | 884.1 |
| 218 | At1g07140 | SIRANBP | Pleckstrin homology (PH) domain superfamily protein | 837225 | 3 | 188.2 | 249.8 |
| 219 | At1g57620 | emp24/gp25L/p24/GOLD | emp24/gp25L/p24 family/GOLD family protein | 842138 | 3 | 189.3 | 438.4 |
| 220 | At5g44340 | TUB4 | tubulin beta chain 4 | 834459 | 3 | 190.9 | 475.9 |
| 221 | At5g47030 | ATPase | ATPase, F1 complex, delta/epsilon subunit | 834749 | 3 | 190.9 | 167.5 |
| 222 | At1g13580 | LOH3 | LAG1 longevity assurance homolog 3 | 837919 | 3 | 191.1 | 327.8 |
| 223 | At2g45540 | beige | WD-40 repeat family protein / beige-related | 819162 | 3 | 193.7 | 917.9 |

| | | | | | | | |
|---|---|---|---|---|---|---|---|
| 224 | At2g14890 | AGP9 | arabinogalactan protein 9 | 815978 | 3 | 194.9 | 1324.4 |
| 225 | At5g66680 | DGL1 | dolichyl-diphosphooligosaccharide-protein glycosyltransferase 48kDa subunit family protein | 836801 | 3 | 196.4 | 32.6 |
| 226 | At5g52560 | USP | UDP-sugar pyrophosphorylase | 835333 | 3 | 197.3 | 111.6 |
| 227 | At1g06470 | transporter | Nucleotide/sugar transporter family protein | 837159 | 3 | 197.9 | 1550.2 |
| 228 | At3g08630 | DUF3411 | Protein of unknown function (DUF3411) | 820010 | 3 | 198.6 | 791 |
| 229 | At3g07100 | SEC24A | Sec23/Sec24 protein transport family protein | 819896 | 3 | 198.6 | 867.1 |
| 230 | At3g02090 | MPPBETA | Insulinase (Peptidase family M16) protein | 821084 | 3 | 199 | 82 |
| 231 | At3g10760 | Homeodomain-like | Homeodomain-like superfamily protein | 820245 | 3 | 203.3 | 2126.9 |
| 232 | At5g27490 | Yip1 | Integral membrane Yip1 family protein | 832808 | 3 | 203.8 | 1700.9 |
| 233 | At3g12610 | DRT100 | Leucine-rich repeat (LRR) family protein | 820441 | 3 | 204.3 | 1183.9 |
| 234 | At5g08160 | PK3 | serine/threonine protein kinase 3 | 830712 | 3 | 206 | 69.8 |
| 235 | At2g39720 | RHC2A | RING-H2 finger C2A | 818556 | 3 | 206.2 | 473.1 |
| 236 | At2g20810 | LGT4 | galacturonosyltransferase 10 | 816611 | 3 | 206.7 | 586.3 |
| 237 | At1g51980 | Insulinase | Insulinase (Peptidase family M16) protein | 841627 | 3 | 207.3 | 147.6 |
| 238 | At2g30460 | transporter | Nucleotide/sugar transporter family protein | 817596 | 3 | 208.5 | 4392.2 |
| 239 | At1g15690 | FUGU5 | Inorganic H pyrophosphatase family protein | 838138 | 2 | 210.4 | 4963.8 |
| 240 | At4g35410 | Clathrin adaptor complex | Clathrin adaptor complex small chain family protein | 829694 | 3 | 211.3 | 1609.4 |
| 241 | At2g20370 | MUR3 | Exostosin family protein | 816556 | 3 | 212.3 | 121.2 |
| 242 | At3g02580 | STE1 | sterol 1 | 820679 | 3 | 212.5 | 288.7 |
| 243 | At5g05010 | clathrin adaptor complexes | clathrin adaptor complexes medium subunit family protein | 830383 | 3 | 212.6 | 174.9 |
| 244 | At2g21160 | Translocon-associated (TRAP), | Translocon-associated protein (TRAP), alpha subunit | 816651 | 3 | 212.6 | 177.3 |
| 245 | At3g05100 | transferase | S-adenosyl-L-methionine-dependent methyltransferases superfamily protein | 819672 | 3 | 212.7 | 3736.5 |
| 246 | At4g03550 | PMR4 | glucan synthase-like 5 | 825650 | 3 | 216.7 | 192.2 |
| 247 | At1g31070 | GlcNAc1pUT1 | N-acetylglucosamine-1-phosphate uridylyltransferase 1 | 839993 | 2 | 218.9 | 3512.5 |
| 248 | At3g47520 | pNAD-MDH | malate dehydrogenase | 823906 | 3 | 219.2 | 1043.4 |
| 249 | At5g06660 | Protein of unknown function DUF106 | Protein of unknown function DUF106, transmembrane | 830554 | 3 | 219.7 | 1050.4 |
| 250 | At1g09780 | iPGAM1 | Phosphoglycerate mutase, 2,3-bisphosphoglycerate-independent | 837507 | 3 | 220.7 | 309.8 |
| 251 | At5g67360 | ARA12 | Subtilase family protein | 836871 | 3 | 220.9 | 3428.6 |
| 252 | At4g12730 | FLA2 | FASCICLIN-like arabinogalactan 2 | 826885 | 3 | 221 | 454.4 |
| 253 | At2g30710 | Ypt/Rab-GAP domain of gyp1p | Ypt/Rab-GAP domain of gyp1p superfamily protein | 817622 | 2 | 221.2 | 1407.1 |
| 254 | At2g35860 | FLA16 | FASCICLIN-like arabinogalactan protein 16 precursor | 818159 | 3 | 221.2 | 1709.9 |

| | | | | | | | |
|---|---|---|---|---|---|---|---|
| 255 | At4g33580 | BCA5 | beta carbonic anhydrase 5 | 829498 | 3 | 221.5 | 5256.8 |
| 256 | At1g53240 | mMDH1 | Lactate/malate dehydrogenase family protein | 841757 | 3 | 221.8 | 2313.1 |
| 257 | At3g27230 | transferase | S-adenosyl-L-methionine-dependent methyltransferases superfamily protein | 822342 | 3 | 222.3 | 338.2 |
| 258 | At5g42080 | RSW9 | dynamin-like protein | 834213 | 2 | 222.4 | 1826.4 |
| 259 | At3g03810 | EDA30 | O-fucosyltransferase family protein | 821131 | 3 | 222.8 | 492.6 |
| 260 | At4g39390 | NST-K1 | nucleotide sugar transporter-KT 1 | 830094 | 3 | 223.7 | 179.6 |
| 261 | At4g29130 | HXK1 | hexokinase 1 | 829034 | 3 | 224.9 | 499.4 |
| 262 | At4g34450 | COP | coatomer gamma-2 subunit, putative / gamma-2 coat protein, putative / gamma-2 COP, putative | 829596 | 3 | 225.3 | 501.1 |
| 263 | At1g69330 | RING/U-box | RING/U-box superfamily protein | 843265 | 3 | 225.6 | 1190.3 |
| 264 | At1g27970 | NTF2B | nuclear transport factor 2B | 839690 | 3 | 226.8 | 771 |
| 265 | At2g29560 | ENOC | cytosolic enolase | 817505 | 3 | 227.2 | 228.3 |
| 266 | At4g26570 | CBL3 | calcineurin B-like 3 | 828764 | 3 | 228.1 | 1226.5 |
| 267 | At4g32140 | transporter | EamA-like transporter family | 829346 | 3 | 228.5 | 28.4 |
| 268 | At3g27240 | Cytochrome C1 | Cytochrome C1 family | 822343 | 3 | 228.7 | 233 |
| 269 | At5g63840 | RSW3 | Glycosyl hydrolases family 31 protein | 836504 | 3 | 229.4 | 446.6 |
| 270 | At5g11710 | ENTH/VHS | ENTH/VHS family protein | 831043 | 3 | 231.5 | 217.1 |
| 271 | At5g12850 | zinc finger | CCCH-type zinc finger protein with ARM repeat domain | 831125 | 3 | 232.2 | 157.1 |
| 272 | At1g20840 | TMT1 | tonoplast monosaccharide transporter1 | 838676 | 3 | 232.9 | 1042.3 |
| 273 | At5g67500 | VDAC2 | voltage dependent anion channel 2 | 836886 | 3 | 237 | 361.6 |
| 274 | At2g38120 | WAV5 | Transmembrane amino acid transporter family protein | 818390 | 3 | 238.4 | 4994.4 |
| 275 | At5g16440 | IPP1 | isopentenyl diphosphate isomerase 1 | 831505 | 2 | 239.3 | 1076 |
| 276 | At2g35100 | ARAD1 | Exostosin family protein | 818076 | 3 | 239.4 | 179.9 |
| 277 | At5g21105 | oxidase | Plant L-ascorbate oxidase | 832235 | 3 | 239.7 | 936.4 |
| 278 | At1g47240 | NRAMP2 | NRAMP metal ion transporter 2 | 841127 | 2 | 240 | 874.4 |
| 279 | At3g49720 | | | 824134 | 3 | 241.1 | 1557.6 |
| 280 | At2g21390 | Coatomer, | Coatomer, alpha subunit | 816678 | 3 | 241.7 | 55.8 |
| 281 | At1g14830 | DRP1C | DYNAMIN-like 1C | 838048 | 3 | 241.8 | 134.5 |
| 282 | At1g71820 | SEC6 | SEC6 | 843512 | 3 | 242.7 | 673.1 |
| 283 | At2g40800 | | | 818677 | 3 | 242.8 | 488.4 |
| 284 | At5g61790 | CNX1 | calnexin 1 | 836301 | 3 | 243.3 | 64.1 |
| 285 | At3g46820 | TOPP5 | type one serine/threonine protein phosphatase 5 | 823835 | 2 | 244.3 | 3439.2 |
| 286 | At2g25610 | ATPase | ATPase, F0/V0 complex, subunit C protein | 817101 | 3 | 244.3 | 2118.7 |
| 287 | At4g22910 | FZR2 | FIZZY-related 2 | 828390 | 3 | 246.1 | 172.7 |
| 288 | At1g60070 | Adaptor complex AP-1, | Adaptor protein complex AP-1, gamma subunit | 842301 | 3 | 247.6 | 456.8 |

| | | | | | | | |
|---|---|---|---|---|---|---|---|
| 289 | At2g44610 | RAB6A | Ras-related small GTP-binding family protein | 819069 | 3 | 249.3 | 1296.8 |
| 290 | At5g65810 | CGR3 | | 836710 | 3 | 249.7 | 1343.9 |
| 291 | At1g53500 | RHM2 | NAD-dependent epimerase/dehydratase family protein | 841785 | 3 | 250.5 | 164 |
| 292 | At2g05755 | transporter | Nodulin MtN21 /EamA-like transporter family protein | 815126 | 2 | 252.5 | 47 |
| 293 | At3g49310 | Major facilitator | Major facilitator superfamily protein | 824093 | 2 | 252.8 | 1193.4 |
| 294 | At1g76670 | transporter | Nucleotide-sugar transporter family protein | 844000 | 3 | 253.1 | 883.4 |
| 295 | At5g64090 | | | 836530 | 3 | 253.4 | 250.7 |
| 296 | At3g05420 | ACBP4 | acyl-CoA binding protein 4 | 819707 | 3 | 253.7 | 71 |
| 297 | At3g42050 | ATP synthase | vacuolar ATP synthase subunit H family protein | 823170 | 3 | 254.7 | 23.4 |
| 298 | At1g08280 | transferase | Glycosyltransferase family 29 (sialyltransferase) family protein | 837348 | 3 | 254.8 | 1993.6 |
| 299 | At4g24840 | | | 828587 | 3 | 254.8 | 302.4 |
| 300 | At5g14030 | TRAPB | translocon-associated protein beta (TRAPB) family protein | 831251 | 3 | 255.2 | 437.2 |

### Supplemental Table 2: A list of seed lines used in this study

| Line | Details | Reference |
|---|---|---|
| *7tm1-1* | SAIL_701_G12 in At5g18520 | this work |
| *7tm1-2* | Salk_134021 in At5g18520 | this work |
| *7tm2-3* | Sail_70_B11 in At3g09570 | this work |
| *7tm3-1* | Salk_062478 in At5g02630 | this work |
| *7tm4-1* | Sail_1_H08 in At5g42090 | this work |
| *7tm5-1* | Salk_044297 in At2g01070 | this work |
| *gpa1-4* | Salk_001846 | Jones et al 2003 Plant Phys 131: 1623-1627 |
| *agb1-2* | Salk_061896 | Ullah et al 2003 Plant Cell 15: 393–409 |
| *agg1-1c* | FLAG_197F06 introgressed into Col-0 | Trusov et al 2007 Plant Cell 19: 1235-1250 |
| *agg2-1* | SALK_010956 | Trusov et al 2007 Plant Cell 19: 1235-1250 |
| *agg3-1* | SALK_018024 | Chakravorty et al 2011 Plant J 67: 840–851 |
| *agg1/2/3* | *agg1-1c agg2-1 agg3-1* | Thung et al 2012 J Plant Phys 169: 542-545 |
| *prc1-1* | Q720Stop | Desnos et al *Development 122*: 683-693 |
| *prc1-1 the1-1* | *the1-1*: G110A (G37D) | Hématy et al 2007 Curr Biol 17: 922-931 |
| YFP-CESA6 | Cellulose synthase marker | Paredez et al 2006 Science 312: 1491-1495 |
| GFP-CESA3 | Cellulose synthase marker | Desprez et al 2007 PNAS 104: 15572-15577 |
| TdTom-CESA6 | Cellulose synthase marker | Sampathkumar et al Plant Phys 162: 675-688 |
| WAVE2-RFP | Late endosome marker | Geldner et al 2009 Plant J 59:169-178 |
| WAVE7-RFP | Late endosome marker | Geldner et al 2009 Plant J 59: 169-178 |
| WAVE13-RFP | TGN marker | Geldner et al 2009 Plant J 59: 169-178 |
| WAVE18-RFP | Golgi marker | Geldner et al 2009 Plant J 59: 169-178 |
| VHAa1-mRFP | TGN marker | Dettmer et al 2006 Plant Cell 18: 715-730 |
| GFP-NAG | Golgi marker | Grebe et al 2003 Curr Biol 13: 1378-1387 |
| GFP-LTI6b | Plasma memrbane marker | Cutler et al 2000 PNAS 97: 3718–3723 |
| PIN2-GFP | Auxin efflux carrier marker | Xu, & Scheres 2005 Plant Cell 17: 525-536 |
| ratioSecGFP | Secretion marker | Samalova et al 2006 Traffic 7: 1701-23 |

**Supplemental Table 3: A list of primers used in this study**

| Primer | Sequence | Use |
|---|---|---|
| SP241 | GACGGTGAAGAGAAGAACAACACG | Genotyping: *7tm1-1* |
| SP242 | TTACGGGTCCTGGATGAAACTATG | Genotyping: *7tm1-1* |
| SP196 | CGGCTAATGAGTAATGGGCAATG | Genotyping: *7tm1-2* |
| SP197 | TGCTTTCCTCGGTTTCTGGAG | Genotyping: *7tm1-2* |
| At3g09570.P1 | atgggatagtcgtggactcg | Genotyping: *7tm2-3* |
| At3g09570.P2 | GGATCGGATCAGACGAAGAA | Genotyping: *7tm2-3* |
| At5g02630.P7 | TGTTCTAATCGGAAACGGTTG | Genotyping: *7tm3-1* |
| At5g02630.P8 | ATTCATGTCACGCCTGTTACC | Genotyping: *7tm3-1* |
| At5g42090.P1 | GTGGGCTTCTTTCTCTGCAC | Genotyping: *7tm4-1* |
| At5g42090.P2 | CTAACTCGCTGGCAACAACA | Genotyping: *7tm4-1* |
| SP624 | AGATTTGACAACCCCAAATCC | Genotyping: *7tm5-1* |
| SP625 | AGAACAAGAGCGACTCTGCAG | Genotyping: *7tm5-1* |
| At2g26300.P3 | GAAATGAGAGGAGGAAATGGA | Genotyping: *gpa1-4* |
| At2g26300.P4 | acacggtcattcagttttcg | Genotyping: *gpa1-4* |
| At4g34460.P1 | CAGGGGATATGTTTCGTGCT | Genotyping: *agb1-2* |
| At4g34460.P2 | GCACTTCCATCTGCTGACAA | Genotyping: *agb1-2* |
| AGG1 38F | TGTGGTTTACGAGCAGGAGGAGTC | Genotyping: *agg1-1c* |
| AGG1 R | TGTCTAGCGAATCAAAGTATTAAGCATCTG | Genotyping: *agg1-1c* |
| AGG2 26F | ATGGAAGCGGGTAGCTCCAATTCGTC | Genotyping: *agg2-1* |
| AGG2 281Rs | TCAAAGAATGGAGCAGCCACATC | Genotyping: *agg2-1* |
| AGG3 71F | TGCTGGTGGAGTGAGTTCATCGTC | Genotyping: *agg3-1* |
| AGG3-1R | TTCATAGTACACATGCACGAACACATTAGG | Genotyping: *agg3-1* |
| SALK LBa1 | TGGTTCACGTAGTGGGCCATCG | Genotyping: Salk Insert |
| LB3 | TAGCATCTGAATTTCATAACCAATCTCGATACAC | Genotyping: Sail Insert |
| INRA Tag5 | CTACAAATTGCCTTTTCTTATCGAC | Genotyping: Flag Insert |
| AT3G18780.RT.P1 | TGGATTTGCAGGAGATGATG | RT-PCR: *ACT2* |
| AT3G18780.RT.P2 | GGCCTTTGGGTTAAGAGGAG | RT-PCR: *ACT2* |
| AT5G18520.RT.P3 | TGGTAAAGCTGCGAGGAACT | RT-PCR: *7TM1* |
| AT5G18520.RT.P4 | GCAGCAGCTTCTTCTTCATCA | RT-PCR: *7TM1* |
| AT2G01070.RT.P3 | TGCCTTCTGGGATGTTATCG | RT-PCR: *7TM5* |
| AT2G01070.RT.P4 | TCCGCATTCTTCTCTTGCTT | RT-PCR: *7TM5* |
| At5g18520CF | ATGAAGAAGAAGCTGCTGCTTTGGCCCTTAGGGACGAAGAGTTTGAGC TTGGAGGTGGAGGTGGAGCT | Cloning: *7TM1-YFP & 7TM1-RFP* |
| At5g18520CR | AAGAATAAATCCTTGCTTACCAAGAAAAATCCTCCATTGTATGCTTTTCA GGCCCCAGCGGCCGCAGCAGCACC | Cloning: *7TM1-YFP & 7TM1-RFP* |
| At5g18520CTF | AGTACCAGTGGGTGAGCTTTG | Cloning: *7TM1-YFP & 7TM1-RFP* |
| At5g18520CTR | ATTTGCTACAGCCCCAAAATC | Cloning: *7TM1-YFP & 7TM1-RFP* |
| At5g18520delRB | GAAATGTTTCCTGTGCAGAAATCTTGCCGCCATTGTAACAACTCCTGAA Gttaccaatgcttaatcagtg | Cloning: *7TM1-YFP & 7TM1-RFP* |
| At5g18520delTRB | ACCACCTTCTAAATAAGTAGC | Cloning: *7TM1-YFP & 7TM1-RFP* |
| At5g18520delLB | CTGGATTGATCTCCCTGAAGATGAAGAAGTTAGAAGTAATCTAAAGAA GGccctcttgggttatcaagagg | Cloning: *7TM1-YFP & 7TM1-RFP* |
| At5g18520delTLB | AACGGAGCTTTGTTGCACCGG | Cloning: *7TM1-YFP & 7TM1-RFP* |
| SP1333 7TM5 for pEZR(K)LNC (EcoRI) LP | GCCGGAATTCATGGATTTGGGAAAACTGA | Cloning: *7TM5-CFP* |
| SP1334 7TM5 for pEZR(K)LNC (BamHI) RP | GCGCGGATCCGGCTCTCTTTTATCTTCTTCAACATC | Cloning: *7TM5-CFP* |
| GPA1 Fwd | CTTTGTACAAAAAAGCAGGCTCCGCGGCCGCATGGGCTTACTCTGCAG TAGAAGTCG | Cloning: *GPA1-GFP & GFP-GPA1* |
| GPA1 Rev-Stop | CTATGGGTCGGCGCGCCTAAAAGGCCAGCCTCCAGTAAATTTCTC | Cloning: *GPA1-GFP* |
| GPA1 Rev+Stop | CAAGAAAGCTGGGTCGGCGCGCCTCATAAAAGGCCAGCCTCCAGTAAA TTTCTC | Cloning: *GFP-GPA1* |

| | | |
|---|---|---|
| SP4557 | **GCGGCCGC**CCGGTTTAGCTGGTTCACTTT | Cloning: *mCherry-AGB1*<br>*(AGB1 promoter & 5` UTR + NotI site)* |
| SP4558 | **GCGGCCGC**TCCGGGATCAGACTTAGGCTT | Cloning: *mCherry-AGB1*<br>*(AGB1 promoter & 5` UTR + NotI site)* |
| SP4553 | CACCATGTCTGTCTCCGAGCTCAA | Cloning: *AGB1-mCherry (AGB1 –stop)* |
| SP4554 | AATCACTCTCCTGTGTCCTCC | Cloning: *AGB1-mCherry (AGB1 –stop)* |
| SP4559 | **GGCGCGCC**ATGGTGAGCAAGGGCGAGGA | Cloning: *AGB1-mCherry*<br>*(mCherry+stop + AscI site)* |
| SP4560 | **GGCGCGCCGGTACC**CTACTTGTACAGCTCGTCCAT | Cloning: *AGB1-mCherry*<br>*(mCherry+stop + AscI & KpnI sites)* |
| SP4563 | **GGTACC**AGAAGATTTAACGAAAGTAGGA | Cloning: *AGB1-mCherry (AGB1 3` UTR + KpnI site)* |
| SP4564 | **GGTACC**TTTCTCGCCAAGTTGGGAATC | Cloning: *AGB1-mCherry (AGB1 3` UTR + KpnI site)* |
| SP4557 | **GCGGCCGC**CCGGTTTAGCTGGTTCACTTT | Cloning: *mCherry-AGB1*<br>*(AGB1 promoter & 5` UTR + NotI site)* |
| SP4558 | **GCGGCCGC**TCCGGGATCAGACTTAGGCTT | Cloning: *mCherry-AGB1*<br>*(AGB1 promoter & 5` UTR + NotI site)* |
| SP4555 | CACCATGGTGAGCAAGGGCGAGGA | Cloning: *mCherry-AGB1, mCherry-AGG1, mCherry-AGG2, mCherry-AGG3*<br>*(mCherry –stop)* |
| SP4556 | CTTGTACAGCTCGTCCATGCC | Cloning: *mCherry-AGB1, mCherry-AGG1, mCherry-AGG2, mCherry-AGG3*<br>*(mCherry –stop)* |
| SP4561 | **GGCGCGCC**ATGTCTGTCTCCGAGCTCAAA | Cloning: *mCherry-AGB1*<br>*(AGB1 and 3` UTR + AscI site)* |
| SP4562 | **GGCGCGCC**TTTCTCGCCAAGTTGGGAATC | Cloning: *mCherry-AGB1*<br>*(AGB1 and 3` UTR + AscI site)* |
| SP4565 | **GCGGCCGC**TGATCCACATGTAGATAGGG | Cloning: *mCherry-AGG1*<br>*(AGG1 promoter & 5` UTR + NotI site)* |
| SP4566 | **GCGGCCGC**CCTCTCGCTAGCAGGTCGCAG | Cloning: *mCherry-AGG1*<br>*(AGG1 promoter & 5` UTR + NotI site)* |
| SP4567 | **GGCGCGCC**ATGCGAGAGGAAACTGTGGTT | Cloning: *mCherry-AGG1*<br>*(AGG1 and 3` UTR + AscI site)* |
| SP4651 | **GGCGCGCC**TATCATGGAGGGTGGATGTGGCCAAAAGTT | Cloning: *mCherry-AGG1*<br>*(AGG1 and 3` UTR + AscI site)* |
| SP4569 | **GCGGCCGC**GGAGTCTTTAGATCGAGTTTT | Cloning: *mCherry-AGG2*<br>*(AGG2 promoter & 5` UTR + NotI site)* |
| SP4570 | **GCGGCCGC**CAAAATTTCTCGAATTCAACC | Cloning: *mCherry-AGG2*<br>*(AGG2 promoter & 5` UTR + NotI site)* |
| SP4571 | **GGCGCGCC**ATGGAAGCGGGTAGCTCCAA | Cloning: *mCherry-AGG2*<br>*(AGG2 and 3` UTR + AscI site)* |
| SP4572 | **GGCGCGCC**GACACCATTGTAGCCTCTCTC | Cloning: *mCherry-AGG2*<br>*(AGG2 and 3` UTR + AscI site)* |
| SP4573 | **GCGGCCGC**GATGTTGAAATTGTTGTAATA | Cloning: *mCherry-AGG3*<br>*(AGG3 promoter & 5` UTR + NotI site)* |
| SP4574 | **GCGGCCGC**TTTCGGAGAGAAATGGTTTGG | Cloning: *mCherry-AGG3*<br>*(AGG3 promoter & 5` UTR + NotI site)* |
| SP4575 | **GGCGCGCC**ATGTCTGCTCCTTCTGGCGGT | Cloning: *mCherry-AGG3*<br>*(AGG3 and 3` UTR + AscI site)* |
| SP4576 | **GGCGCGCC**ATGGTCCATATATAACTCGTA | Cloning: *mCherry-AGG3*<br>*(AGG3 and 3` UTR + AscI site)* |
| N/A | See Gookin & Assmann 2014 Plant J<br>https://doi.org/10.1111/tpj.12639 | Cloning: GPA1 for BiFC |

| | | |
|---|---|---|
| 7TM1 RsrII 24F | ATGCGGTCCGCGAAAATGCCCCTCTCCGTC | Cloning: 7TM1 for BiFC |
| 7TM1 AatII 1300R | TCGCGACGTCAGCTCAAACTCTTCGTCCCTAAG | Cloning: 7TM1 for BiFC |
| 7TM5 RsrII 25F | ATGCGGTCCGATTTGGGAAAACTGATCCAGC | Cloning: 7TM5 for BiFC |
| 7TM5sv1 AatII 1458R | TCCGGACGTCTCTCTTTTATCTTCTTCAACATCTTCTTCC | Cloning: 7TM5 for BiFC |
| PIP2pD10/12MCS3F | CTGGAGCTTCAGGAATGGGG ATGGCAAAGGATGTGGAAGCCGTTC | Cloning: PIP2 for BiFC |
| PIP2pD10/12MCS3R | CACGTGACTACGTAGGGAC TTAGACGTTGGCAGCACTTCTGAATGATC | Cloning: PIP2 for BiFC |
